# *flashfmZoom*: a tool for joint fine-mapping and exploration of GWAS results in the UK Biobank

**DOI:** 10.1101/2023.03.21.533700

**Authors:** Feng Zhou, Colin Starr, Adam S Butterworth, Jennifer L Asimit

## Abstract

**Summary:** *flashfmZoom* is an all-in-one tool for analysis and interactive visualisation of potential causal genetic variants that underlie associations with quantitative traits from the UK Biobank. It offers a user-friendly interface and guides users in the selection of pleiotropic regions among subsets of 134 quantitative traits, such as cardiometabolic, hematologic, and respiratory traits. Users may then run single-trait fine-mapping, allowing for multiple causal variants, and leverage information between the traits using multi-trait fine-mapping to improve resolution. A series of interactive plots and downloadable tables are generated within *flashfmZoom* to identify potential causal variants that are shared or distinct between the traits; it also lists relevant literature for the traits and/or variants. Besides exploring traits that are well-known to be related, *flashfmZoom* encourages interactive exploration for the joint analysis of traits that may not often be considered together. This may reveal common aetiological pathways between traits related to different disorders.

**Availability and Implementation:** *flashfmZoom* is an interactive open-source R shiny app software available online directly at https://mrc-bsu.shinyapps.io/flashfmZoomOnline/, with source code in GitHub under the MIT license https://github.com/fz-cambridge/flashfmZoom and at https://doi.org/10.5281/zenodo.7756205.

## Introduction

Construction of an accurate shortlist of potential causal variants that underlie genetic associations with diseases and traits assists in deciphering the findings of genome-wide association studies (GWAS). Statistical fine-mapping aims to reduce this shortlist of genetic variants for follow-up in downstream functional validation experiments, which may lead to new biological insights for diseases or even new therapeutic targets (Hutchinson et al., 2020). Mult-trait fine-mapping that leverages information between traits can improve precision beyond fine-mapping each trait independently, since biologically related traits often have shared causal variants (Hernandez et al., 2021). *Flashfm* jointly fine-maps multiple quantitative traits, allowing for multiple causal variants without the restriction of shared causal variants (Hernandez et al., 2021).

Comparing fine-mapping results between traits and methods is simplified by visualisations such as regional association plots and Venn diagrams. This motivated the development of *flashfm-ivis* (Zhou et al., 2022), which enables users to interact with a series of visualisations of their single-trait and multi-trait fine-mapping results. Unlike most bioinformatics plotting tools, no programming knowledge is required for the user-friendly interface of *flashfm-ivis*.

Biobank data resources offer genetic data and many phenotypes from a large sample in a single population, with the UK Biobank (UKBB) being one of the largest (Bycroft et al., 2018). Building on our accessibility principle, we have developed *flashfmZoom*, which combines various online data sources to explore genetic association signals among user-selected subsets of 134 UKBB traits and fine-maps signals in multiple traits, independently (*FINEMAP* (Benner et al., 2016) or *JAM* (Newcombe et al., 2016)) and jointly (*flashfm* (Hernandez et al., 2021)) (**Supplementary Materials S.1**); traits are measured in 361,194 unrelated European ancestry participants from UKBB (**Supplementary Materials S.2**). Our collection of traits includes cardiometabolic, hematologic, respiratory, and anthropometric, among others (**Supplementary Table S1**). Traits across different classes may share common aetiological pathways and analysis with *flashfm* could reveal shared causal variants. In turn, this could open new avenues of research between traits that may not have been considered previously.

We considered comparisons of credible sets from single-trait and multi-trait fine-mapping for several trait pairs and regions, indicating the percentage reduction in 99% credible set (CS99) size by *flashfm* (**Table 1**). We highlight the trait results for “*standing height*” and “*forced vital capacity*” (*FVC*; a lung function test that measures the total amount of air exhaled forcefully) in a region containing the protein-coding gene *HGFAC*; the correlation between these traits is 0.434. *FVC* is expected to be influenced by height, which determines body size, and height is a highly polygenic trait (Guo et al., 2018). For *standing height*, multi-trait fine-mapping using *flashfm* gave a CS99 reduction over single-trait fine-mapping using *JAM* of 62.5%, down to nine. Of these variants, *flashfm* strongly favours rs13108218 with highest marginal posterior probability (MPP) of a variant being causal for height (MPP 0.899). By contrast, *JAM* favoured rs59950280 (MPP 0.991), which has r^2^=0.47 with rs13108218. For *FVC, flashfm* also strongly favoured rs13108218 (MPP 0.999), with a substantial increase in support beyond *JAM* (MPP 0.681).

**Table 1:**
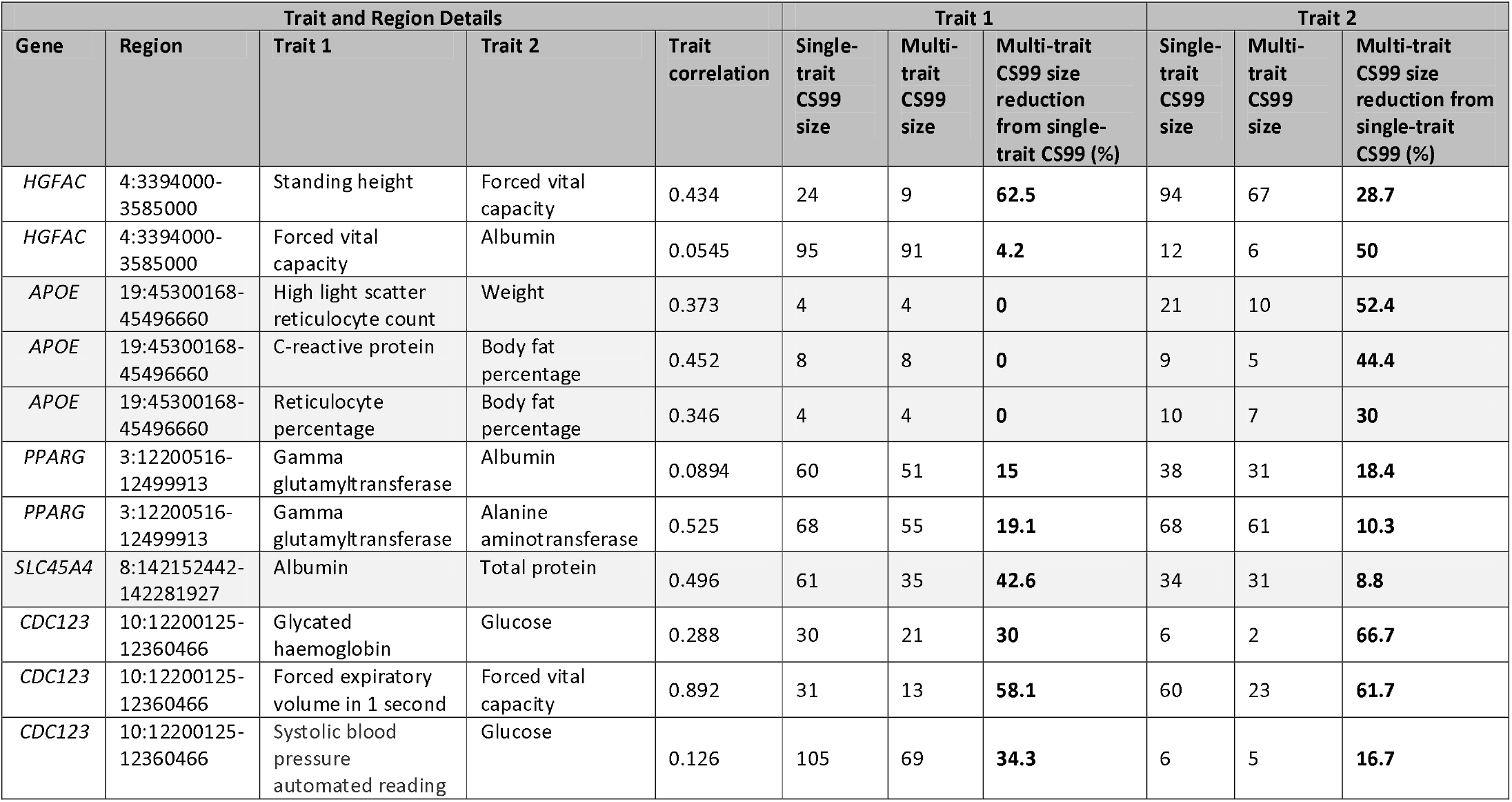
Results from single-trait and multi-trait fine-mapping for 5 exemplar regions. For each region, signals from a pair of traits from UK Biobank were fine-mapped by single and multi-trait fine-mapping. Comparisons of the 99% credible sets (CS99) from both methods are given for traits 1 and 2. For each trait, the reduction (%) in credible set size of multi-trait fine-mapping compared to single-trait fine-mapping is also provided. In these analyses all variants with MAF>0.001 were included.

| Trait and Region Details |  |  |  |  | Trait 1 |  |  | Trait 2 |  |  |
| --- | --- | --- | --- | --- | --- | --- | --- | --- | --- | --- |
| Gene | Region | Trait 1 | Trait 2 | Trait correlation | Single-trait CS99 size | Multi-trait CS99 size | Multi-trait CS99 size reduction from single-trait CS99 (%) | Single-trait CS99 size | Multi-trait CS99 size | Multi-trait CS99 size reduction from single-trait CS99 (%) |
| *HGFAC* | 4:3394000-3585000 | Standing height | Forced vital capacity | 0.434 | 24 | 9 | **62.5** | 94 | 67 | **28.7** |
| *HGFAC* | 4:3394000-3585000 | Forced vital capacity | Albumin | 0.0545 | 95 | 91 | **4.2** | 12 | 6 | **50** |
| *APOE* | 19:45300168-45496660 | High light scatter reticulocyte count | Weight | 0.373 | 4 | 4 | **0** | 21 | 10 | **52.4** |
| *APOE* | 19:45300168-45496660 | C-reactive protein | Body fat percentage | 0.452 | 8 | 8 | **0** | 9 | 5 | **44.4** |
| *APOE* | 19:45300168-45496660 | Reticulocyte percentage | Body fat percentage | 0.346 | 4 | 4 | **0** | 10 | 7 | **30** |
| *PPARG* | 3:12200516-12499913 | Gamma glutamyltransferase | Albumin | 0.0894 | 60 | 51 | **15** | 38 | 31 | **18.4** |
| *PPARG* | 3:12200516-12499913 | Gamma glutamyltransferase | Alanine aminotransferase | 0.525 | 68 | 55 | **19.1** | 68 | 61 | **10.3** |
| *SLC45A4* | 8:142152442-142281927 | Albumin | Total protein | 0.496 | 61 | 35 | **42.6** | 34 | 31 | **8.8** |
| *CDC123* | 10:12200125-12360466 | Glycated haemoglobin | Glucose | 0.288 | 30 | 21 | **30** | 6 | 2 | **66.7** |
| *CDC123* | 10:12200125-12360466 | Forced expiratory volume in 1 second | Forced vital capacity | 0.892 | 31 | 13 | **58.1** | 60 | 23 | **61.7** |
| *CDC123* | 10:12200125-12360466 | Systolic blood pressure automated reading | Glucose | 0.126 | 105 | 69 | **34.3** | 6 | 5 | **16.7** |

*FlashfmZoom* has a simple interface and offers a “one-stop shop” for:

1. Interactive visualisations of UKBB genetic associations with multiple traits, together with functional annotation, gene locations, and published genetic associations (**Supplementary Figure S1, Supplementary Table S2**);
2. Conducting a PheWAS (phenome-wide association study) at a single variant, rapidly identifying all traits that are associated with the variant, and visualising the correlations between these traits;
3. Choosing a suitable fine-mapping region based on GWAS summary statistics and exploring an index of previous existing studies/publications in this region;
4. Running *JAM* (Newcombe et al., 2016) or *FINEMAP* (Benner et al., 2016) with *flashfm* (Hernandez et al., 2021) directly from the website (**Supplementary Materials S.1**);
5. Interactive visualisations of fine-mapping results, comparing results by method and trait (**Supplementary Table S3**).

An overview is given in **Figure 1** (with complementary figures in **Supplementary Figure S2-S6**); see **Supplementary Materials S.4** for instructions and other features (e.g. dynamic “Need Help” features, flexible tabs, interactive widgets, and downloadable tables).

**Figure 1:**
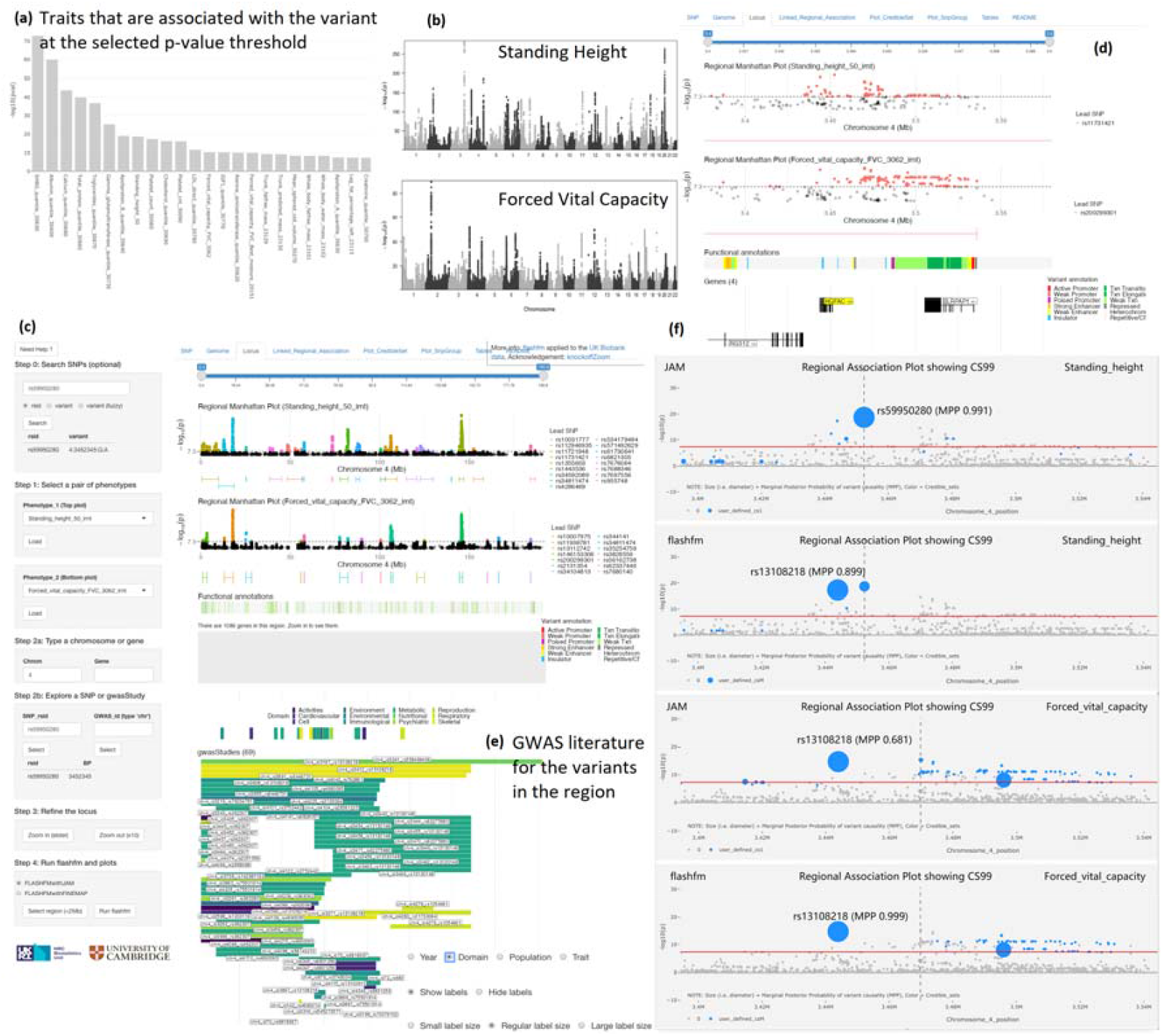
Example FlashfmZoom displays for exploring standing height and forced vital capacity in HGFAC. **(a)** Phewas results are shown for a single variant (rs59950280; chr4:3452345) as a bar plot of the -log10(p-value) of association with traits that have p<5e-8 (other thresholds may be chosen, i.e. 1e-6); the traits are sorted by the pvalue and the SNP is specified in the Step_0 control widget. **(b)** Manhattan plots for standing height and forced vital capacity, the two selected phenotypes in Step_1, show many genome-wide significant signals at multiple chromosomes, including chromosome 4. **(c)** For the pair of traits, regional association plots are shown for chromosome 4, as specified using the control widget in Step_2a. **(d)** The further refined region, centred around rs59950280 (chr4:3452345), includes details such as gene locations and functional annotation; the variant to form the region around is specified in the Step_2b widget. **(e)** Display of the previously published GWAS studies (labelled by lead SNP and other related information such as chr_id, PMID, traits, etc. (Supplementary Figure S7 is a larger, clearer version) of variants within the refined region. Here, studies are grouped by domain, though other options for grouping are publication year, population(s) in the study, and traits. The horizontal bar for each publication indicates the region defined in that study and the vertical bars above the publications indicate locations of lead SNPs from the studies. The bar length for each study is as defined in the published region, the rsid is the lead SNP in each publication. **(f)** Interactive regional association plots integrated with fine-mapping results. The diameter of each point is proportional to the marginal posterior probability (MPP) that the SNP is causal for the trait under either single (cs1) or multi-trait fine-mapping (csM). Points belonging to the 99% credible set are coloured in blue; users may specify different levels for the credible set and different colours for the points. The dashed line indicates the location of the lead SNP. For standing height, its lead SNP, rs59950280, has the highest MPP under single-trait fine-mapping; under flashfm the MPP for rs13108218 increases and becomes the largest. For FVC, rs13108218 is not the lead SNP, but it has the highest MPP under both JAM and flashfm.

## Implementation

Similar in concept to *flashfm-ivis* (Zhou et al., 2022), *flashfmZoom* is built in R and its web-based version does not require users to have any programming skills.

1. Data inputs and connected online sources Users do not need to input any data, as all on-line datasets are pre-loaded (“Data Availability”, **Supplementary Material S.2**). This includes GWAS summary statistics of 134 UKBB quantitative traits (http://www.nealelab.is/uk-biobank/); variants with MAF < 0.001 are excluded, being flagged as low confidence variants, and genomic positions are based on genome assembly GRCh37/hg19.
2. PheWAS and trait correlations To help guide the selection of traits for fine-mapping, users may input a genetic variant (rsID or genomic position) and view the associated UKBB traits and their p-value (Figure 1a); available p-value thresholds are 1×10^−5^, 1×10^−6^ or 5×10^−8^. As the highest gains in using multi-trait fine-mapping over single-trait fine-mapping tend to be when the traits have low to moderate correlation, a trait correlation heatmap of all associated traits is displayed (**Supplementary Figure S2**).
3. Manhattan plots, regional association plots, and region selection Based on PheWAS information or previous knowledge, users can select a pair of traits from the drop-down lists to display their Manhattan plots (**Figure 1b**). Regional association plots for the selected traits can be displayed by selecting a chromosome or gene (**Figure 1c, Figure 1d**).
4. Regional annotations and previously published signals If the region displayed in the regional association plots is small enough (e.g., 500kb), additional details will be shown for the region: (i) locations of genes; (ii) variant annotations (ENCODE Project Consortium, 2012); (iii) publications (PMID) on variants associated with any trait (Watanabe et al., 2019). Publications may be viewed according to publication date, trait, medical domain (e.g., Metabolic), or ancestral populations (**Figure 1e**). Regions may also be defined by selecting a SNP as a region midpoint (± 250kb) or selecting a region defined in a published GWAS.
5. Refinement and selection of locus The selected region for fine-mapping must be smaller than 2Mb, for computational efficiency. Once selected, LD is approximated using the pre-loaded 1000 Genomes phase 3 European super-population data (The 1000 Genomes Project Consortium, 2015). The page then displays information on the number of variants with MAF >0.001 and MAF >0.005, in the selected region, and the minimum p-value for each of the traits. The region can then be further refined. Both traits should have a variant (not necessarily the same variant) with p-value < 1×10^−5^ in the region to proceed with fine-mapping.
6. Single and multi-trait fine-mapping *Flashfm* (Hernandez et al., 2021) multi-trait fine-mapping is run with single-trait fine-mapping by *FINEMAP* (Benner et al., 2016) or *JAM* (Newcombe et al., 2016). A series of interactive plots are available for the fine-mapping results: (i) linked regional association plots with probabilities that each variant is causal for each trait and colours that indicate exchangeable variants (**Supplementary Figure S3**); (ii) regional association plots indicating credible sets (default 99%, though adjustable) for each trait (**Figure 1f and Supplementary Figure S4**); (iii) Venn diagrams to show shared and distinct potential causal variants within the CS99 of each trait (**Supplementary Figure S5**); (iv) Sankey diagrams to show the variants that belong to each SNP group under single-trait and multi-trait fine-mapping – this also shows the MPP for each variant/group being causal (**Supplementary Figure S6**).
7. Download results and tables of outputs For transparency and further investigation of the results, all information and outputs that are used and displayed in the plots can be easily downloaded in the final tab. This includes the full table of GWAS association results, SNP groups and credible sets for both traits from different fine-mapping methods (**Supplementary Material S.4**).

## Conclusion

We have created a user-friendly interactive web tool *flashfmZoom*, that makes use of several sources of data to enable users to explore and fine-map genetic associations in multiple traits within UKBB - all without any programming knowledge. It enables joint fine-mapping of traits that may not often be considered together and helps with comparing results between traits and methods (single-trait and multi-trait fine-mapping) through interactive visualisations. Summary tables of credible sets that partition variants by inclusion for one or both traits are also available for download for further follow-up. *FlashfmZoom* also compiles a list of relevant publications for a selected trait or genetic variant, to raise awareness of existing studies and make connections with UKBB results. We believe that *flashfmZoom* will assist researchers in making new links between traits and contribute to unravelling the complex underlying mechanisms of various diseases.

## Acknowledgements

The authors would like to thank the authors of KnockoffZoom (Sesia et al., 2020, 2021), echolocatoR (Schilder et al., 2020) and LocusZoom (Pruim et al., 2010; Boughton et al., 2020), as their clear software documentation and coding implementations were a key ingredient in the development of *flashfmZoom*.

## Data Availability

The datasets can be found at the *flashfmZoom* GitHub repository (https://github.com/fz-cambridge/flashfmZoom). Other external data sources: (1) UK Biobank GWAS summary statistics: http://www.nealelab.is/uk-biobank and https://github.com/Nealelab/UK_Biobank_GWAS; (2) 1000 Genomes European super-population data to approximate LD matrices: https://ctg.cncr.nl/software/MAGMA/ref_data/; (3) Trait correlation between traits in the UK Biobank: https://ukbb-rg.hail.is or the associated data in Supplementary Materials (Bulik-Sullivan et al., 2015) https://www.ncbi.nlm.nih.gov/pmc/articles/PMC4797329/#sd2; (4) GWAS publications from GWASATLAS https://atlas.ctglab.nl (Watanabe et al., 2019); (5) Gene positions file ncbiRefSeq.txt https://www.ncbi.nlm.nih.gov/refseq/ (O’Leary et al., 2016); (6) Variant annotations (ENCODE Project Consortium, 2012) file wgEncodeBroadHmmGm12878HMM.txt from http://hgdownload.cse.ucsc.edu/goldenpath/hg19/encodeDCC/wgEncodeBroadHmm/

## Supplementary Material

See online appendix and README at https://github.com/fz-cambridge/flashfmZoom which also includes YouTube video demonstrations of the tool.

