## Supplementary Materials for "*flashfmZoom*: a tool for joint fine-mapping and exploration of GWAS results in the UK Biobank"

**Table of contents (Labels as cited in the main paper)**


- **[Page 2] Supplementary Material S.1:** FLASHFMwithJAM and FLASHFMwithFINEMAP
- **[Page 3] Supplementary Material S.2:** Details on the UK Biobank and other data sources
- **[Page 4] Supplementary Table S1:** Selected 134 quantitative traits from the UK Biobank
- **[Page 5] Supplementary Material S.3:** Complementary figures to the main paper Figure 1
    - **[Page 5] Supplementary Figure S1:** Package flowchart
    - **[Page 6] Supplementary Figure S2:** PheWAS and trait correlation
    - **[Page 7] Supplementary Figure S3:** Linked regional association plots
    - **[Page 8] Supplementary Figure S4:** Interactive regional association
    - **[Page 9] Supplementary Figure S5:** Venn diagrams
    - **[Page 10] Supplementary Figure S6:** Sankey diagrams
    - **[Page 11] Supplementary Figure S7:** GWAS literature for variants in this region
- **[Page 12] Supplementary Material S.4:** FlashfmZoom details and instructions
    - **[Page 12] Supplementary Table S2:** Control widgets of flashfmZoom interface
    - **[Page 13] Supplementary Table S3:** Main tabs of flashfmZoom interface
        - Step_0: Search SNPs (optional)
        - Step_1: Select a pair of phenotypes or traits
        - Step_2a: Type a chromosome or gene
        - Step_2b: Explore a SNP or GWAS
        - Step_3: Refine the locus
        - Step_4: Run flashfm and plots
        - Download results and tables
        - Useful features: “Need Help?” button
    - **[Page 40] Supplementary Table S4:** Modebar icons in plots
    - **[Page 41] Terminology in the plots**
- **[Page 41] Reference (as included in Supplementary Material)**


**Availability and Implementation:** *flashfmZoom* is an interactive open-source R shiny app software available online directly at https://mrc-bsu.shinyapps.io/flashfmZoomOnline/, with source code in GitHub under the MIT license, https://github.com/fz-cambridge/flashfmZoom and at https://doi.org/10.5281/zenodo.7756205 . Additional features can be downloaded locally as standalone R libraries to encourage reuse.

**Contact:**

### S.1 FLASHFMwithJAM and FLASHFMwithFINEMAP

The flashfm R package (https://jennasimit.github.io/flashfm/ ; Hernandez et al. 2021) includes two wrapper functions to run flashfm multi-trait fine-mapping together with single-trait fine-mapping using either JAM (Newcombe et al., 2016) or FINEMAP (Benner et al., 2016). Each of these methods allows for multiple causal variants and the model posterior probabilities from single-trait fine-mapping are used as input into flashfm.

Both functions require GWAS summary statistics from each trait, the LD matrix for the region, effect allele frequencies, and the trait correlation matrix. Only variants that appear in all traits and in the LD matrix will be carried forward for analysis, and this variant filtering is carried out within flashfm. Prior to running either wrapper function, the effective sample size should be calculated for each trait using all variants with MAF>0.01; this is done using the flashfm::Neff function with the effect allele frequencies (or MAF) and the standard error of the GWAS effect estimates (beta.std.err). Trait correlation (or genetic correlation) may be estimated using LD score regression as was done for UK Biobank (Bulik-Sullivan et al., 2015).

Output from flashfm (both wrapper functions) includes:

- **PP** (posterior probability) for each multi-SNP model: the model PP for each configuration of variants being joint causal variants for the trait (i.e., multiple causal variants). The PPs sum to 1.
- **MPP** (marginal posterior probability of variant causality): the posterior probability that the SNP is included in any causal model; it is the sum over all model PPs that include the SNP. MPP is also known as PIP (Posterior Inclusion Probability).
- **SNP groups:** SNPs belonging to the same group can be viewed as exchangeable. SNPs with MPP>0.001 are assigned to the same group if they are strongly correlated (pairwise $r^2$>0.6) and rarely appear in a model together (marginal posterior probability that a pair of SNPs are both included in a model < 0.01). Each SNP group based on flashfm tends to contain a subset of variants from groups based on single-trait fine-mapping.
- **PPg (PP based on SNP groups):** for a particular group-based model containing multiple SNP groups (e.g., groups A and B), PPg is the sum of PPs for all models composed of exactly one SNP from group A and one SNP from group B. As the SNPs in each group are exchangeable, the PPg represents the probability that the correct causal model involves a SNP from group A and a SNP from group B.
- **MPPg (MPP based on SNP groups):** MPPg for SNP group A is the sum of MPPs for all SNPs that belong to group A; it is the sum over all model PPs that include a SNP from group A. This represents the probability of at least one of the SNPs in SNP group A being a causal variant for the traits.

### S.2 Details on the UK Biobank and other data sources

(1) GWAS summary statistics for the UK Biobank: http://www.nealelab.is/uk-biobank and https://github.com/Nealelab/UK_Biobank_GWAS
All analysis details are given at https://github.com/Nealelab/UK_Biobank_GWAS , but we provide some of the details here.
These are the v3 GWAS summary statistics for round 2 results of the UK Biobank [released 1st August 2018], based on 361,194 unrelated samples. The Neale group ran linear regression using Hail with covariates: 1st 20 PCs + sex + age + age^2 + sex*age + sex*age2; we use their results for both sexes. Only variants with MAF > 0.001 are deemed to pass quality control and we also use this as a default threshold for our analyses.

(2) LD data from European super-population of 1000 Genomes, phase 3: https://ctg.cncr.nl/software/MAGMA/ref_data/
We allow a maximum region size of 2MB for computational efficiency.

(3) Trait correlation between traits in the UK Biobank: https://ukbb-rg.hail.is or the associated data in Supplementary Materials (Bulik-Sullivan et al., 2015)
https://www.ncbi.nlm.nih.gov/pmc/articles/PMC4797329/#sd2
Trait correlation between all pairs of traits have been calculated by Bulik-Sullivan et al., 2015 using LD score regression.

(4) GWAS publications from GWASATLAS: https://atlas.ctglab.nl
This database includes all publications on GWAS up to 2019 and is searchable on trait and genetic variant.

(5) Gene locations and Annotation files: ncbiRefSeq.txt https://www.ncbi.nlm.nih.gov/refseq/ and wgEncodeBroadHmmGm12878HMM.txt from
http://hgdownload.cse.ucsc.edu/goldenpath/hg19/encodeDCC/wgEncodeBroadHmm/
All positions are based on genome assembly GRCh37/hg19.

**Table S1**: **Lists of all phenotype codes and their descriptions for the selected 134 quantitative traits from the UK Biobank that are included in the current version of flashfmZoom**. The detailed excel file (with more information and links to the original data source) can be downloaded from the GitHub. Note: users can use this table (either trait code or name) to search a pair of phenotypes in Step_1 of flashfmZoom.

| # of traits | Phenotype Code | Phenotype Description | # of traits | Phenotype Code | Phenotype Description |
|---|---|---|---|---|---|
| 1 | 30620_irnt | Alanine aminotransferase (quantile) | 68 | 23107_irnt | Impedance of leg (right) |
| 2 | 30600_irnt | Albumin (quantile) | 69 | 23106_irnt | Impedance of whole body |
| 3 | 30610_irnt | Alkaline phosphatase (quantile) | 70 | 5262_irnt | Intra-ocular pressure, corneal-compensated (left) |
| 4 | 4100_irnt | Ankle spacing width (left) | 71 | 5254_irnt | Intra-ocular pressure, corneal-compensated (right) |
| 5 | 4119_irnt | Ankle spacing width (right) | 72 | 5263_irnt | Intra-ocular pressure, Goldmann-correlated (left) |
| 6 | 30630_irnt | Apoliprotein A (quantile) | 73 | 5255_irnt | Intra-ocular pressure, Goldmann-correlated (right) |
| 7 | 30640_irnt | Apoliprotein B (quantile) | 74 | 30780_irnt | LDL direct (quantile) |
| 8 | 23124_irnt | Arm fat mass (left) | 75 | 23116_irnt | Leg fat mass (left) |
| 9 | 23120_irnt | Arm fat mass (right) | 76 | 23112_irnt | Leg fat mass (right) |
| 10 | 23123_irnt | Arm fat percentage (left) | 77 | 23115_irnt | Leg fat percentage (left) |
| 11 | 23119_irnt | Arm fat percentage (right) | 78 | 23111_irnt | Leg fat percentage (right) |
| 12 | 23125_irnt | Arm fat-free mass (left) | 79 | 23117_irnt | Leg fat-free mass (left) |
| 13 | 23121_irnt | Arm fat-free mass (right) | 80 | 23113_irnt | Leg fat-free mass (right) |
| 14 | 23126_irnt | Arm predicted mass (left) | 81 | 23118_irnt | Leg predicted mass (left) |
| 15 | 23122_irnt | Arm predicted mass (right) | 82 | 23114_irnt | Leg predicted mass (right) |
| 16 | 30650_irnt | Aspartate aminotransferase (quantile) | 83 | 30790_irnt | Lipoprotein A (quantile) |
| 17 | 23105_irnt | Basal metabolic rate | 84 | 30120_irnt | Lymphocyte count |
| 18 | 30220_irnt | Basophill percentage | 85 | 30180_irnt | Lymphocyte percentage |
| 19 | 20022_irnt | Birth weight | 86 | 6033_irnt | Maximum heart rate during fitness test |
| 20 | 23099_irnt | Body fat percentage | 87 | 6032_irnt | Maximum workload during fitness test |
| 21 | 21001_irnt | Body mass index (BMI) | 88 | 30050_irnt | Mean corpuscular haemoglobin |
| 22 | 23104_irnt | Body mass index (BMI) | 89 | 30060_irnt | Mean corpuscular haemoglobin concentration |
| 23 | 30710_irnt | C-reactive protein (quantile) | 90 | 30040_irnt | Mean corpuscular volume |
| 24 | 30680_irnt | Calcium (quantile) | 91 | 30100_irnt | Mean platelet (thrombocyte) volume |
| 25 | 30690_irnt | Cholesterol (quantile) | 92 | 30260_irnt | Mean reticulocyte volume |
| 26 | 5264_irnt | Corneal hysteresis (left) | 93 | 30270_irnt | Mean sphered cell volume |
| 27 | 5256_irnt | Corneal hysteresis (right) | 94 | 30500_irnt | Microalbumin in urine |
| 28 | 5265_irnt | Corneal resistance factor (left) | 95 | 30130_irnt | Monocyte count |
| 29 | 5257_irnt | Corneal resistance factor (right) | 96 | 30190_irnt | Monocyte percentage |
| 30 | 30510_irnt | Creatinine (enzymatic) in urine | 97 | 20127_irnt | Neuroticism score |
| 31 | 30700_irnt | Creatinine (quantile) | 98 | 30140_irnt | Neutrophill count |
| 32 | 30720_irnt | Cystatin C (quantile) | 99 | 30200_irnt | Neutrophill percentage |
| 33 | 4079_irnt | Diastolic blood pressure, automated reading | 100 | 30800_irnt | Oestradiol (quantile) |
| 34 | 94_irnt | Diastolic blood pressure, manual reading | 101 | 3064_irnt | Peak expiratory flow (PEF) |
| 35 | 30660_irnt | Direct bilirubin (quantile) | 102 | 30810_irnt | Phosphate (quantile) |
| 36 | 5983_irnt | ECG, heart rate | 103 | 30080_irnt | Platelet count |
| 37 | 5984_irnt | ECG, load | 104 | 30090_irnt | Platelet crit |
| 38 | 5986_irnt | ECG, phase time | 105 | 30110_irnt | Platelet distribution width |
| 39 | 30210_irnt | Eosinophill percentage | 106 | 30520_irnt | Potassium in urine |
| 40 | 30897_irnt | Estimated sample dilution factor (quantile) | 107 | 4194_irnt | Pulse rate |
| 41 | 3063_irnt | Forced expiratory volume in 1-second (FEV1) | 108 | 30010_irnt | Red blood cell (erythrocyte) count |
| 42 | 20150_irnt | Forced expiratory volume in 1-second (FEV1), Best measure | 109 | 30070_irnt | Red blood cell (erythrocyte) distribution width |
| 43 | 20153_irnt | Forced expiratory volume in 1-second (FEV1), predicted | 110 | 30250_irnt | Reticulocyte count |
| 44 | 3062_irnt | Forced vital capacity (FVC) | 111 | 30240_irnt | Reticulocyte percentage |
| 45 | 20151_irnt | Forced vital capacity (FVC), Best measure | 112 | 30820_irnt | Rheumatoid factor (quantile) |
| 46 | 30730_irnt | Gamma glutamyltransferase (quantile) | 113 | 30830_irnt | SHBG (quantile) |
| 47 | 30740_irnt | Glucose (quantile) | 114 | 30530_irnt | Sodium in urine |
| 48 | 30750_irnt | Glycated haemoglobin (quantile) | 115 | 50_irnt | Standing height |
| 49 | 30030_irnt | Haematocrit percentage | 116 | 4080_irnt | Systolic blood pressure, automated reading |
| 50 | 30020_irnt | Haemoglobin concentration | 117 | 93_irnt | Systolic blood pressure, manual reading |
| 51 | 47_irnt | Hand grip strength (right) | 118 | 30840_irnt | Total bilirubin (quantile) |
| 52 | 30760_irnt | HDL cholesterol (quantile) | 119 | 30860_irnt | Total protein (quantile) |
| 53 | 4105_irnt | Heel bone mineral density (BMD) (left) | 120 | 30870_irnt | Triglycerides (quantile) |
| 54 | 4124_irnt | Heel bone mineral density (BMD) (right) | 121 | 23128_irnt | Trunk fat mass |
| 55 | 4106_irnt | Heel bone mineral density (BMD) T-score, automated (left) | 122 | 23127_irnt | Trunk fat percentage |
| 56 | 4125_irnt | Heel bone mineral density (BMD) T-score, automated (right) | 123 | 23129_irnt | Trunk fat-free mass |
| 57 | 4101_irnt | Heel broadband ultrasound attenuation (left) | 124 | 23130_irnt | Trunk predicted mass |
| 58 | 4120_irnt | Heel broadband ultrasound attenuation (right) | 125 | 30880_irnt | Urate (quantile) |
| 59 | 4104_irnt | Heel quantitative ultrasound index (QUI), direct entry (left) | 126 | 30670_irnt | Urea (quantile) |
| 60 | 30300_irnt | High light scatter reticulocyte count | 127 | 30890_irnt | Vitamin D (quantile) |
| 61 | 30290_irnt | High light scatter reticulocyte percentage | 128 | 48_irnt | Waist circumference |
| 62 | 49_irnt | Hip circumference | 129 | 21002_irnt | Weight |
| 63 | 30770_irnt | IGF-1 (quantile) | 130 | 23098_irnt | Weight |
| 64 | 30280_irnt | Immature reticulocyte fraction | 131 | 30000_irnt | White blood cell (leukocyte) count |
| 65 | 23110_irnt | Impedance of arm (left) | 132 | 23100_irnt | Whole body fat mass |
| 66 | 23109_irnt | Impedance of arm (right) | 133 | 23101_irnt | Whole body fat-free mass |
| 67 | 23108_irnt | Impedance of leg (left) | 134 | 23102_irnt | Whole body water mass |

### S.3 Complementary Figures

#### Figure S1. Flowchart of FlashfmZoom

*In this flowchart, the unshaded cells represent datasets/methods and their manipulations in the background of this flashfmZoom package, the light grey coloured cells represent main steps of users' implementation, and the dark grey coloured cells represent main outputs and plots. Examples of detailed plots and outputs are explained further in **Figure_1**. There are 4 main parts in this flowchart: **(1)** This relates to Step_0 and Step_1, which focus on users' selection of phenotypes, optionally using a PheWAS to identify traits associated with a SNP. In the background, UK Biobank GWAS summary statistics are uploaded and refined based on MAF>0.001 and pval thresholds (e.g. pval < 1e-6 or 5e-8), as well as the selected 134 traits (for more details, see online supplementary materials). Manhattan plots for all 134 traits are also saved/pre-loaded in the background. **(2)** This is related to Step_2a and Step_2b (see purple-dashed box), when users explore and refine a region based on chromosome number, gene name, SNP rsid or gwasStudy PMID. Gene files and a previous existing GWAS publication list are pre-loaded in the background. **(3)** This is Step_3, which focuses on preparing to run the flashfm package (see green-dashed box). Users can refine sub-regions further, to run different flashfm methods efficiently. Both fine-mapping methods and external datasets (i.e. both LD matrix and trait correlation for users' selected traits in the UK Biobank) are connected together. **(4)** Outputs and results from these single- and multi-trait fine-mapping methods can be downloaded as common .csv/.RData/.xlsx files and tables for users to do further analyses in their local machines, and flashfmZoom offers many linked interactive plots for users to explore the outputs visually (see blue-dashed box). The plots are mainly focused on regional association results, SNP groups and credible sets.*

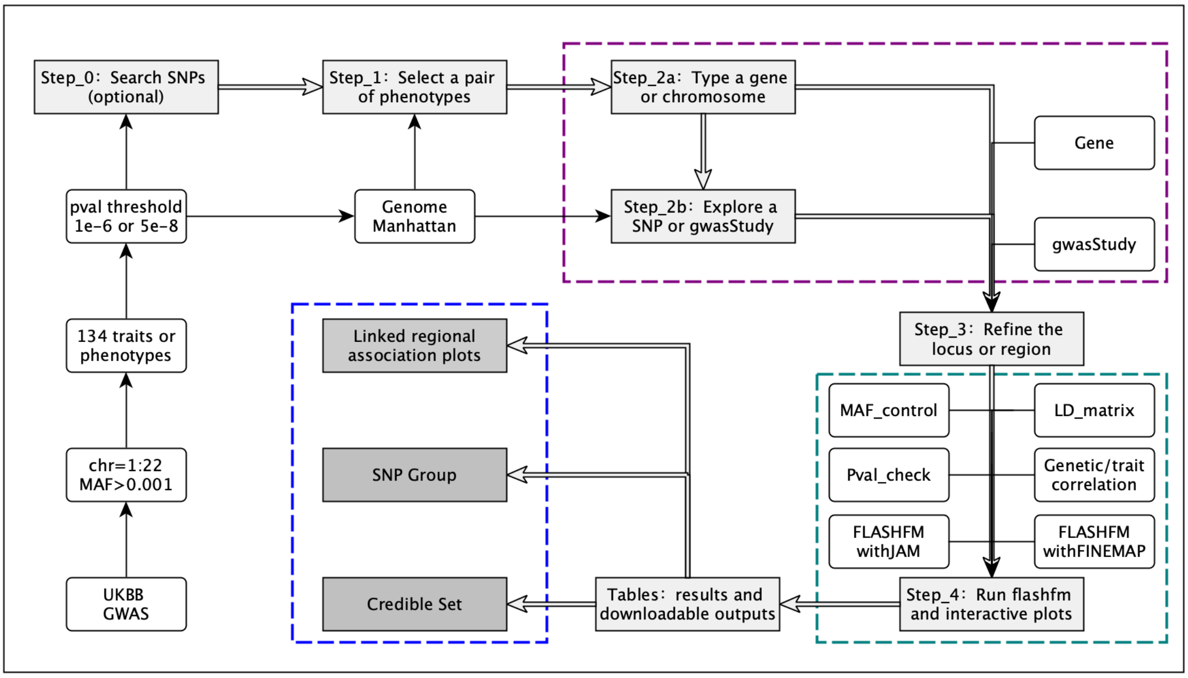

### Figure S2: PheWAS at a variant in *HGFAC* and correlation among associated traits

*To help guide the selection of traits for fine-mapping, users may input a genetic variant and view the associated UKBB traits and their p-value. The default p-value threshold is 1E-5, and more stringent thresholds are also available (1E-6 or 5E-8). Users can search a genetic variant using its rsID (e.g. rs59950280) or genomic position (e.g. 4:3452345:G:A) by an exact or partial match ("fuzzy search"; e.g. 4:3452345); if more than one SNP is found, the most appropriate SNP may be selected from the list. As the highest gains in using multi-trait over single-trait fine-mapping are when the traits have low to moderate correlation, the trait correlations (with a heatmap) are provided for all traits associated with the variant. Correlations are given as a matrix that allows users to sort correlations within a given trait, with the option to download. (Complementary to Figure 1)*

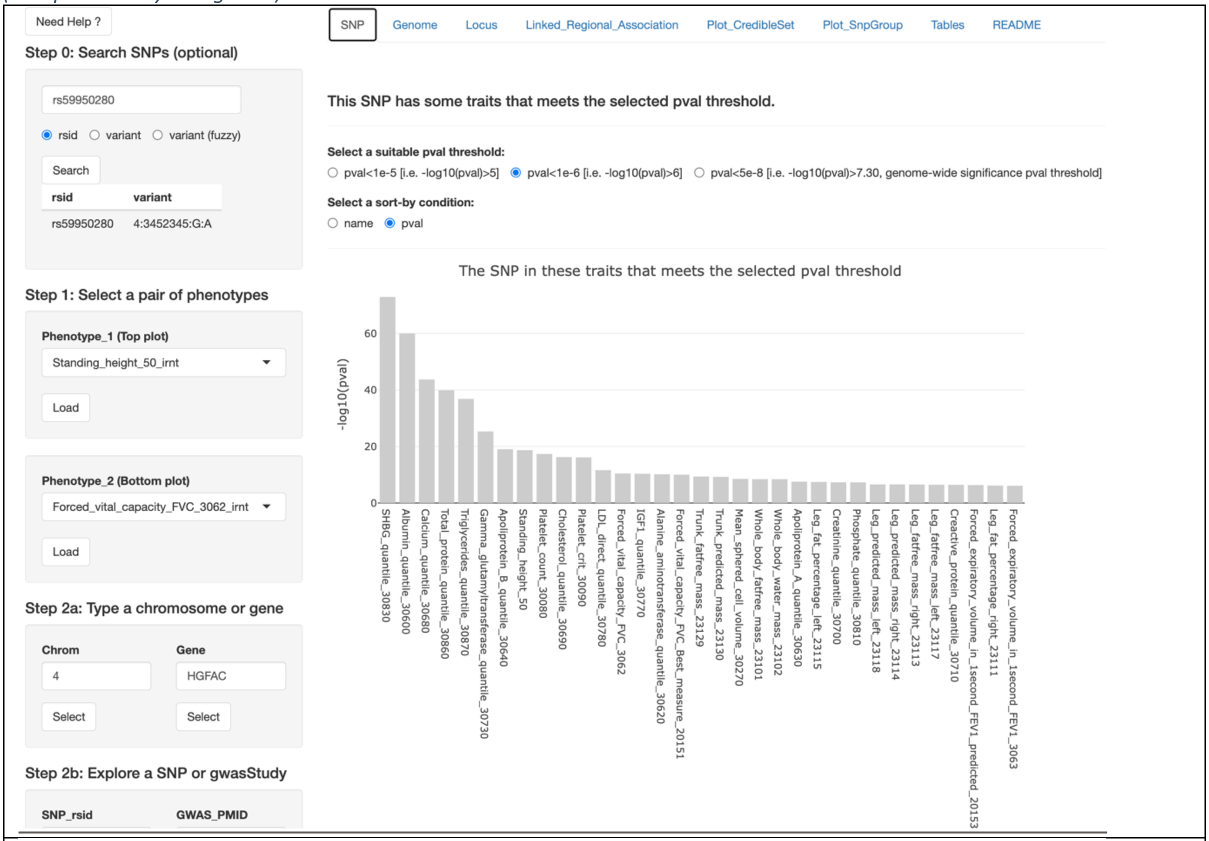


#### Trait correlation table

| | trait_20151 | trait_20153 | trait_23101 | trait_23102 | trait_30080 | trait_30090 | trait_30270 | trait_3062 | trait_3063 | trait_50 | trait_30600 | trait_30700 | trait_30620 | trait_30630 | trait_30640 |
|---|---|---|---|---|---|---|---|---|---|---|---|---|---|---|---|
| trait_20151 | 1 | 0.457 | 0.174 | 0.171 | -0.065 | -0.073 | 0.017 | 0.95 | 0.868 | 0.462 | 0.052 | 0.054 | -0.113 | 0.081 | -0.02 |
| trait_20153 | 0.457 | 1 | 0.528 | 0.522 | -0.093 | -0.092 | -0.007 | 0.45 | 0.373 | 0.969 | -0.007 | 0.06 | -0.032 | -0.009 | -0.027 |
| trait_23101 | 0.174 | 0.528 | 1 | 0.998 | -0.058 | -0.043 | -0.108 | 0.162 | 0.174 | 0.539 | -0.132 | 0.147 | 0.162 | -0.206 | 0 |
| trait_23102 | 0.171 | 0.522 | 0.998 | 1 | -0.057 | -0.042 | -0.109 | 0.16 | 0.172 | 0.534 | -0.134 | 0.148 | 0.162 | -0.208 | 0 |
| trait_30080 | -0.065 | -0.093 | -0.058 | -0.057 | 1 | 0.863 | -0.059 | -0.065 | -0.066 | -0.1 | -0.02 | -0.053 | -0.006 | 0 | 0.102 |
| trait_30090 | -0.073 | -0.092 | -0.043 | -0.042 | 0.863 | 1 | -0.067 | -0.072 | -0.07 | -0.097 | -0.036 | -0.041 | 0 | -0.038 | 0.083 |
| trait_30270 | 0.017 | -0.007 | -0.108 | -0.109 | -0.059 | -0.067 | 1 | 0.019 | -0.013 | -0.008 | 0 | -0.052 | -0.05 | 0.194 | -0.042 |
| trait_3062 | 0.95 | 0.45 | 0.162 | 0.16 | -0.065 | -0.072 | 0.019 | 1 | 0.893 | 0.434 | 0.055 | 0.051 | -0.108 | 0.084 | -0.018 |
| trait_3063 | 0.868 | 0.373 | 0.174 | 0.172 | -0.066 | -0.07 | -0.013 | 0.893 | 1 | 0.373 | 0.068 | 0.058 | -0.068 | 0.059 | -0.003 |
| trait_50 | 0.462 | 0.969 | 0.539 | 0.534 | -0.1 | -0.097 | -0.008 | 0.434 | 0.373 | 1 | -0.008 | 0.058 | -0.039 | -0.007 | -0.032 |
| trait_30600 | 0.052 | -0.007 | -0.132 | -0.134 | -0.02 | -0.036 | 0 | 0.055 | 0.068 | -0.008 | 1 | -0.016 | 0.098 | 0.186 | 0.101 |
| trait_30700 | 0.054 | 0.06 | 0.147 | 0.148 | -0.053 | -0.041 | -0.052 | 0.051 | 0.058 | 0.058 | -0.016 | 1 | -0.009 | -0.053 | -0.022 |
| trait_30620 | -0.113 | -0.032 | 0.162 | 0.162 | -0.006 | 0 | -0.05 | -0.108 | -0.068 | -0.039 | 0.098 | -0.009 | 1 | -0.096 | 0.101 |
| trait_30630 | 0.081 | -0.009 | -0.206 | -0.208 | 0 | -0.038 | 0.194 | 0.084 | 0.059 | -0.007 | 0.186 | -0.053 | -0.096 | 1 | -0.047 |
| trait_30640 | -0.02 | -0.027 | 0 | 0 | 0.102 | 0.083 | -0.042 | -0.018 | -0.003 | -0.032 | 0.101 | -0.022 | 0.101 | -0.047 | 1 |
| trait_30680 | 0.003 | -0.024 | -0.071 | -0.072 | 0.093 | 0.111 | -0.011 | 0.008 | 0.019 | -0.024 | 0.522 | 0.065 | 0.123 | 0.148 | 0.139 |
| trait_30690 | 0.022 | -0.02 | -0.076 | -0.077 | 0.097 | 0.065 | 0.03 | 0.023 | 0.029 | -0.023 | 0.154 | -0.039 | 0.046 | 0.279 | 0.89 |
| trait_30710 | -0.223 | -0.069 | 0.195 | 0.196 | 0.154 | 0.163 | -0.071 | -0.212 | -0.194 | -0.077 | -0.199 | 0.034 | 0.169 | -0.195 | 0.12 |
| trait_30730 | -0.152 | -0.062 | 0.108 | 0.108 | 0.074 | 0.069 | 0.013 | -0.143 | -0.118 | -0.072 | 0.089 | -0.023 | 0.526 | -0.005 | 0.134 |
| trait_30770 | 0.116 | 0.056 | -0.019 | -0.019 | -0.019 | -0.025 | -0.043 | 0.112 | 0.105 | 0.062 | 0.127 | 0.068 | -0.057 | 0.001 | -0.006 |
| trait_30780 | 0.012 | -0.014 | -0.03 | -0.03 | 0.1 | 0.074 | -0.019 | 0.012 | 0.024 | -0.017 | 0.118 | -0.024 | 0.056 | 0.062 | 0.958 |
| trait_30810 | 0.002 | -0.014 | -0.054 | -0.055 | 0.076 | 0.079 | 0.016 | 0.004 | -0.012 | -0.01 | 0.114 | 0.038 | -0.009 | 0.06 | 0.001 |
| trait_30830 | 0.124 | 0.023 | -0.26 | -0.261 | -0.075 | -0.073 | 0.164 | 0.117 | 0.075 | 0.027 | -0.036 | -0.077 | -0.238 | 0.258 | -0.093 |
| trait_30860 | -0.047 | -0.039 | -0.074 | -0.075 | 0.116 | 0.117 | 0.003 | -0.042 | -0.02 | -0.042 | 0.497 | 0.017 | 0.133 | 0.104 | 0.163 |
| trait_30870 | -0.133 | -0.046 | 0.179 | 0.18 | 0.09 | 0.099 | -0.169 | -0.128 | -0.099 | -0.051 | 0.013 | 0.032 | 0.262 | -0.27 | 0.345 |

### Figure S3: Linked Regional Association Plots for standing height and forced vital capacity in *HGFAC*

*Linked regional association plots with probability that each variant is causal for each trait and colours that indicate exchangeable variants. The diameter of each point is proportional to the marginal posterior probability (MPP) that the SNP is causal for the trait under either single or multi-trait fine-mapping. Information of individual SNPs are displayed in the box. The first and last portion of the LD matrix of the region is also provided. (Complementary to Figure 1)*

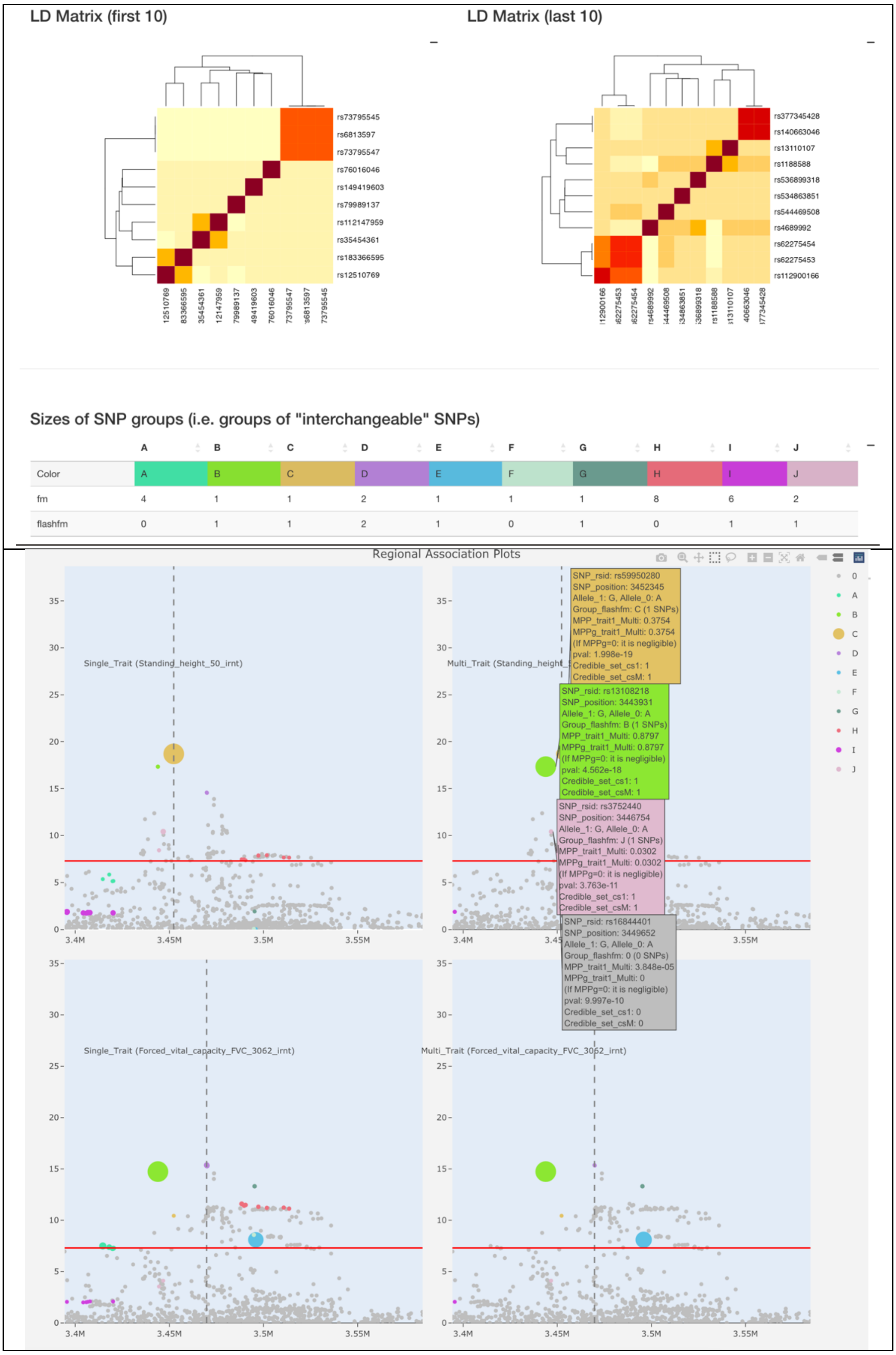

### Figure S4: Interactive Regional Association plots with 99% credible sets for standing height and forced vital capacity in *HGFAC*

*Regional association plots integrated with fine-mapping results. The diameter of each point is proportional to the marginal posterior probability (MPP) that the SNP is causal for the trait under either single or multi-trait fine-mapping. Coloured points indicate variants that belong to credible sets (default 99% and adjustable by slider bar) for each trait (i.e. Trait_1 in the top two panels is Standing Height and Trait_2 in the bottom two panels is Forced Vital Capacity). Information for individual SNPs are displayed in the box. (Complementary to Figure 1)*

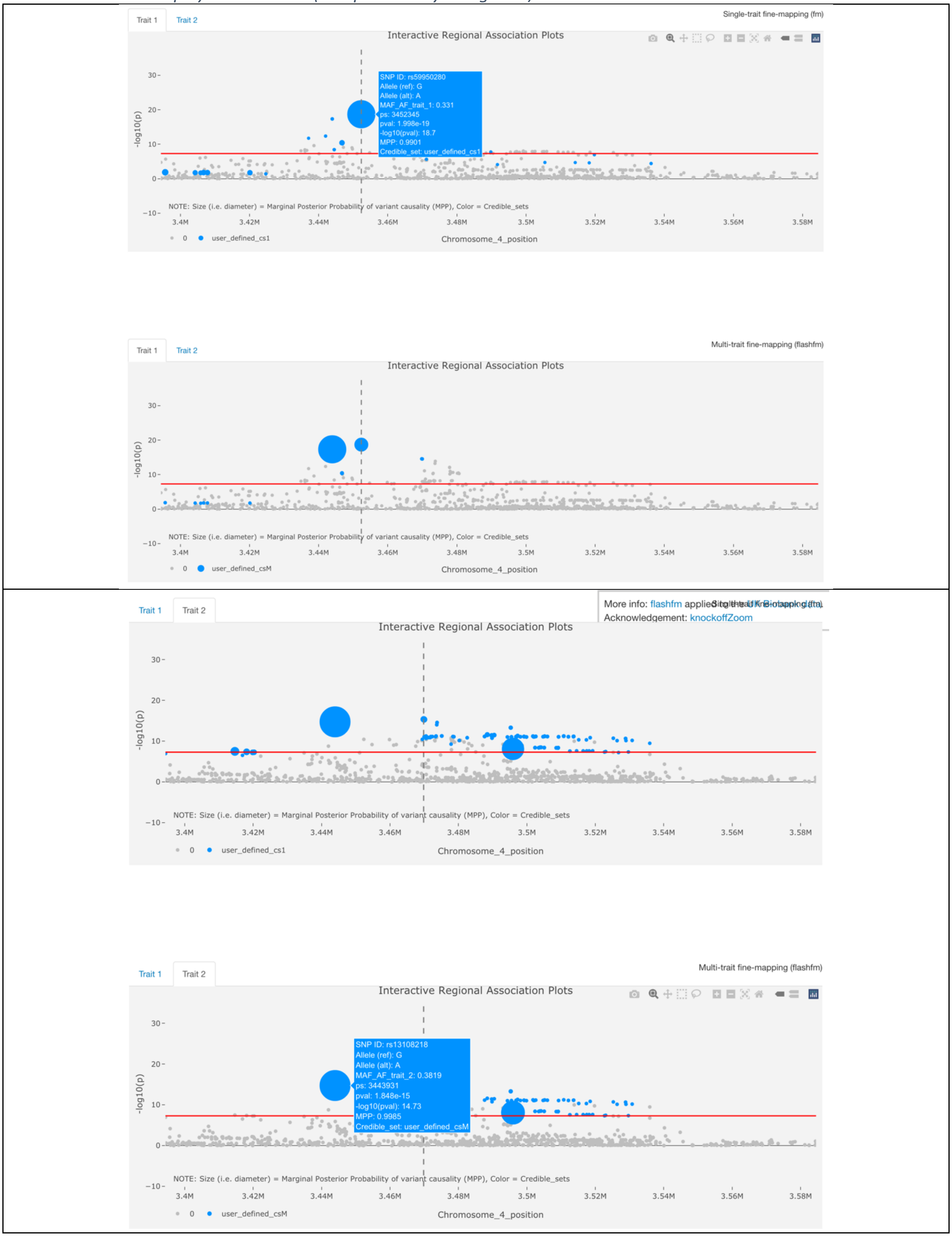

### Figure S5: Venn Diagrams of credible sets for standing height and forced vital capacity in *HGFAC*

*Venn diagrams to show shared and distinct potential causal variants within the 99% credible sets (i.e., cs_1 is based on single-trait and cs_M is based on multi-trait fine-mapping) of each trait (i.e., The first trait is Standing Height as named "trait_50" and the second trait is Forced Vital Capacity as named "trait_3062"). The details of SNPs that are shared or distinct among the traits can be downloaded as a csv/excel table. (Complementary to Figure 1)*

Venn diagrams: Credible Sets | Interactive Venn diagram 1: cs1 | Interactive Venn diagram 2: csM

Venn diagrams (single-trait cs_1) with "eulerr" R Package based on "ellipse" shape

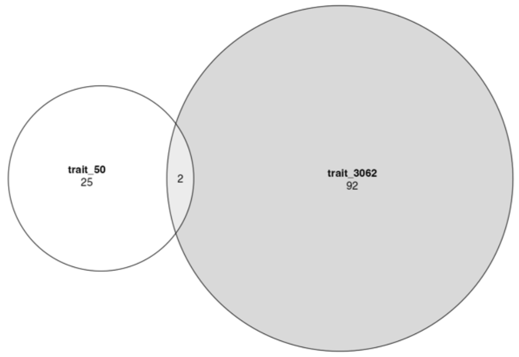


Venn diagrams (multi-trait cs_M) with "eulerr" R Package based on "ellipse" shape

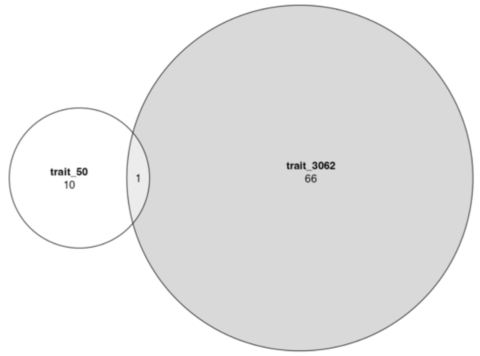


Venn diagrams: Credible Sets | Interactive Venn diagram 1: cs1 | Interactive Venn diagram 2: csM

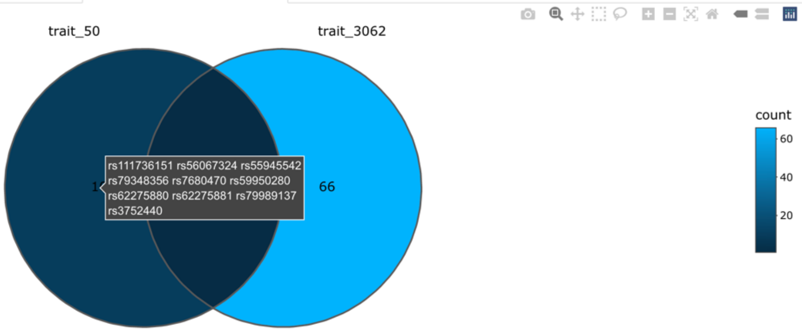


#### Table of trait credible sets from multi-trait fine-mapping (cs_M) and their intersections

| | component | id | name | count | item |
|---|---|---|---|---|---|
| 1 | region | 1 | trait_50 | 10 | rs111736151,rs56067324,rs55945542,rs79348356,rs7680470,rs59950280,rs62275880,rs62275881,rs79989137,rs3752440 |
| 2 | region | 2 | trait_3062 | 66 | rs10014466,rs10028098,rs1049574,rs112795540,rs13109504,rs13112054,rs13115353,rs13116668,rs13117941,rs13125213,rs131[illegible] |
| 3 | region | 12 | trait_50..trait_3062 | 1 | rs13108218 |

CSV Excel

NOTE: The column "id" indicates the intersection of different traits, e.g. "1" is "trait_1", "12" is an intersection of "trait_1" and "trait_2".

### Figure S6: Sankey Diagrams for standing height and forced vital capacity in *HGFAC*

*Sankey diagrams to show the variants that belong to each SNP group under single-trait and multi-trait fine-mapping – this also shows the amount of support for each variant/group being causal. Both the combined values of two selected traits and values of individual traits (Top panel is Trait_1 Standing Height and bottom panel is Trait_2 Forced Vital Capacity) can be displayed.*

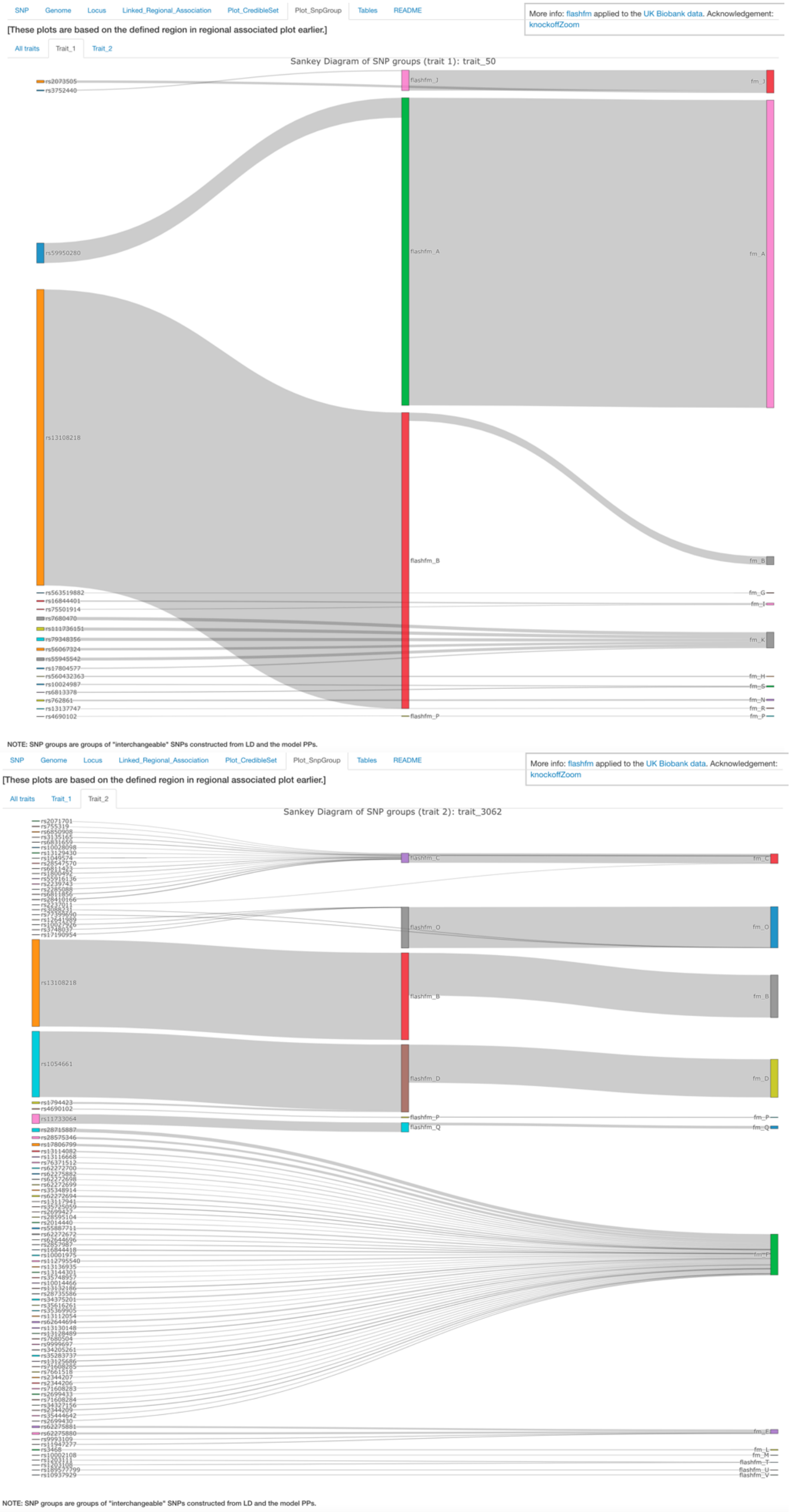

### Figure S7: GWAS literature for variants in this region

*Display of the previously published GWAS studies (labelled by PMID and lead SNP) of variants within the refined region. Here, studies are grouped by domain, though other options for grouping are publication year, population(s) in the study, and traits. The horizontal bar for each publication indicates the region defined in that study and the vertical bars above the publications indicate locations of lead SNPs from the studies. The bar length for each study is as defined in the published region, the rsid is the lead SNP in each publication. (Complementary to Figure 1)*

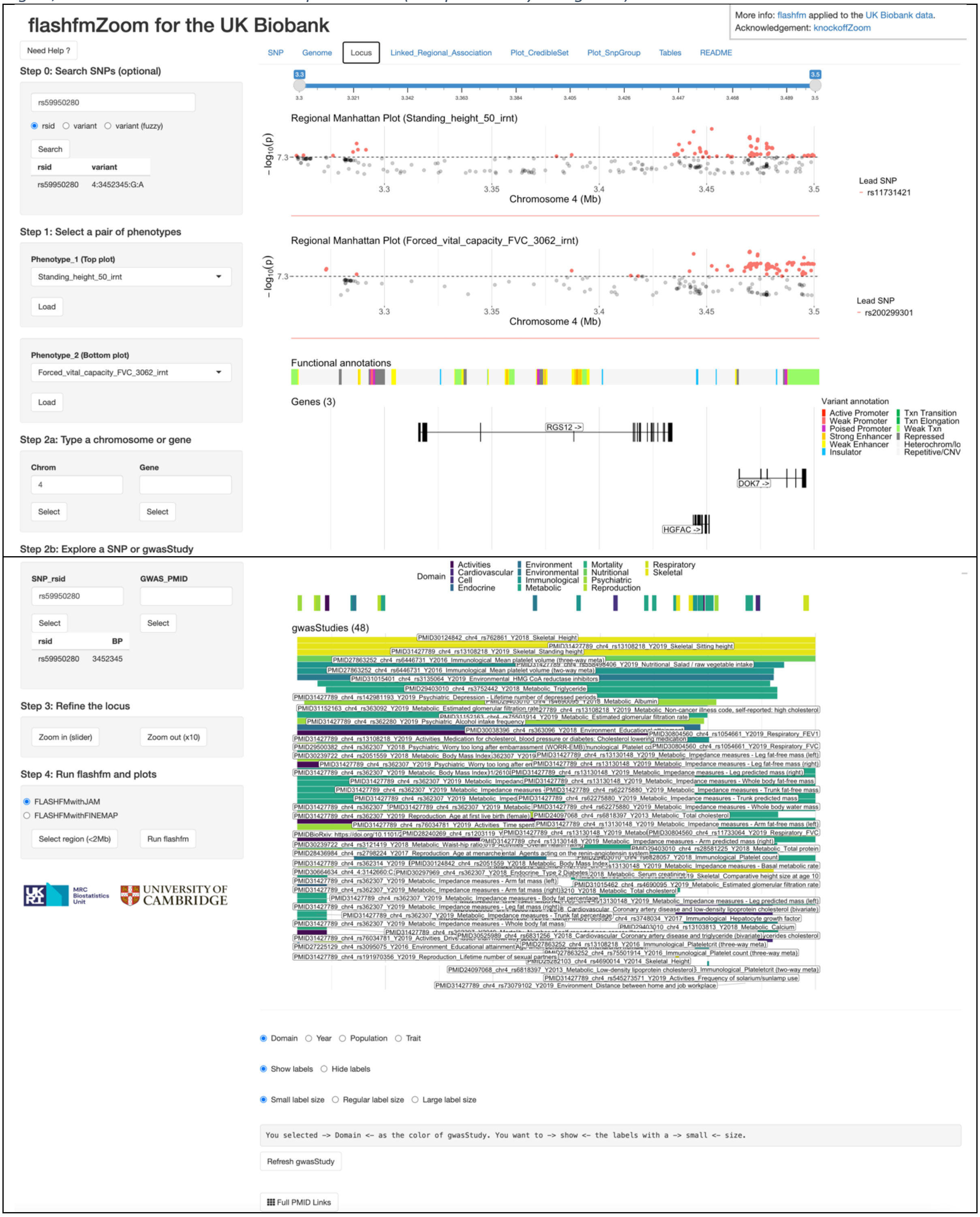

### S.4 FlashfmZoom details and instructions

**Table S2:** Left panel (control widgets) of flashfmZoom interface

| | |
|---|---|
| 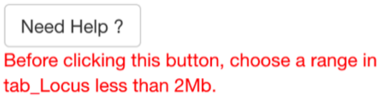 | The dynamic "Need Help?" button contains information that helps users to understand key features of each tab in the main panel, it provides unique information based on users' selected tab. If there is any error or warning message during the analysis, the text will be highlighted in red and displayed under the button. More details, see the Section in the end of this supplementary materials: Need help? |
| 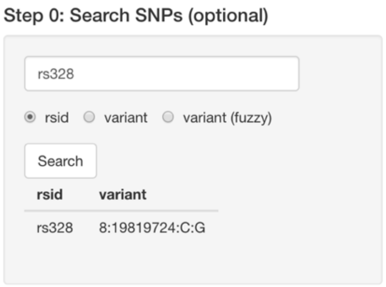 | Step_0 is optional but offers a useful way to search a single SNP for associated traits (i.e. PheWAS). Users can search a rs_id (e.g. rs328) or variant (e.g. 8:19819724:C:G) that matches the string exactly in the database of flashfmZoom. A fuzzy search (partial match) of a variant is also available. If there is more than one SNP output in the fuzzy search, users can select the most appropriate one. More details, see the Section below: Step_0 Search SNPs (optional). |
| 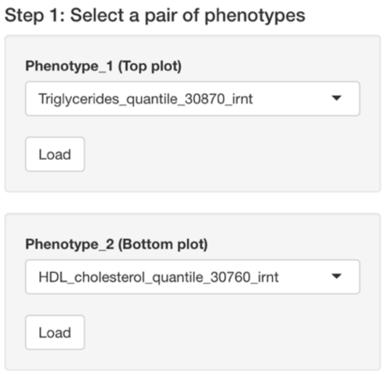 | Step_1 selects a pair of phenotypes. Users can use the outputs from Step_0 to inform their selection based on the selected SNP's pval in different traits. Also, users can use the drop-down list and options in the cell directly without searching SNPs. After uploading a pair of phenotypes, the right panel will show two Manhattan plots of these selected traits. More details, see the Section below: Step_1 Select a pair of phenotypes. |
| 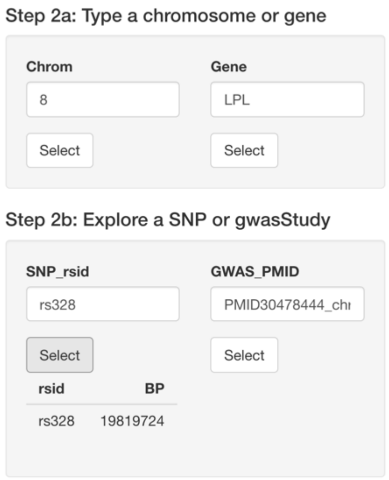 | Step_2 allows users to refine the SNP region, according to chromosome/gene in Step_2a and SNP/gwasStudy in Step_2b. In order to improve the speed of loading dataset (due to a large number of SNPs and existing gwasStudy publications in the whole dataset), Step_2b is a subset of Step_2a, which means that Step_2b is a refinement based on Step_2a. After users click one of the "select" buttons, the right panel will refine the "Locus" tab to show SNP/Gene/gwasStudy in the refined region by using PMID to search the literature, since it will be easier for researchers in this field to find a relevant paper. More details, see the Section below: Step_2a and Step_2b. |
| 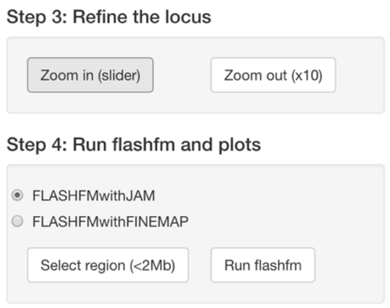 | Step_3 allows users to further adjust the region based on Zoom In/Out, by using the slider bar in the right panel to "Zoom in" or clicking "Zoom out" to increase the range of the existing region by 10 times.<br>Step_4 is the final step to run flashfm multi-trait fine-mapping, with two single-trait methods: (1) flashfm with FINEMAP software; (2) flashfm with JAM method. To run flashfm correctly and efficiently, there are two conditions before selecting a region: (1) the range must be less than 2Mb and (2) both traits have at least one SNP that has pval less than 1e-5. More details, see the Section below: Step_3 and Step_4. |

**Table S3:** Right panel (main tabs) of flashfmZoom interface

As shown below, the right panel has 8 main tabs, they interact with the left panel control widgets but can be clicked directly. Note: users can use the dynamic “Need Help?” button to check more information and read useful instructions (or error/warning messages) when using these tabs.

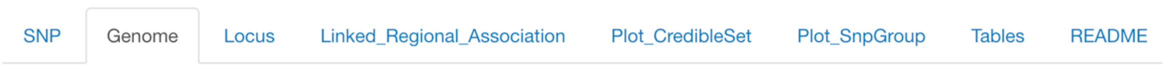


| | |
|---|---|
| SNP | This “tab: SNP” shows the -log10(pval) results of traits that are associated with the searched SNP in Step_0 among all 134 traits in the flashfmZoom package; displayed traits have -log10(pval) that meet the users’ defined threshold: pval<1e-6 or pval<5e-8 (as well as a default threshold pval<1e-5). Users can also sort the bar plot based on trait name or pval. The trait correlation matrix (with a heatmap) of displayed traits is also shown on the page, users can sort and download it as a csv/excel table.<br><br>More details, see the Section below: Step_0 Search SNPs (optional).<br>The full list of 134 traits can be found in Section: Supplementary Table S1. |
| Genome | This “tab: Genome” shows two Manhattan plots that were selected in the Step_1. Users can check the pval of SNPs in these plots, to refine a chromosome.<br><br>More details, see the Section below: Step_1 Select a pair of phenotypes. |
| Locus | This “tab: Locus” shows a particular region that users refined in the previous steps. The top slider bar refines the range of this region, for smaller regions, more detailed information (e.g. gene names, rs_id in the existing GWAS publications) will be displayed in the charts of this tab.<br><br>The two regional Manhattan plots highlight the lead SNPs according to the pval in different sub-regions, other SNPs around the lead SNP are highlighted in the same colour. Please note, the name of the lead SNP in the legend states the SNP with the minimum pval (or the first SNP if there are more than one SNP with the minimum pval) in each subset of Manhattan plots.<br><br>Both Gene names and functional annotations are displayed below the regional Manhattan plots according to the same range. A list of existing GWAS studies/publications within this region is also attached below Gene. The publications can be searched by the PMID, as well as sorted and coloured by the year/domain/population/trait. Users can adjust the font of labels in the publication list in order to refine the region.<br><br>More details, see the Section below: Step_2a and Step_2b. |
| Linked_Regional_Association | This “tab: Linked_Regional_Association” contains detailed sub-steps of running the flashfm R package:<br><br>(1) It informs the users to download the LD matrix based on the selected region in the previous steps. Since it may take at least 3 minutes to download the LD matrix, the tab informs users to check the selected files carefully before downloading the data.<br><br>(2) After downloading the LD matrix, the tab will disable the download button (as well as the “Select region (<2MB)” button in the left panel) in order to avoid downloading a new LD matrix. |

| | |
|---|---|
| | (3) The middle slider bar control widget will be shown after users downloaded the LD matrix, so they can further refine a sub-region. Usually, a region that contains less than 1,500 SNPs and has at least one SNP with pval less than 1e-6 in both traits will run flashfm efficiently (Note: these required SNPs with pval<1e-6 can be different in both traits). Users can also use numeric inputs of positions to refine the region and select a MAF threshold.<br><br>(4) Users can also check the LD matrix based on their selected sub-region by looking at the two correlation heatmaps below the slider bar. The first left-side heatmap shows the correlation of first 10 SNPs, while the second right-side heatmap shows the last 10 SNPs. It takes time to load a heatmap of the whole LD matrix, therefore only these two parts are displayed for users to double check the data inputs.<br><br>(5) Once users are happy with the selected sub-region and its LD matrix, they can click "Run flashfm" to run the flashfm R package directly from the website. The default method is "FLASHFMwithJAM", but users can select FINEMAP software by clicking the "FLASHFMwithFINEMAP" option.<br><br>(6) The summary results of flashfm will be displayed below the two LD correlation heatmaps. The coloured bar table contains all SNP groups, while the linked interactive plots display these SNP groups on the regional association plots. Please note, users can interact with the plots by using control widgets and clicking the buttons on the plots. More details, see the Section below: Step_3 and Step_4. |
| Plot_CredibleSet | This "tab: Plot_CredibleSet" focuses on displaying credible sets in the current regional association analysis. Users can re-define different levels of credible sets in this region. Four regional association plots for single- and multi-trait fine-mapping (i.e. 2 traits x 2 fine-mapping). Venn diagrams show a clearer view about the duplications/intersections of SNPs in different credible sets. Users can also download the tables.<br><br>More details, see the Section below: Credible Sets |
| Plot_SnpGroup | This "tab: Plot_SnpGroup" focuses on displaying the link between individual SNPs and their SNP groups in both single- and multi-trait fine-mapping. Users can have a clearer view about the names of SNPs in each group.<br><br>More details, see the Section below: SnpGroup |
| Tables | This "tab: Tables" contains all information and outputs that were used in the previous tabs, to be more transparent for users to check the results. The full table of GWAS association results can be downloaded as well, users can run further analyses based on this table in their local machine.<br><br>More details, see the Section below: Downloadable tables |
| README | This "tab: README" contains other relevant information, as well as a few YouTube videos for helping users to use this package.<br><br>Again, users can use the dynamic "NEED HELP?" button in the top corner of left panel to check useful instructions (or to read error/warning messages) if needed. |

### Step_0: Search SNPs (optional)

In addition to the information and instructions in **Table S1/S2/S3** above, the use of interactive modebar icons in the plots can be found in **Table S4** at the end of this Supplementary Material.

Note:
(1) The "fuzzy search" can be used if users remember the variant names partially.
(2) The SNP variant info (e.g. rs_id or chr_id) under the "Search" button in Step_0 can be used in Step_2.
(3) Users can simply use the trait_id (e.g. 30780 for LDL) or the full name to input the Phenotypes in Step_1.

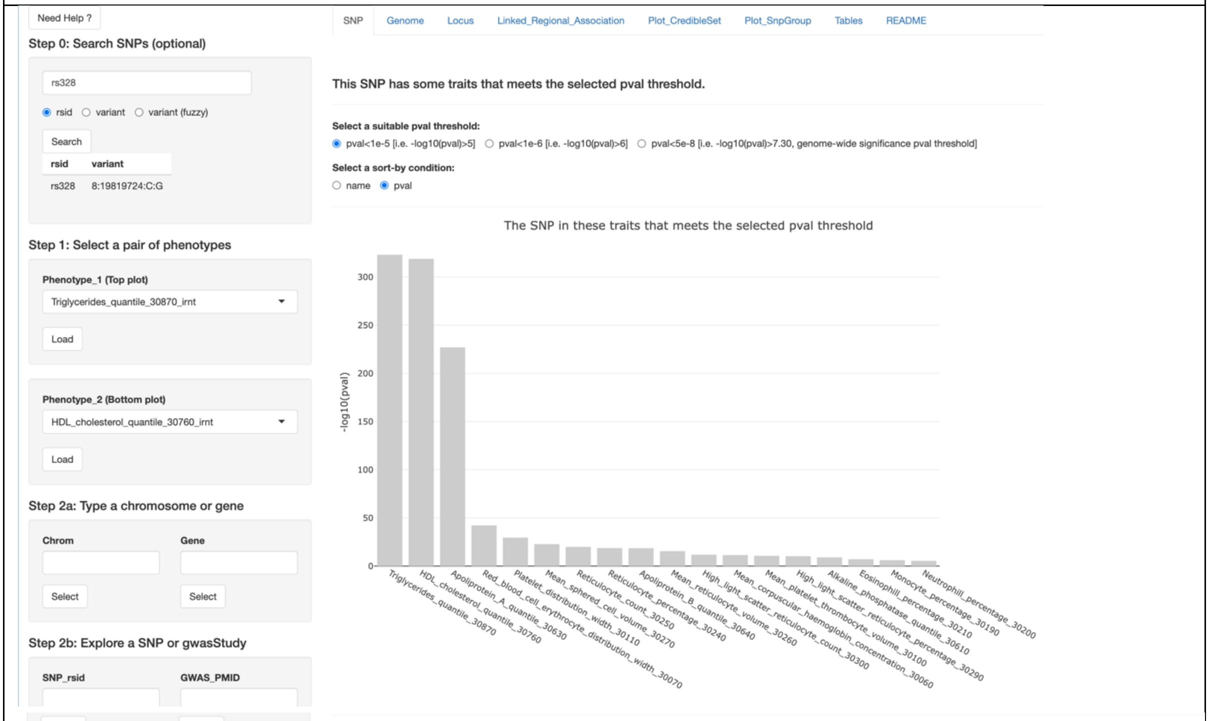


Trait correlation table

| | _30240 | trait_30250 | trait_30260 | trait_30270 | trait_30290 | trait_30300 | trait_30610 | trait_30630 | trait_30640 | trait_30760 | trait_30870 |
|---|---|---|---|---|---|---|---|---|---|---|---|
| | All | All | All | All | All | All | All | All | All | All | All |
| trait_30060 | 0.067 | 0.03 | -0.071 | -0.021 | 0.054 | 0.026 | -0.026 | -0.013 | 0.023 | -0.03 | 0.082 |
| trait_30070 | -0.018 | -0.009 | 0.139 | 0.011 | 0.026 | 0.032 | 0.058 | -0.028 | -0.021 | -0.004 | -0.043 |
| trait_30100 | -0.01 | 0.001 | -0.038 | 0.002 | -0.02 | -0.012 | 0.004 | -0.064 | -0.06 | -0.055 | -0.008 |
| trait_30110 | 0.089 | 0.098 | -0.11 | -0.126 | 0.07 | 0.078 | 0.031 | -0.089 | 0.012 | -0.113 | 0.151 |
| trait_30190 | -0.042 | -0.05 | 0.017 | 0.016 | -0.038 | -0.045 | -0.036 | 0.011 | -0.046 | 0.029 | -0.057 |
| trait_30200 | 0.06 | 0.069 | 0.001 | -0.027 | 0.061 | 0.068 | 0.071 | 0.006 | -0.043 | -0.003 | -0.008 |
| trait_30210 | 0.001 | 0.002 | -0.036 | -0.047 | 0.003 | 0.003 | 0.016 | -0.05 | 0.006 | -0.057 | 0.038 |
| trait_30240 | 1 | 0.98 | -0.152 | -0.188 | 0.912 | 0.903 | 0.085 | -0.122 | 0.068 | -0.22 | 0.308 |
| trait_30250 | 0.98 | 1 | -0.214 | -0.258 | 0.889 | 0.911 | 0.107 | -0.137 | 0.096 | -0.243 | 0.33 |
| trait_30260 | -0.152 | -0.214 | 1 | 0.744 | -0.017 | -0.066 | -0.007 | 0.113 | -0.044 | 0.145 | -0.139 |
| trait_30270 | -0.188 | -0.258 | 0.744 | 1 | -0.091 | -0.148 | -0.058 | 0.194 | -0.042 | 0.235 | -0.169 |
| trait_30290 | 0.912 | 0.889 | -0.017 | -0.091 | 1 | 0.986 | 0.094 | -0.13 | 0.061 | -0.232 | 0.315 |
| trait_30300 | 0.903 | 0.911 | -0.066 | -0.148 | 0.986 | 1 | 0.111 | -0.143 | 0.083 | -0.251 | 0.333 |
| trait_30610 | 0.085 | 0.107 | -0.007 | -0.058 | 0.094 | 0.111 | 1 | -0.111 | 0.05 | -0.127 | 0.154 |
| trait_30630 | -0.122 | -0.137 | 0.113 | 0.194 | -0.13 | -0.143 | -0.111 | 1 | -0.047 | 0.908 | -0.27 |
| trait_30640 | 0.068 | 0.096 | -0.044 | -0.042 | 0.061 | 0.083 | 0.05 | -0.047 | 1 | -0.039 | 0.345 |
| trait_30760 | -0.22 | -0.243 | 0.145 | 0.235 | -0.232 | -0.251 | -0.127 | 0.908 | -0.039 | 1 | -0.474 |
| trait_30870 | 0.308 | 0.33 | -0.139 | -0.169 | 0.315 | 0.333 | 0.154 | -0.27 | 0.345 | -0.474 | 1 |

CSV Excel

### Step_1: Select a pair of phenotypes or traits

In addition to the information and instructions in **Table S1/S2/S3** earlier, if users click the "Need Help?" in the top-left of flashfmZoom interface (see below), the relevant instructions will be displayed accordingly as shown in the right cell here.

Note:
(1) All 134 traits in the current version of flashfmZoom are from the Neale lab (see data source), here we only select the European ancestry (not trans-ancestry), quantitative traits (not case-control for diseases), and male+female version instead of gender separated.
(2) All Manhattan plots are created by this R code: [https://genome.sph.umich.edu/wiki/Code_Sample:_Generating_Manhattan_Plots_in_R](https://genome.sph.umich.edu/wiki/Code_Sample:_Generating_Manhattan_Plots_in_R)

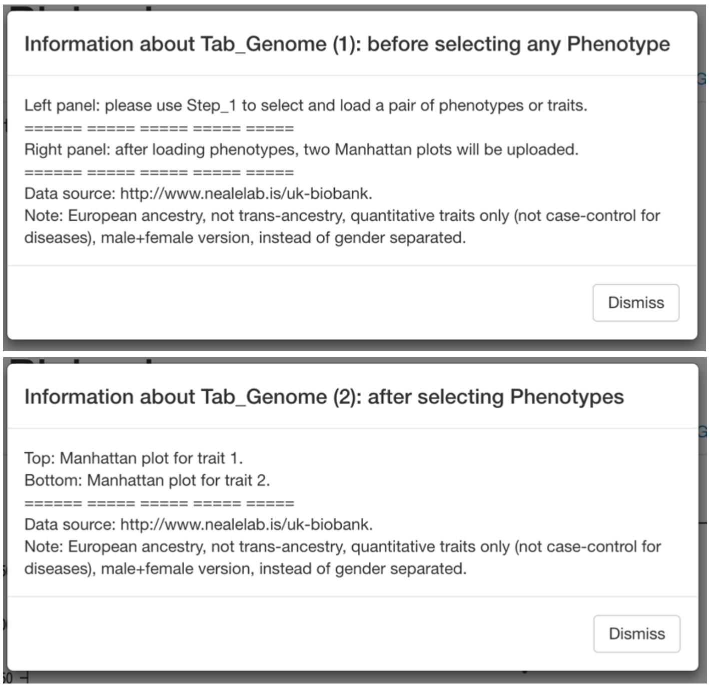


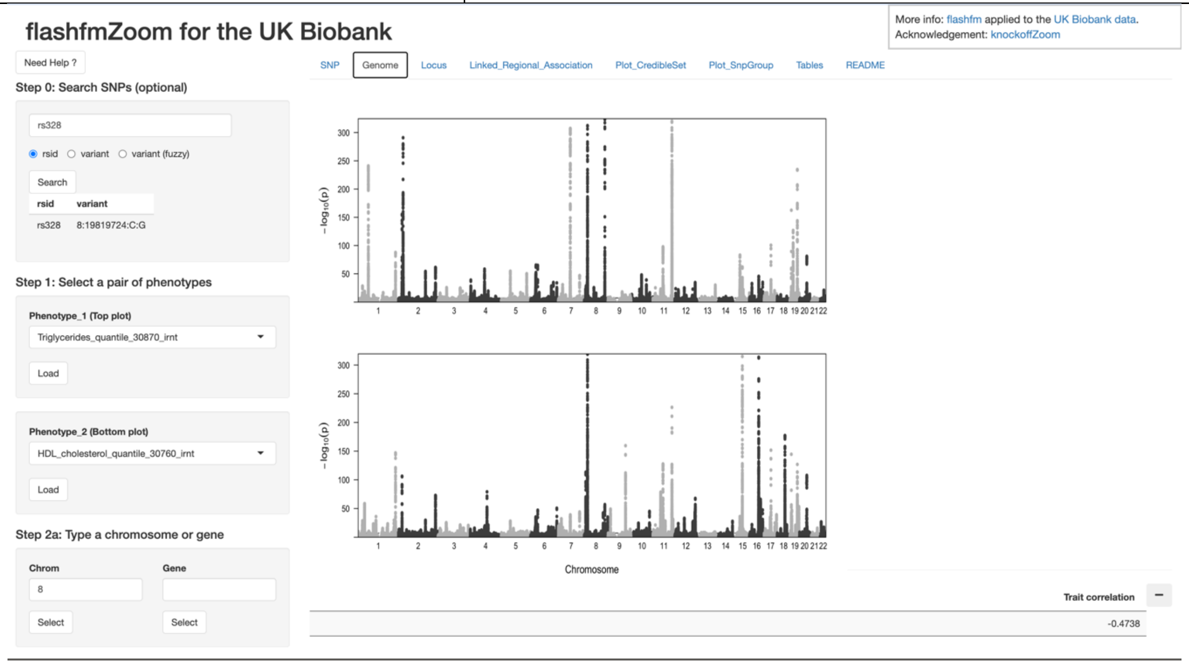

### Step_2a: Type a chromosome or gene

In addition to the information and instructions in **Table S1/S2/S3** earlier, if users click the "Need Help?" in the top-left of flashfmZoom interface (see below), the relevant instructions will be displayed accordingly (see right cell).

Note: After users input the chr_id in the Step_2a "Chrom", the right-side main panel will display the selected whole chromosome. Both Manhattan plots will be refined to regional association plots. As an initial view, there are too many genes in this region, so users can refine the region to see the genes and other detailed information (e.g. the list of existing GWAS publications in a corresponding region).

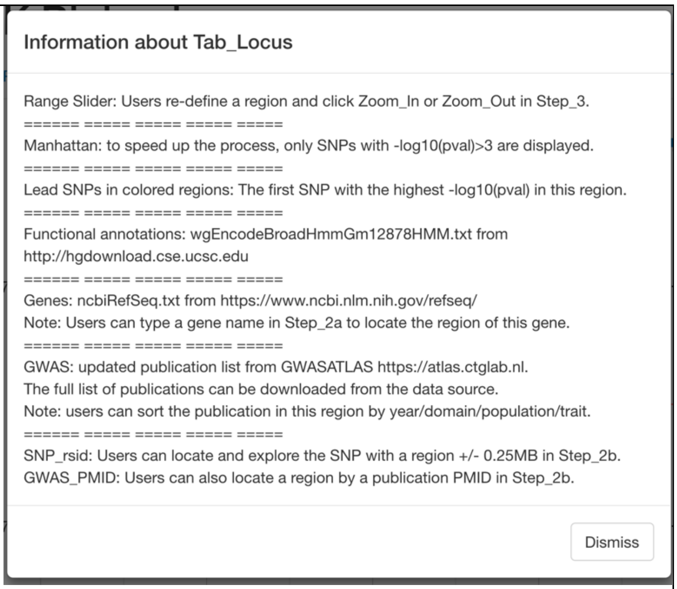


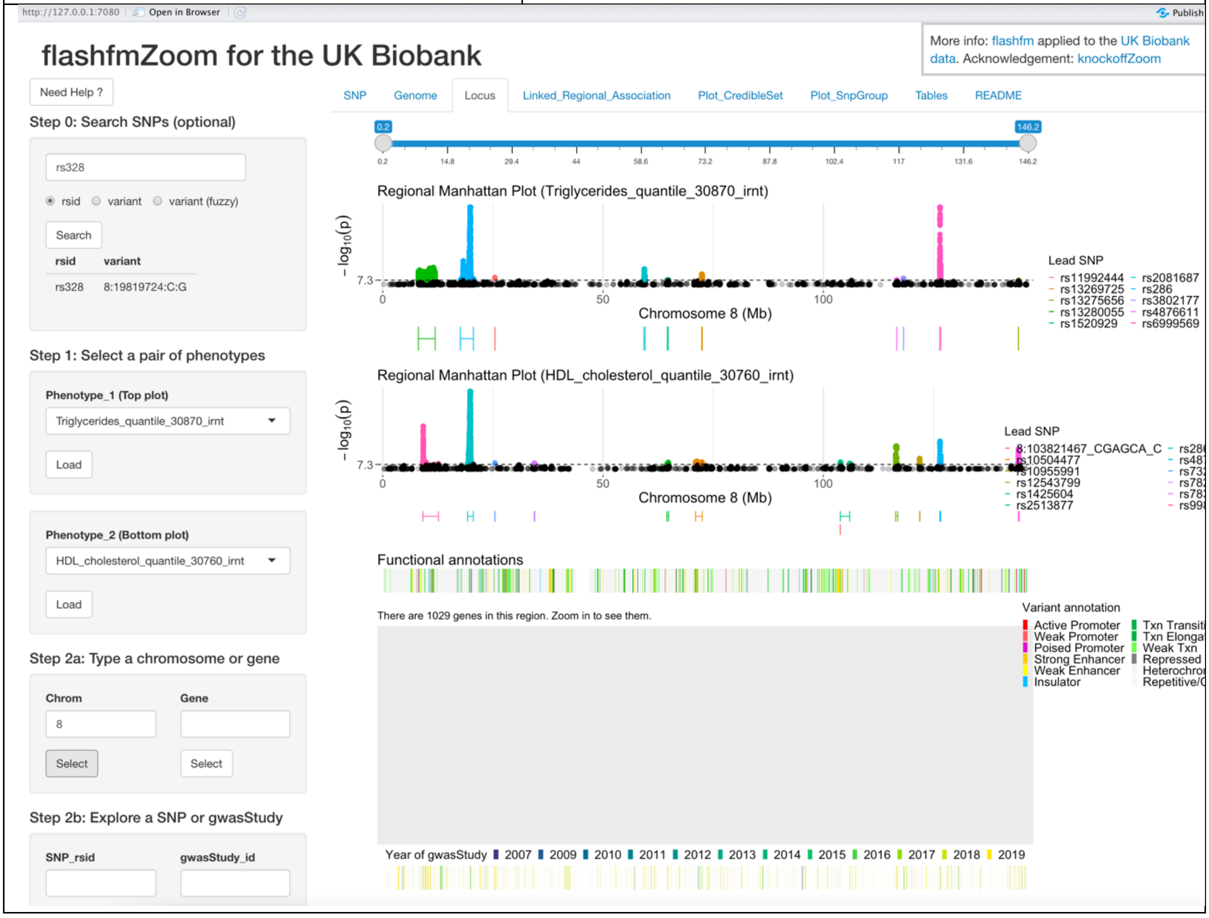

Users can select the region based on either chr_id or a specific gene name in any Chromosome to locate the regional Manhattan plots.

After users refine/narrow the region by using the top blue slider bar, more detailed plots will be displayed. Once the number of genes is reduced to 50 or less, both the names and the locations of genes will be displayed. Users can use options in Step_3 to Zoom In/Out the region/locus (see right). All gene names can be found in the drop-down list of Step_2a (e.g. LPL here).

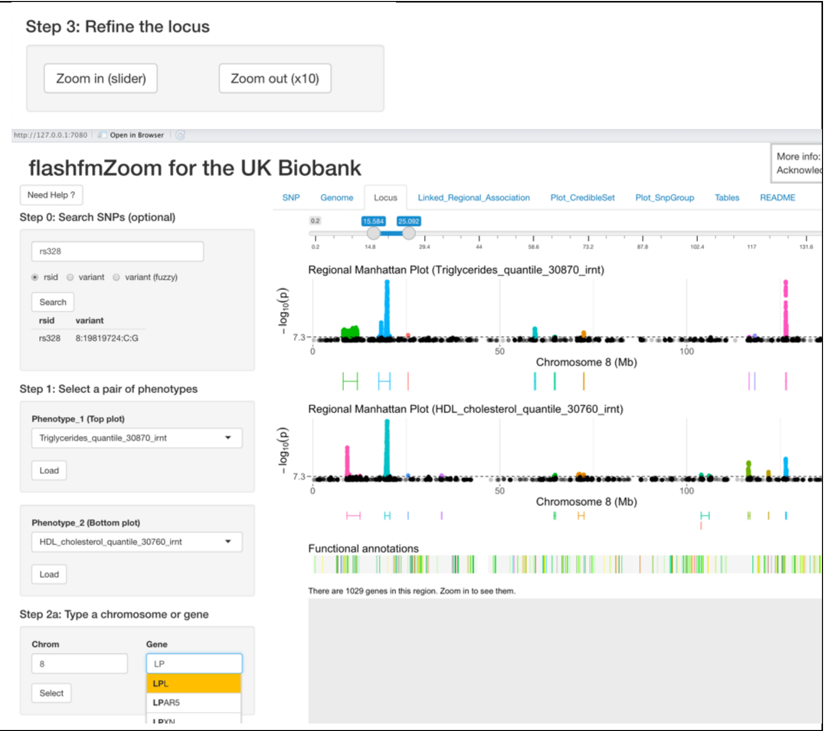


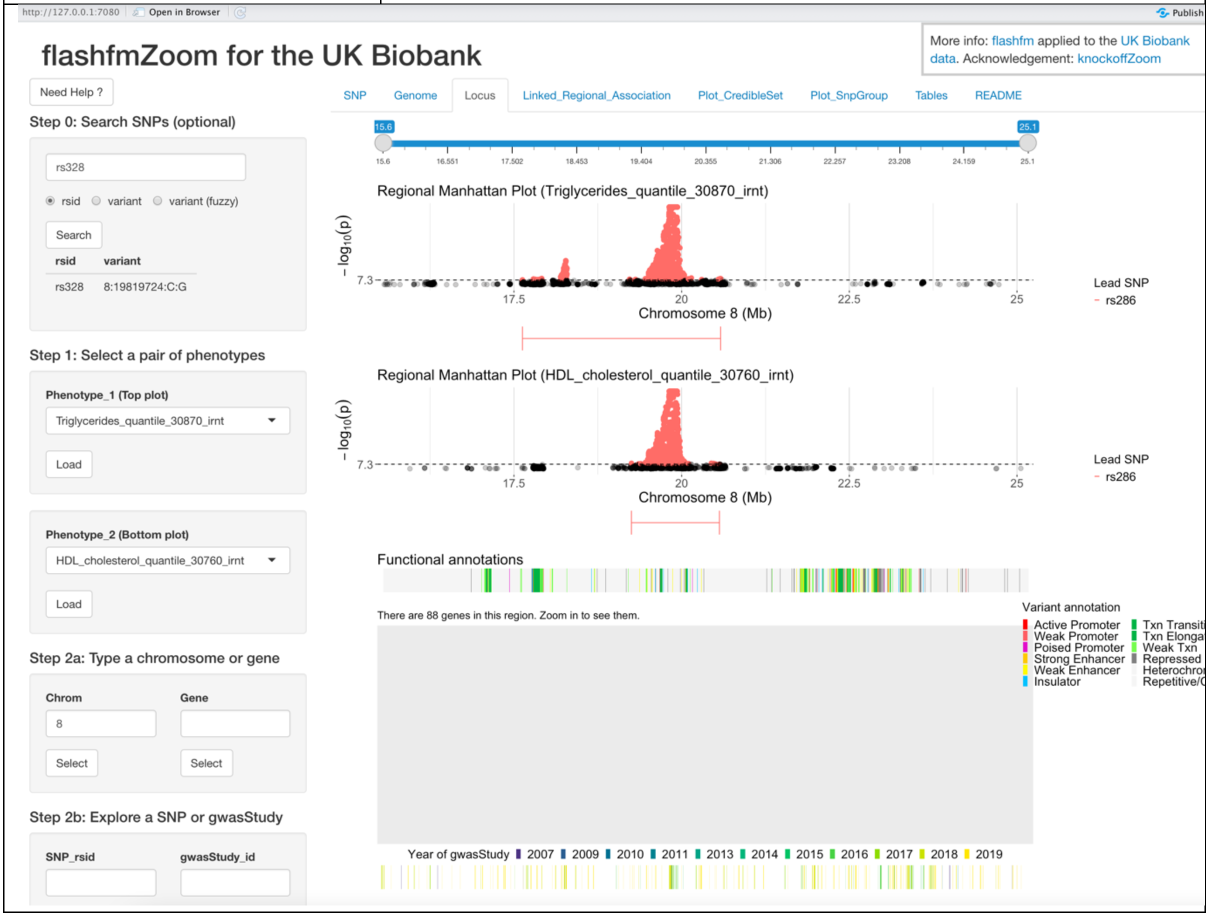

### Step_2b: Explore a SNP or GAWS

Users can refine the region by entering a focal SNP or a gwasStudy_id in Step_2b (as an optional sub-step). In order to show the names of gene and GWAS publications in the main panel, the default range of this selected region is small (i.e. 500Kb, the position of the entered SNP or the lead SNP in the gwasStudy is the middle location +/- 250Kb). For example as below, the SNP "rs328" is located at "BP=19819724", while the region in the main panel is between 19.6Mb and 20.1Mb.

As shown in next page, GWAS publications can be displayed and coloured by the Year, Domain, Population, also the type of Traits.

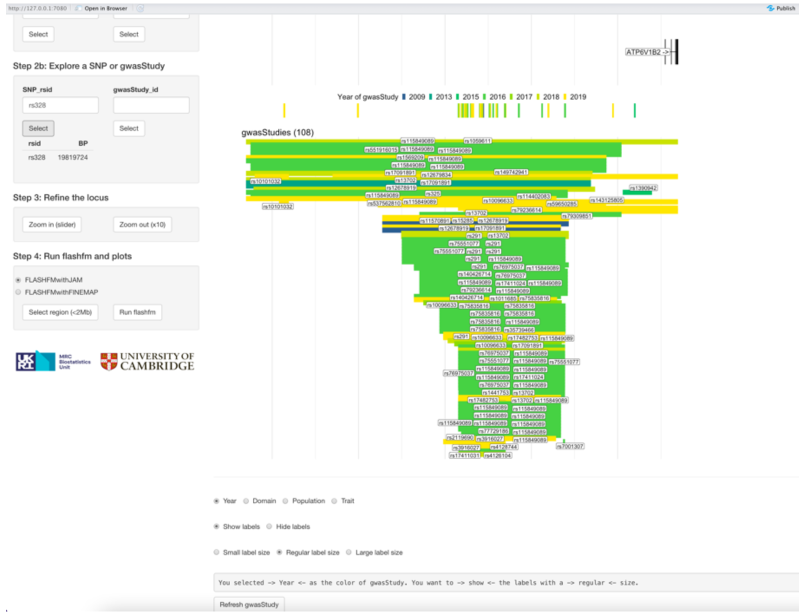


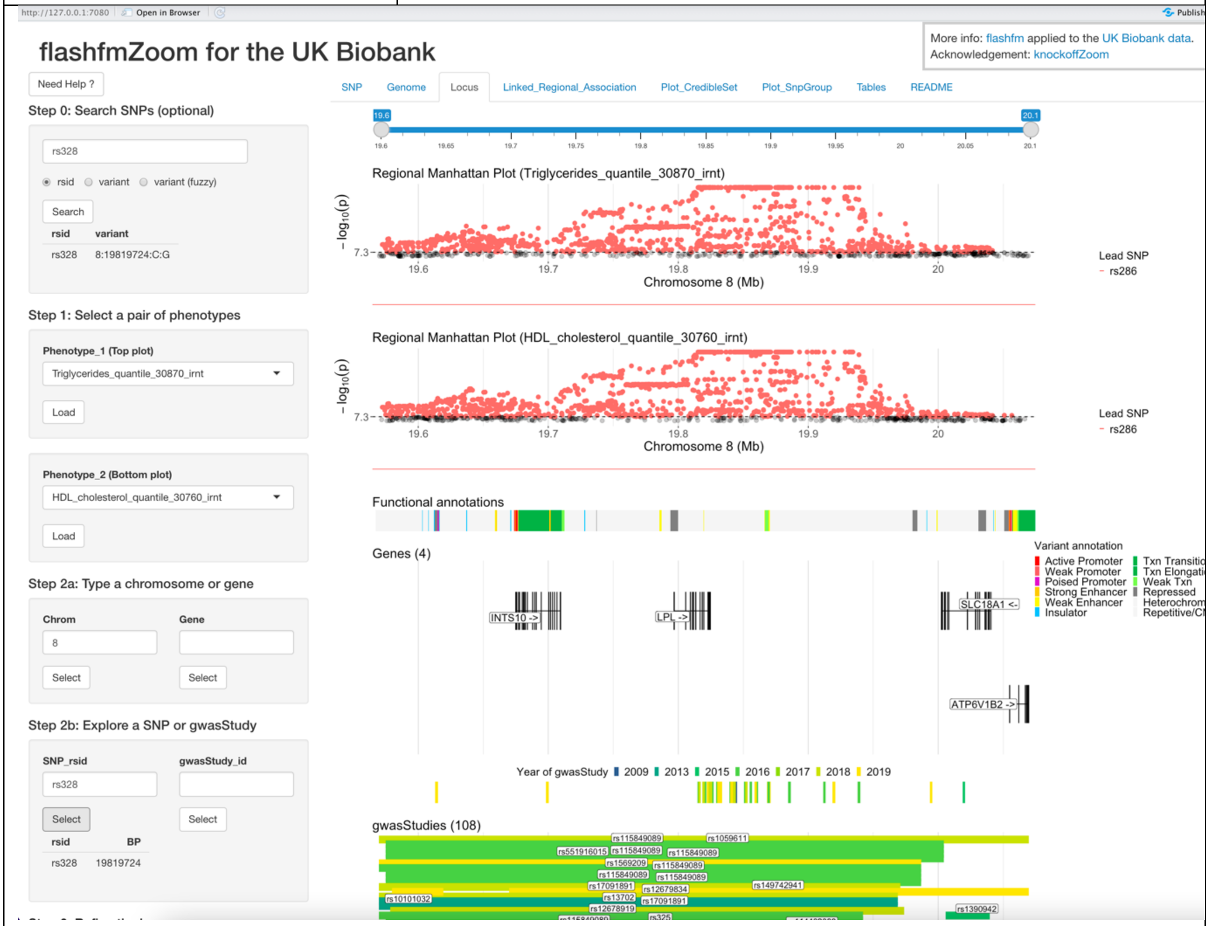

Users can apply the control widgets in the bottom of the main panel (see right cell here) to display GWAS publications by the year/domain/population/tye of traits. The labels in the plot can be hidden if needed. The size of labels can also be adjusted for a more comfortable read.

The little vertical bars under the legend indicate the exact positions of lead SNPs in these GWAS publications, while the horizontal long bars indicate the range/region around the lead SNPs. All colours are matched consistently between legend and plots.

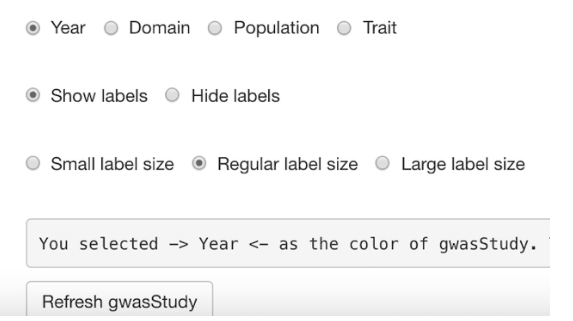


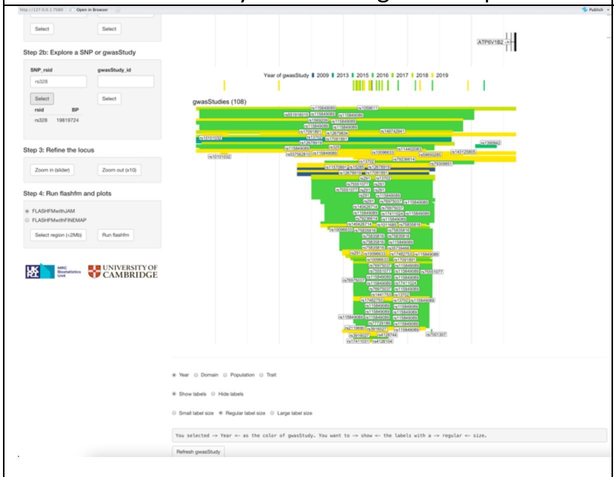

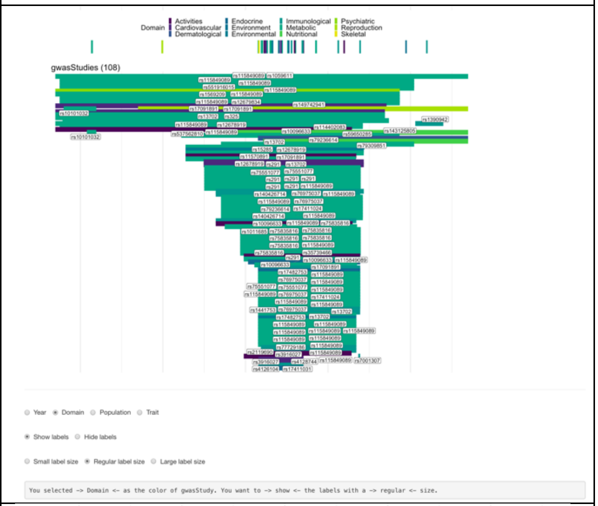

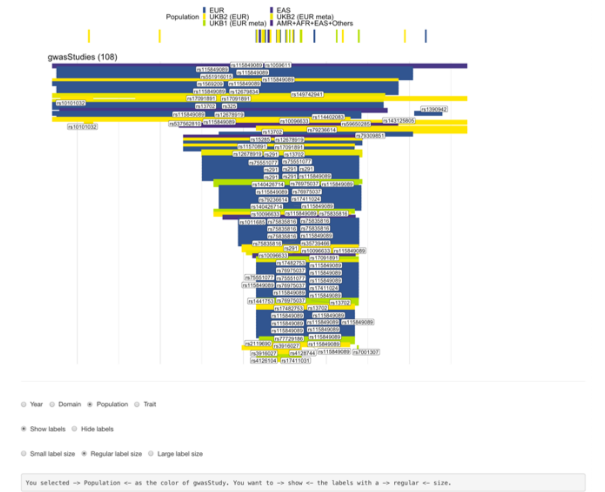

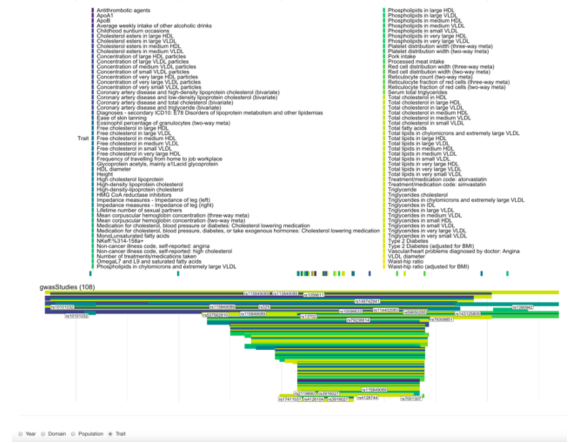

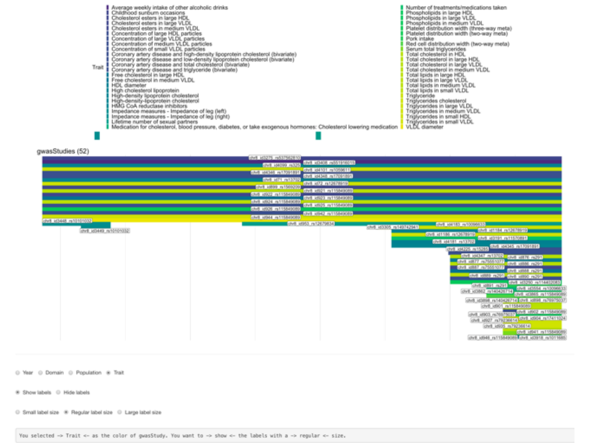

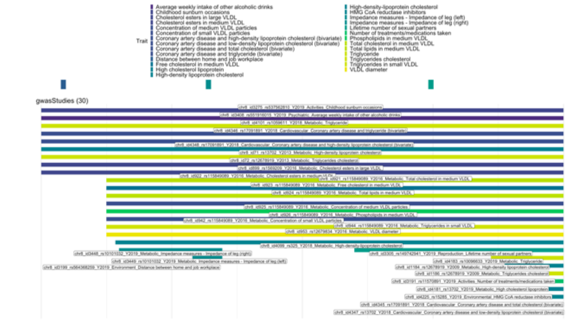

When users reduce/narrow the region further (see below), more information about the GWAS publications will be displayed. The unique label (e.g. PMID21926974) can be used in the Step_2b (optional) as a focal GWAS_PMID (see right).

Note: the labels on the coloured horizontal long bars may have different formats when the number of GWAS publications is small or large, in order to display the information more clearly. For example, if the number of acticles is more than 150, then only the rsid of lead SNPs will be displayed. If the number of articles is not more than 50, then more information will be displayed, such as PMID, chr_id, rsid, etc.

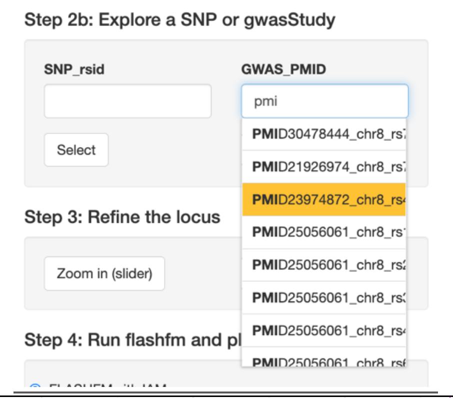


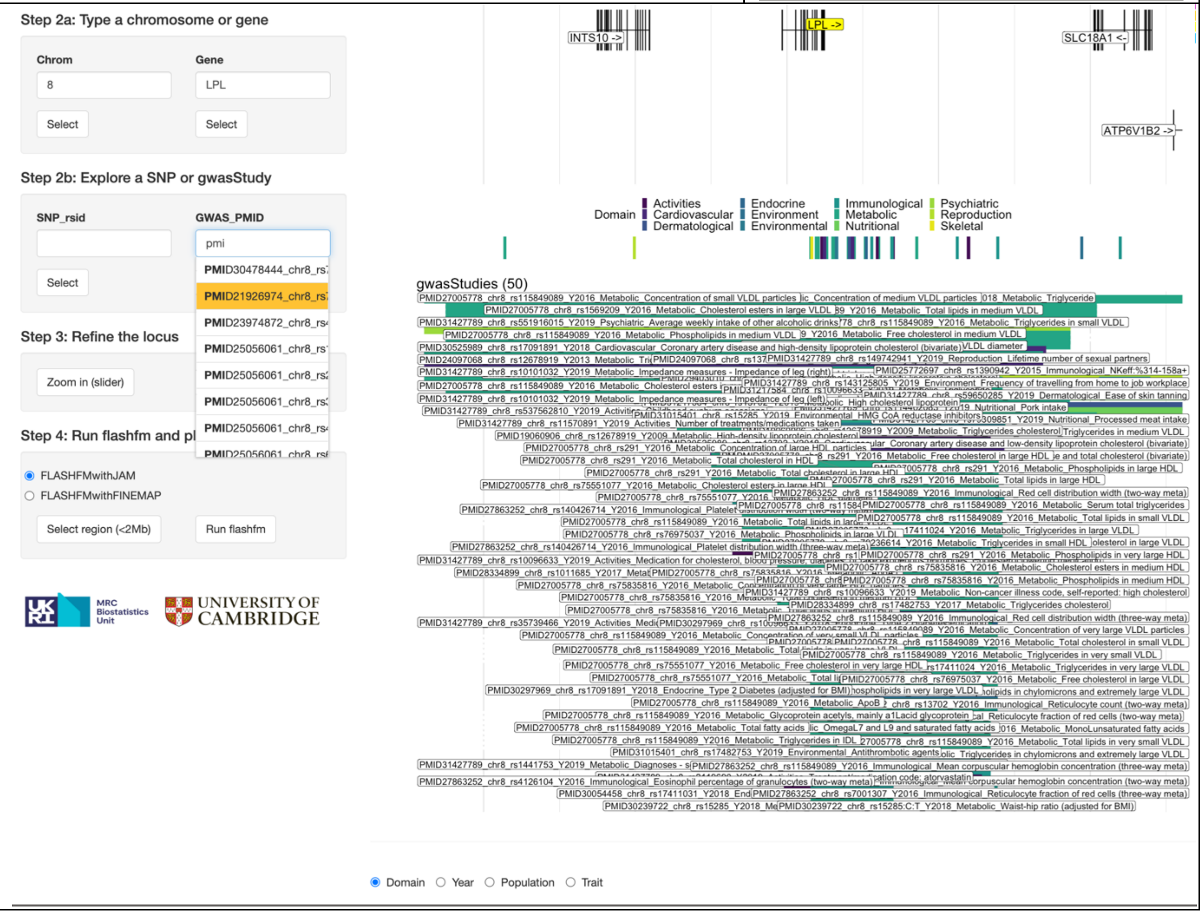

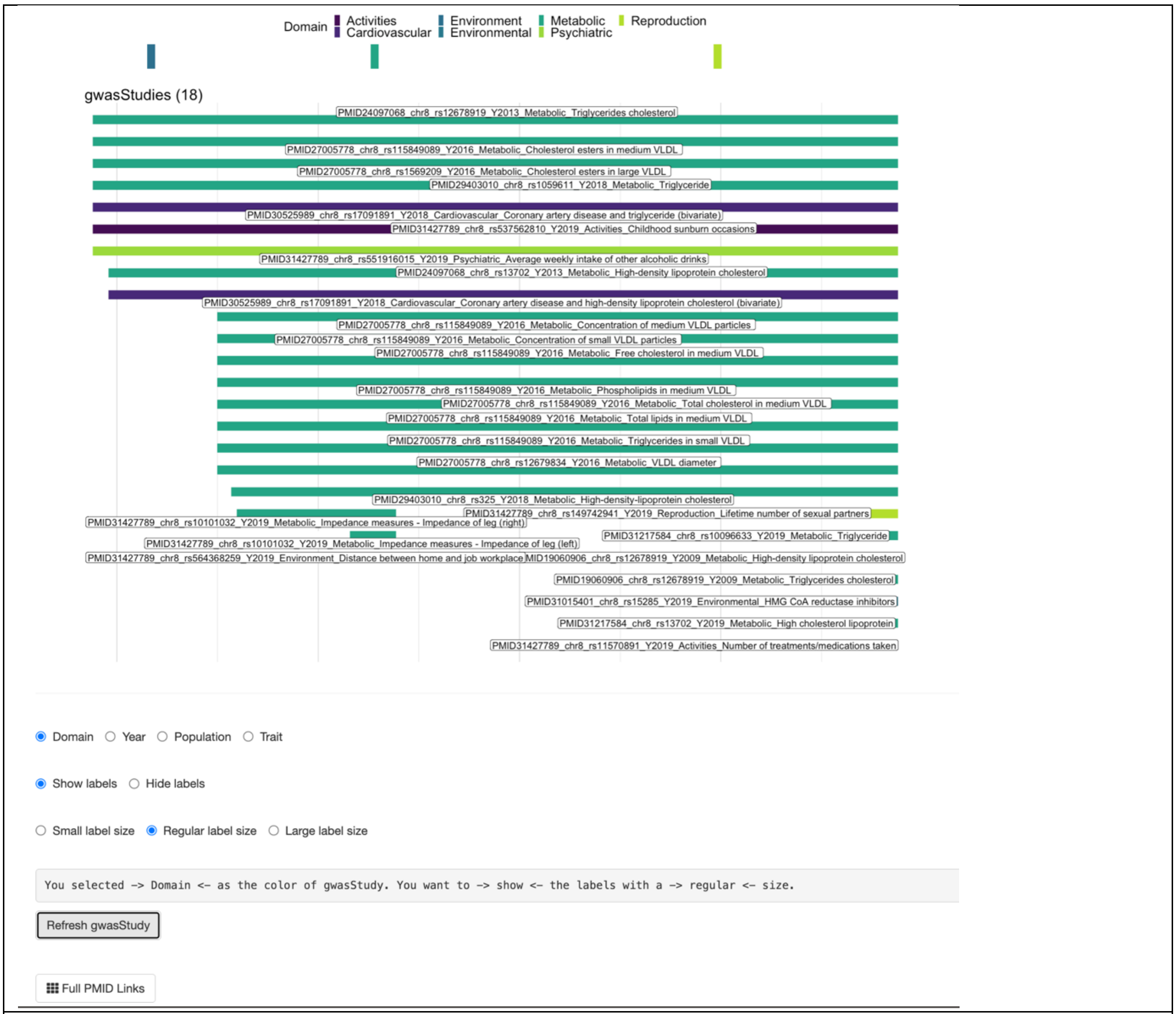
Domain Activities Cardiovascular Environment Environmental Metabolic Psychiatric Reproduction
gwasStudies (18)
PMID24097068_chr8_rs12678919_Y2013_Metabolic_Triglycerides cholesterol
PMID27005778_chr8_rs115849089_Y2016_Metabolic_Cholesterol esters in medium VLDL
PMID27005778_chr8_rs1569209_Y2016_Metabolic_Cholesterol esters in large VLDL
PMID29403010_chr8_rs1059611_Y2018_Metabolic_Triglyceride
PMID30525989_chr8_rs17091891_Y2018_Cardiovascular_Coronary artery disease and triglyceride (bivariate)
PMID31427789_chr8_rs537562810_Y2019_Activities_Childhood sunburn occasions
PMID31427789_chr8_rs551916015_Y2019_Psychiatric_Average weekly intake of other alcoholic drinks
PMID24097068_chr8_rs13702_Y2013_Metabolic_High-density lipoprotein cholesterol
PMID30525989_chr8_rs17091891_Y2018_Cardiovascular_Coronary artery disease and high-density lipoprotein cholesterol (bivariate)
PMID27005778_chr8_rs115849089_Y2016_Metabolic_Concentration of medium VLDL particles
PMID27005778_chr8_rs115849089_Y2016_Metabolic_Concentration of small VLDL particles
PMID27005778_chr8_rs115849089_Y2016_Metabolic_Free cholesterol in medium VLDL
PMID27005778_chr8_rs115849089_Y2016_Metabolic_Phospholipids in medium VLDL
PMID27005778_chr8_rs115849089_Y2016_Metabolic_Total cholesterol in medium VLDL
PMID27005778_chr8_rs115849089_Y2016_Metabolic_Total lipids in medium VLDL
PMID27005778_chr8_rs115849089_Y2016_Metabolic_Triglycerides in small VLDL
PMID27005778_chr8_rs12679834_Y2016_Metabolic_VLDL diameter
PMID29403010_chr8_rs325_Y2018_Metabolic_High-density-lipoprotein cholesterol
PMID31427789_chr8_rs149742941_Y2019_Reproduction_Lifetime number of sexual partners
PMID31427789_chr8_rs10101032_Y2019_Metabolic_Impedance measures - Impedance of leg (right)
PMID31217584_chr8_rs10096633_Y2019_Metabolic_Triglyceride
PMID31427789_chr8_rs10101032_Y2019_Metabolic_Impedance measures - Impedance of leg (left)
PMID31427789_chr8_rs564368259_Y2019_Environment_Distance between home and job workplace
MID19060906_chr8_rs12678919_Y2009_Metabolic_High-density lipoprotein cholesterol
PMID19060906_chr8_rs12678919_Y2009_Metabolic_Triglycerides cholesterol
PMID31015401_chr8_rs15285_Y2019_Environmental_HMG CoA reductase inhibitors
PMID31217584_chr8_rs13702_Y2019_Metabolic_High cholesterol lipoprotein
PMID31427789_chr8_rs11570891_Y2019_Activities_Number of treatments/medications taken
Domain Year Population Trait
Show labels Hide labels
Small label size Regular label size Large label size
You selected -> Domain <- as the color of gwasStudy. You want to -> show <- the labels with a -> regular <- size.
Refresh gwasStudy
Full PMID Links

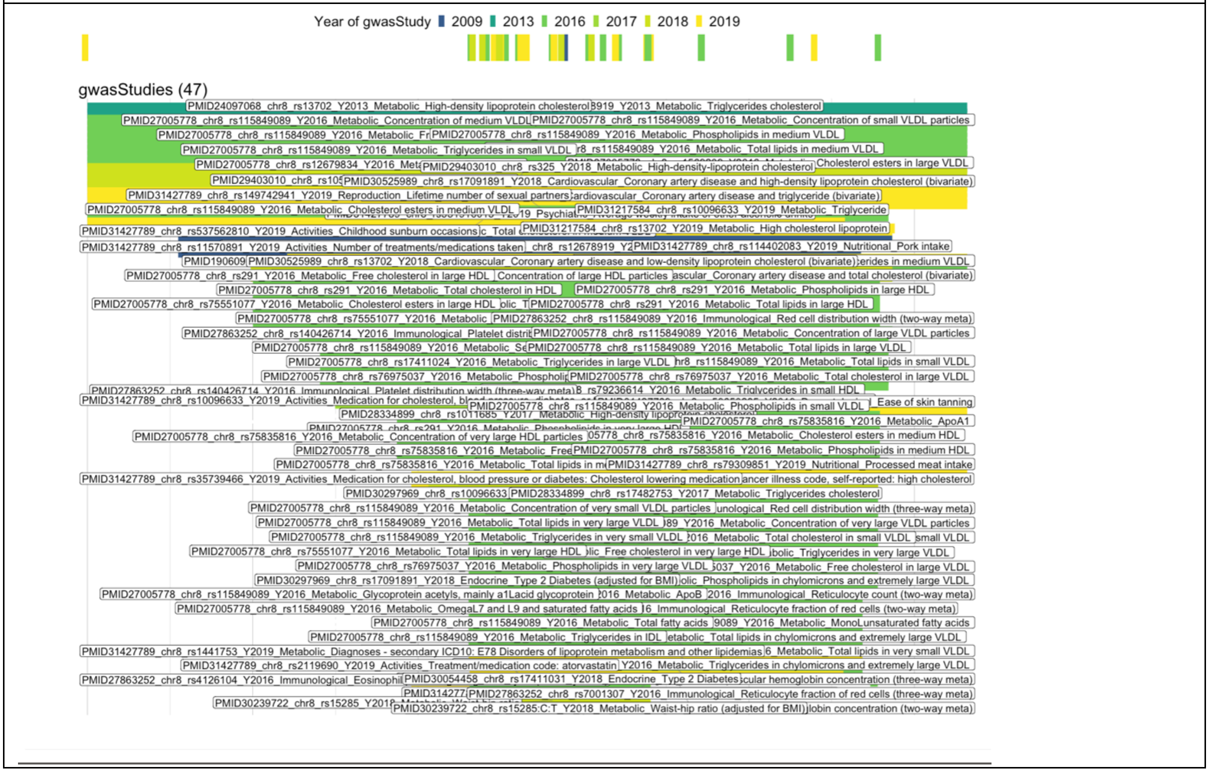
Year of gwasStudy 2009 2013 2016 2017 2018 2019
gwasStudies (47)

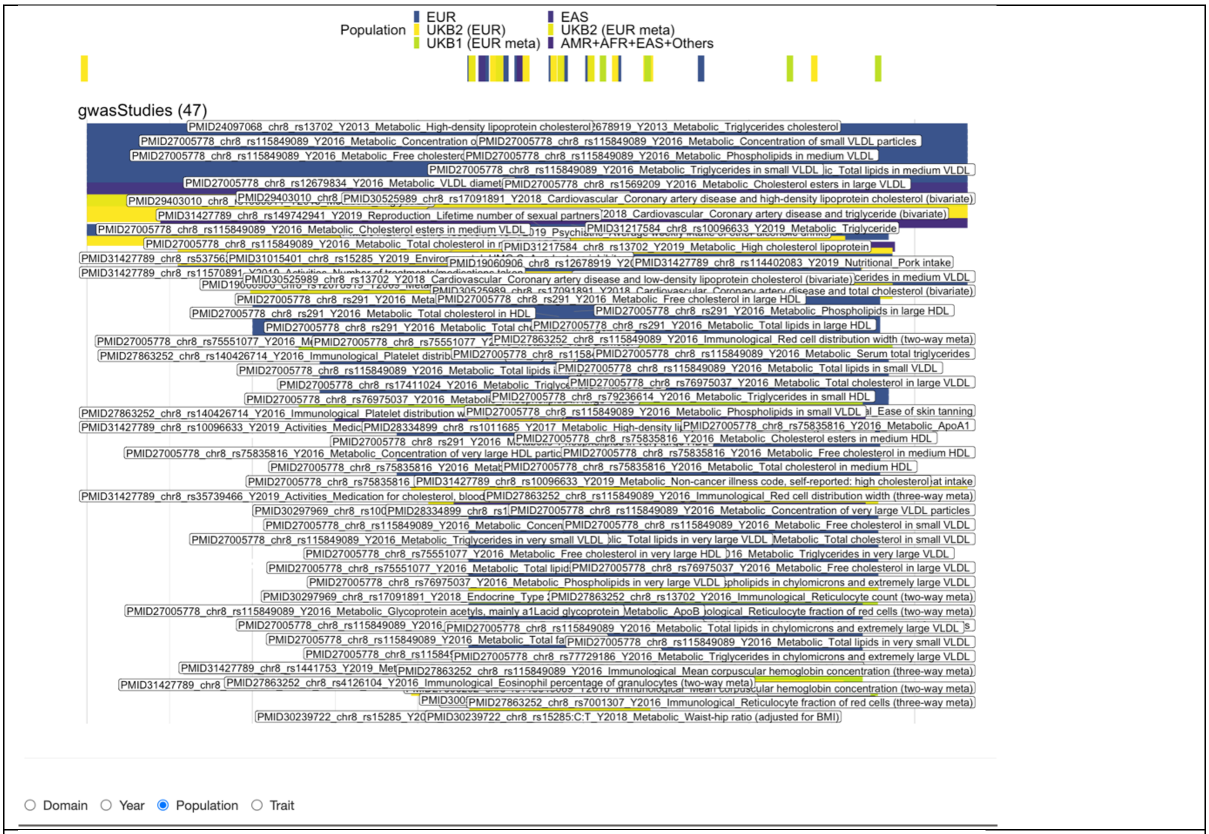
Population
EUR
UKB2 (EUR)
UKB1 (EUR meta)
EAS
UKB2 (EUR meta)
AMR+AFR+EAS+Others
gwasStudies (47)
Domain
Year
Population
Trait

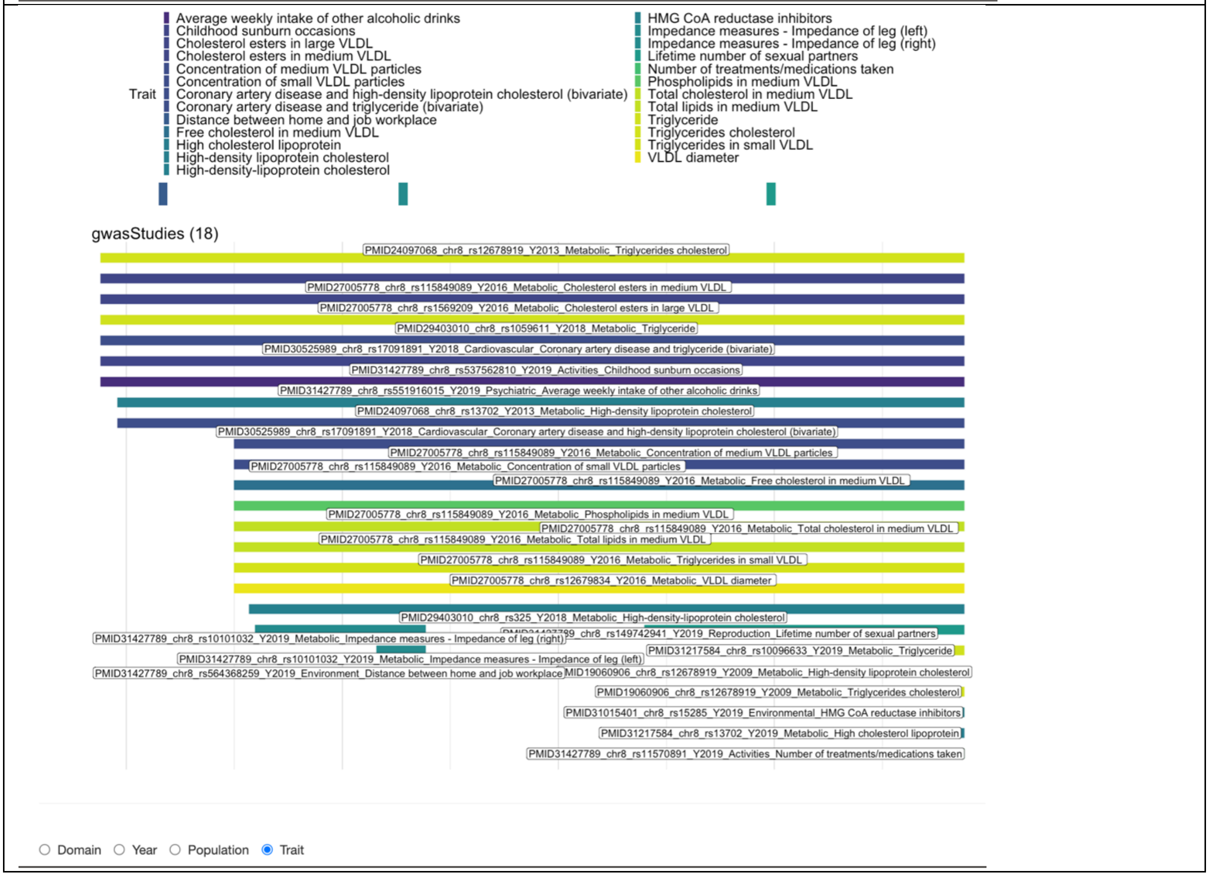
Trait
Average weekly intake of other alcoholic drinks
Childhood sunburn occasions
Cholesterol esters in large VLDL
Cholesterol esters in medium VLDL
Concentration of medium VLDL particles
Concentration of small VLDL particles
Coronary artery disease and high-density lipoprotein cholesterol (bivariate)
Coronary artery disease and triglyceride (bivariate)
Distance between home and job workplace
Free cholesterol in medium VLDL
High cholesterol lipoprotein
High-density lipoprotein cholesterol
High-density-lipoprotein cholesterol
HMG CoA reductase inhibitors
Impedance measures - Impedance of leg (left)
Impedance measures - Impedance of leg (right)
Lifetime number of sexual partners
Number of treatments/medications taken
Phospholipids in medium VLDL
Total cholesterol in medium VLDL
Total lipids in medium VLDL
Triglyceride
Triglycerides cholesterol
Triglycerides in small VLDL
VLDL diameter
gwasStudies (18)
PMID24097068_chr8_rs12678919_Y2013_Metabolic_Triglycerides cholesterol
PMID27005778_chr8_rs115849089_Y2016_Metabolic_Cholesterol esters in medium VLDL
PMID27005778_chr8_rs1569209_Y2016_Metabolic_Cholesterol esters in large VLDL
PMID29403010_chr8_rs1059611_Y2018_Metabolic_Triglyceride
PMID30525989_chr8_rs17091891_Y2018_Cardiovascular_Coronary artery disease and triglyceride (bivariate)
PMID31427789_chr8_rs537562810_Y2019_Activities_Childhood sunburn occasions
PMID31427789_chr8_rs551916015_Y2019_Psychiatric_Average weekly intake of other alcoholic drinks
PMID24097068_chr8_rs13702_Y2013_Metabolic_High-density lipoprotein cholesterol
PMID30525989_chr8_rs17091891_Y2018_Cardiovascular_Coronary artery disease and high-density lipoprotein cholesterol (bivariate)
PMID27005778_chr8_rs115849089_Y2016_Metabolic_Concentration of medium VLDL particles
PMID27005778_chr8_rs115849089_Y2016_Metabolic_Concentration of small VLDL particles
PMID27005778_chr8_rs115849089_Y2016_Metabolic_Free cholesterol in medium VLDL
PMID27005778_chr8_rs115849089_Y2016_Metabolic_Phospholipids in medium VLDL
PMID27005778_chr8_rs115849089_Y2016_Metabolic_Total cholesterol in medium VLDL
PMID27005778_chr8_rs115849089_Y2016_Metabolic_Total lipids in medium VLDL
PMID27005778_chr8_rs115849089_Y2016_Metabolic_Triglycerides in small VLDL
PMID27005778_chr8_rs12679834_Y2016_Metabolic_VLDL diameter
PMID29403010_chr8_rs325_Y2018_Metabolic_High-density-lipoprotein cholesterol
PMID31427789_chr8_rs10101032_Y2019_Metabolic_Impedance measures - Impedance of leg (right)
PMID31427789_chr8_rs149742941_Y2019_Reproduction_Lifetime number of sexual partners
PMID31217584_chr8_rs10096633_Y2019_Metabolic_Triglyceride
PMID31427789_chr8_rs10101032_Y2019_Metabolic_Impedance measures - Impedance of leg (left)
PMID31427789_chr8_rs564368259_Y2019_Environment_Distance between home and job workplace
PMID19060906_chr8_rs12678919_Y2009_Metabolic_High-density lipoprotein cholesterol
PMID19060906_chr8_rs12678919_Y2009_Metabolic_Triglycerides cholesterol
PMID31015401_chr8_rs15285_Y2019_Environmental_HMG CoA reductase inhibitors
PMID31217584_chr8_rs13702_Y2019_Metabolic_High cholesterol lipoprotein
PMID31427789_chr8_rs11570891_Y2019_Activities_Number of treatments/medications taken
Domain
Year
Population
Trait

### Step_3: Refine the locus

As explained in Step_2, users can select the region based on either chr_id or a specific Gene name in any Chromosome to locate and refine the regional Manhattan plots.

After users refined/narrowed the region by using the top blue slider bar in the main Locus panel, more detailed plots will be displayed. Once the number of genes is reduced to 50 or less, both the names and the locations of genes will be displayed, as well as the detailed list of GWAS publications as shown earlier. Users can use options in Step_3 to Zoom In/Out the region/locus (see right).

Users can click "Zoom out (x10)" to increase the range of the existing region by 10 times. Note: It is important to select a final region that is less than 2Mb to run single and multi-trait fine-mapping efficiently in the next step.

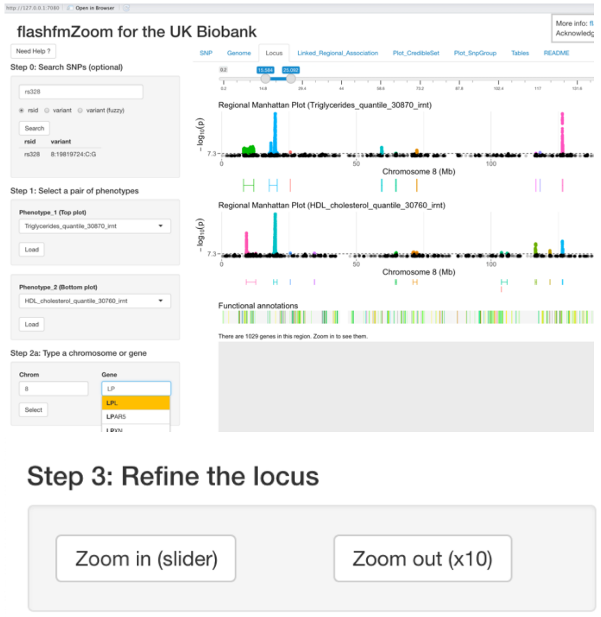


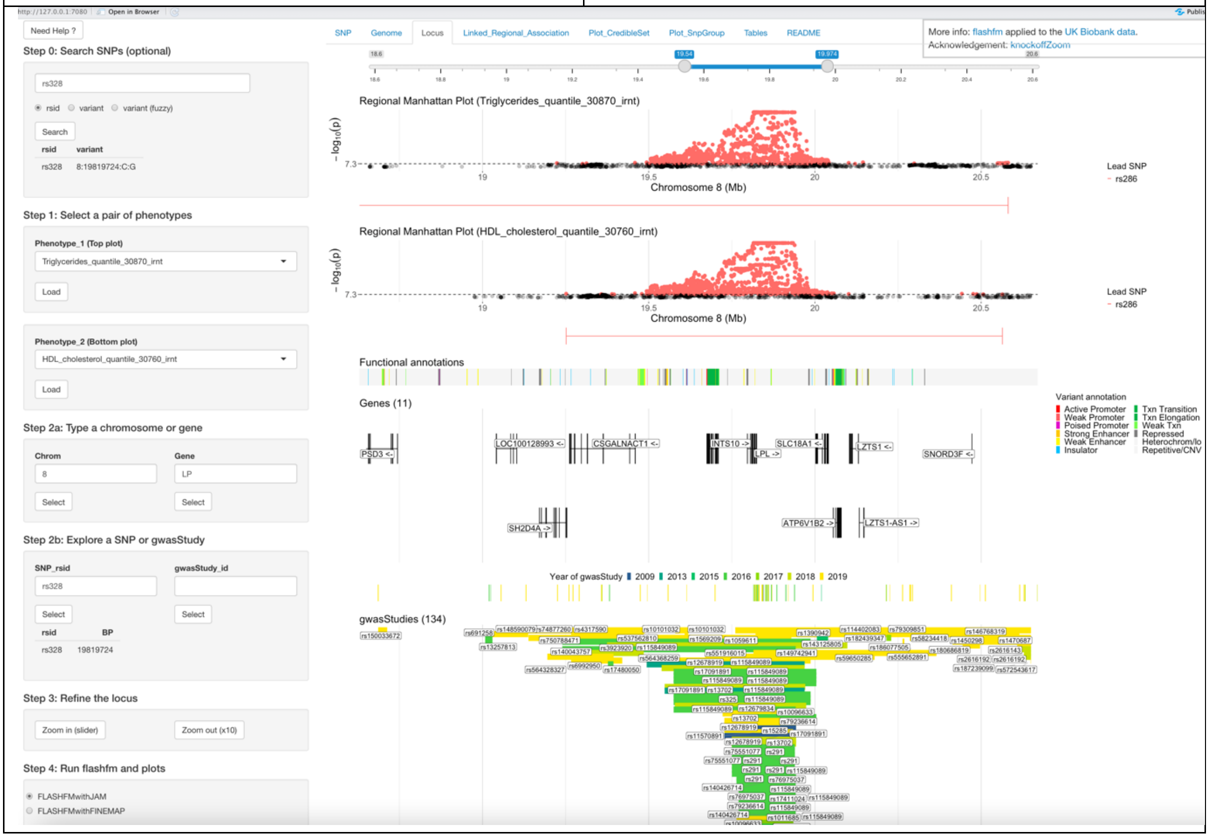

### Step_4: Run flashfm and plots

In addition to the information and instructions in **Table S1/S2/S3** earlier, if users click the "Need Help?", the relevant instructions will be displayed accordingly. Also, if users select a region that is larger than 2Mb (see an example below as the left cell) or the minimum Pval in one of traits is larger than 1e-5 (see the red coloured warning message as in the right cell below), the warning message will be displayed in red colour under the "Need Help?" button (see right). Once users selected a suitable region and moved to the "tab: Linked_Regional_Association" to prepare running flashfm, the "Select region (<2Mb)" button in Step_4 will be disabled to prevent users from wasting time to re-download a new LD matrix.

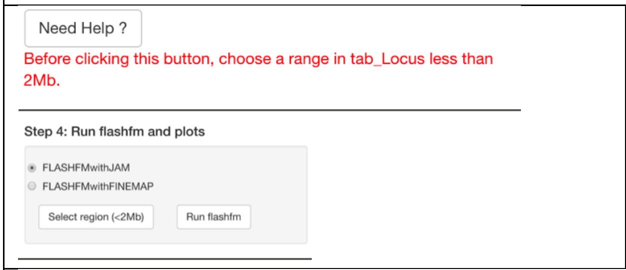


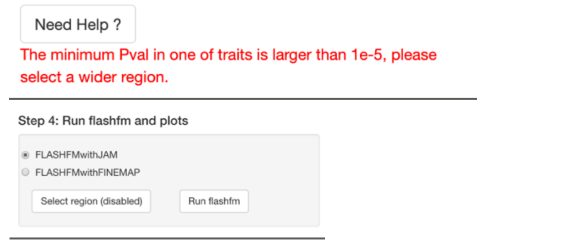


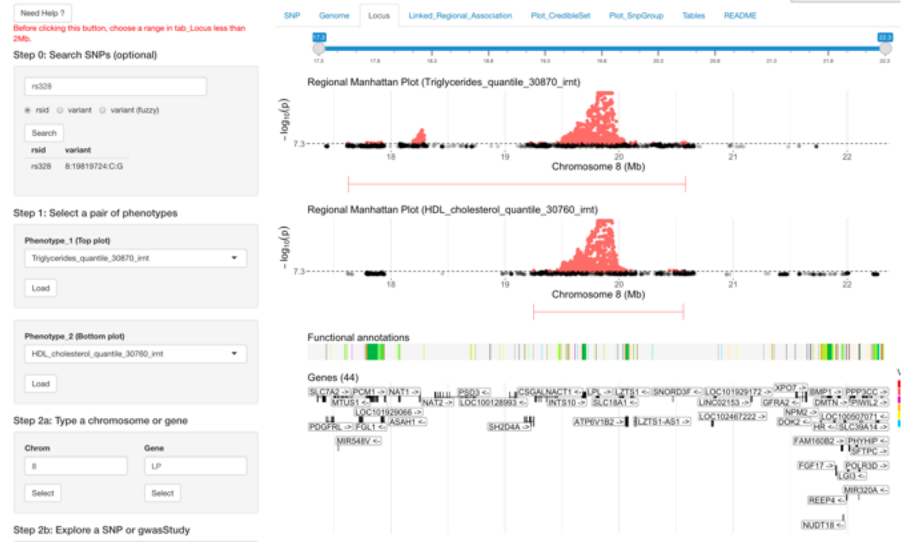


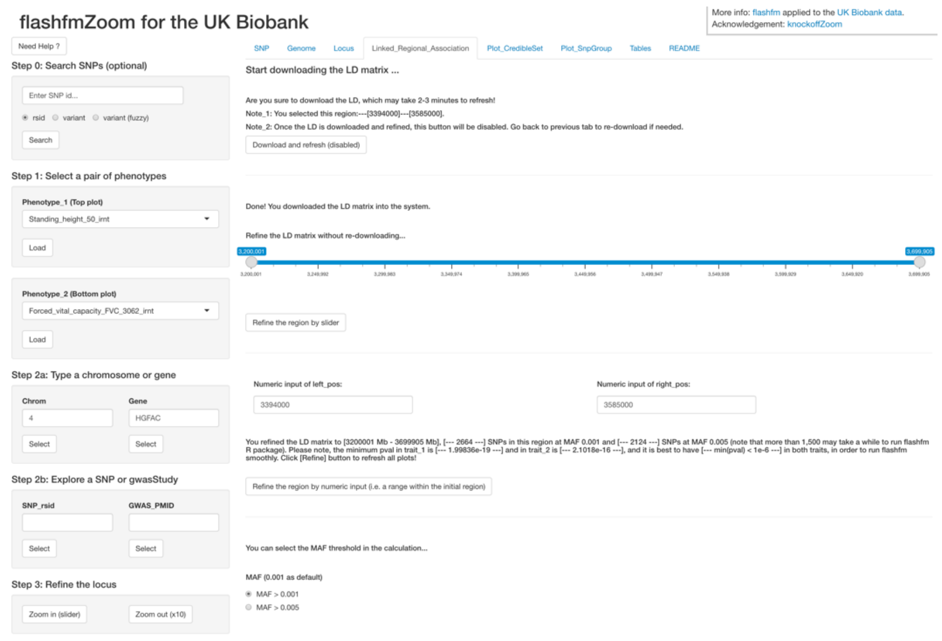

Users can double check the final selected region before downloading the corresponding LD matrix. After users downloaded the LD matrix, the correlations of both first and last 10 SNPs will be shown on the two heatmaps, in order for the users to check the LD matrix. Users can refine the range of this downloaded LD matrix by using the blue slider bar and click the “Refine” button. After they clicked the “Refine”, the “Download and refresh” button will be disabled. Users can also input numeric positions to refine the exact region and select a MAF threshold.

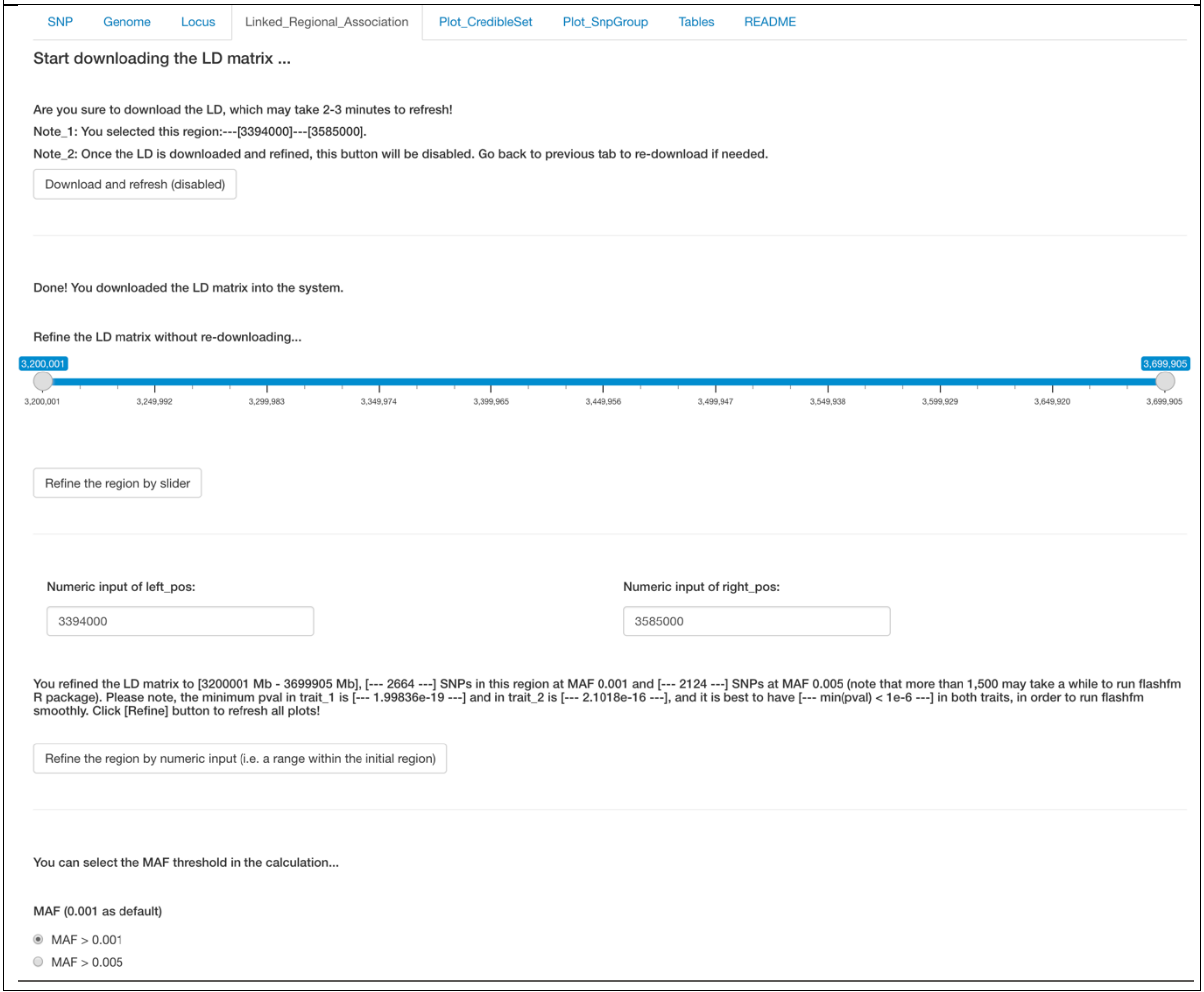

Since the flashfm package will be ran more efficiently when the number of SNPs is between 1000 and 2000, as well as there are at least one SNP with Pval<1e-6 in both traits, users can refine the region further by adjusting the blue slider bar. The text below the slider bar provides some useful information to the users.

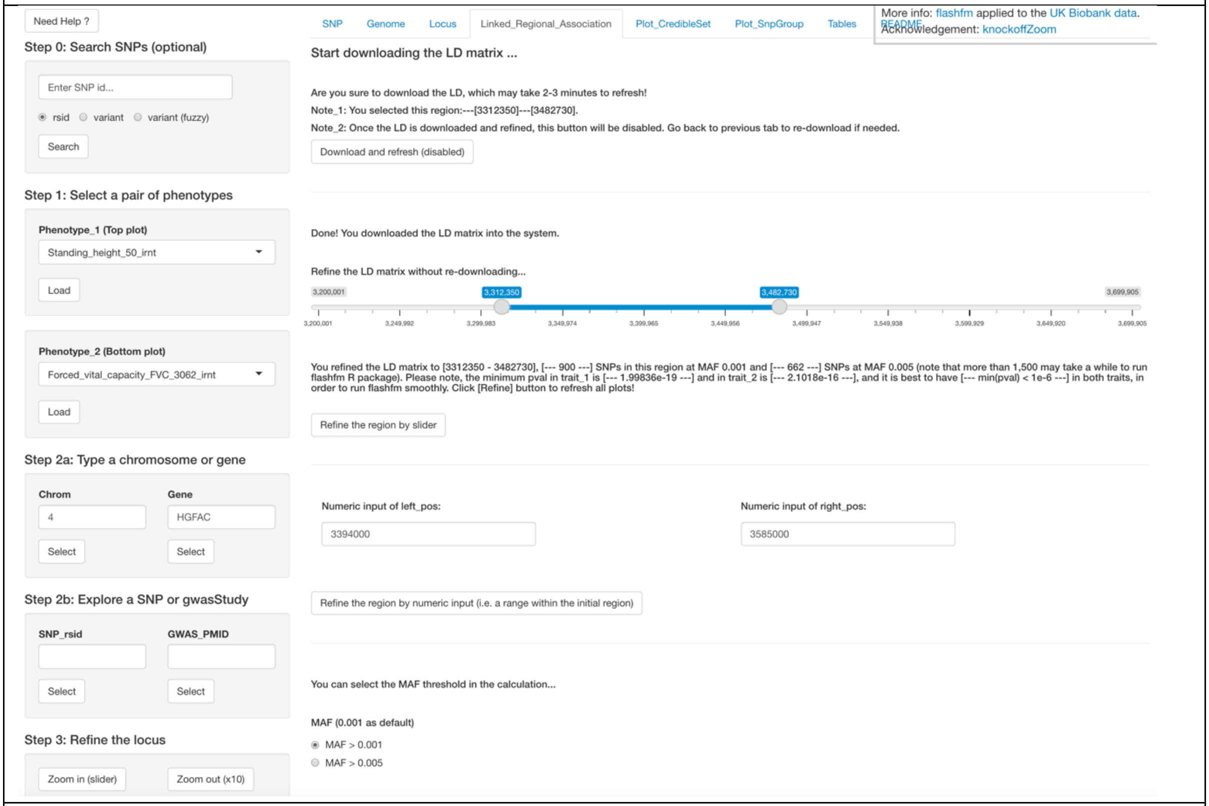


Once users are happy with the final region, they can select one of two flashfm methods/software and click "Run flashfm" button in left-side control widget of Step_4.

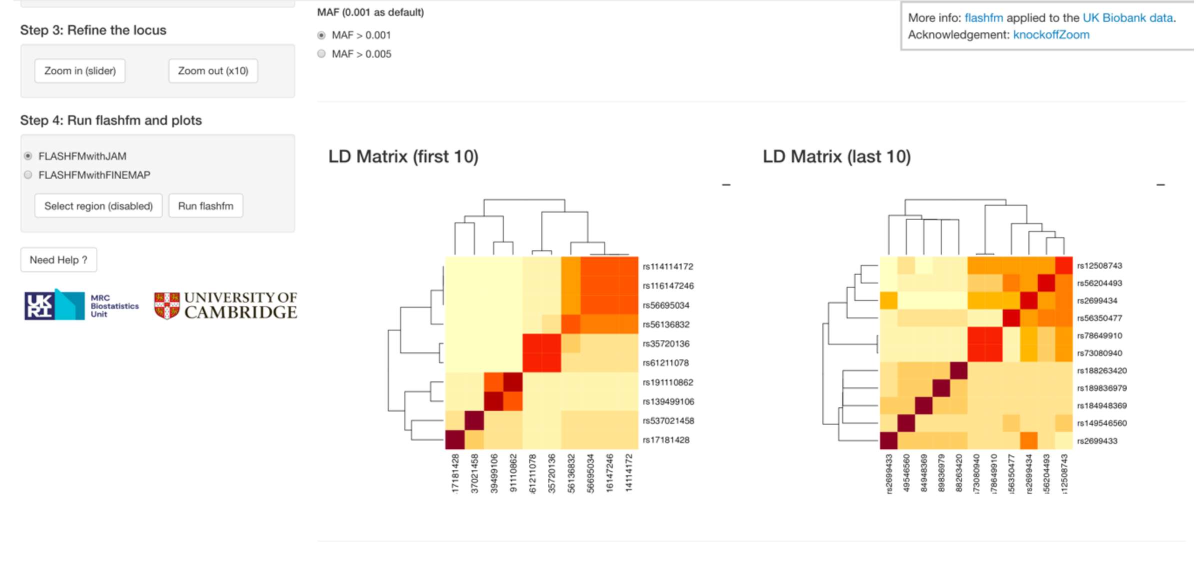

Users click the "Run flashfm" button in Step_4, then the outputs from flashfm package will be displayed. The detailed information about these plots is explained in next page.

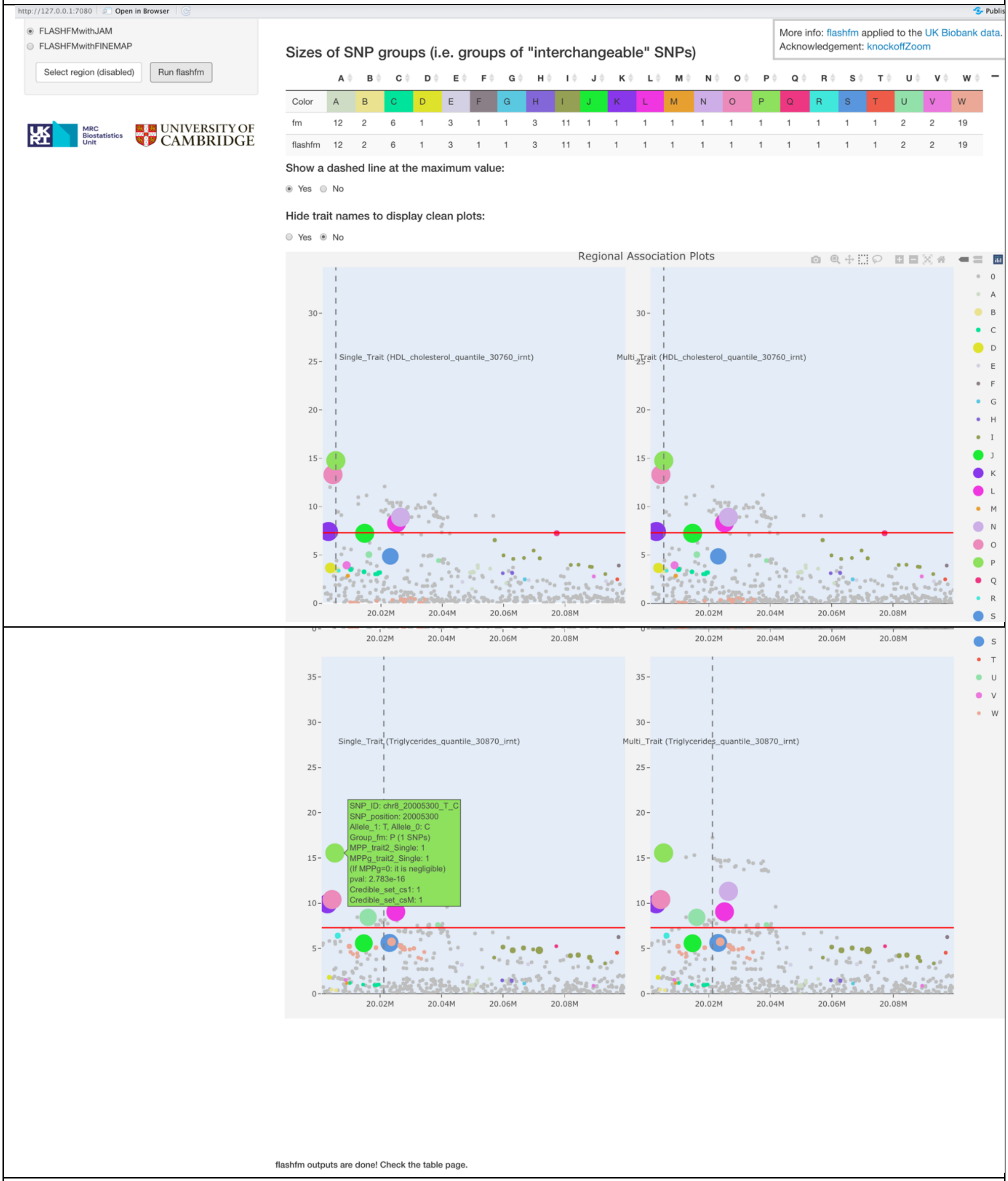


**Information about the plots:**

**Please note: For the terminology as mentioned here, please read the Section** "Terminology in the plots and the flashfm package (summary)" at the end of this Supplementary Materials.

**Coloured SNP Group– colours match those in the legend of the interactive regional association plots**

a) View SNP group sizes from both single and multi-trait fine-mapping
b) The table will be adjusted automatically depending on the computer screen size, but users can also use their mouse/touchpad to scroll left or right of the table.

**Coloured and linked regional association plots – view and interact with both GWAS and fine-mapping results**

The left panel is for single-trait fine-mapping results and the right panel is for multi-trait fine-mapping results. Each row shows individual trait results - a regional association plot ($-\log_{10}(p)$ against SNP position) with the following additional features:

a) Hover over a point to see SNP details (SNP ID, alleles, allele frequency, etc. Please note, if MPPg is negligible, we show it as equal to 0.) or click on “Compare data on hover” to see details for several SNPs nearby at once.

b) Colour of points: SNP group membership according to fine-mapping results. SNPs belonging to the same group can be viewed as exchangeable. SNPs with MPP>0.001 are assigned to the same group if they have high LD (pairwise $r^2$>0.6) and rarely appear in a model together (marginal posterior probability that a pair of SNPs are both included in a model < 0.01).

c) Size of points: proportional to fine-mapping posterior probabilities of SNP causality (referred to as MPP – Marginal Posterior Probability)

d) Click on “Box Select” or “Lasso Select” to draw a box or a lasso (free drawing of any shape) around points to focus on and fade other points. Automatically, this same set of points will become the focus in all other plots allowing simplified comparisons.

e) Click on “Zoom”, then draw a box around points to zoom in and out for point selection.

f) Double click on a SNP group in the legend to remove all points not belonging to that group. Double click again to show all points not belonging to the SNP group

g) Click on “Pan” and then drag the plot to the left or right to change the centre of the plot.

h) Click on reset axis to res-set to default plot

i) “Download plot as a PNG” option can download the plot to the local machine

j) “Autoscale” to adjust the view of the current plot

k) Users can change their internet browser’s size to adjust the view of the whole plot.

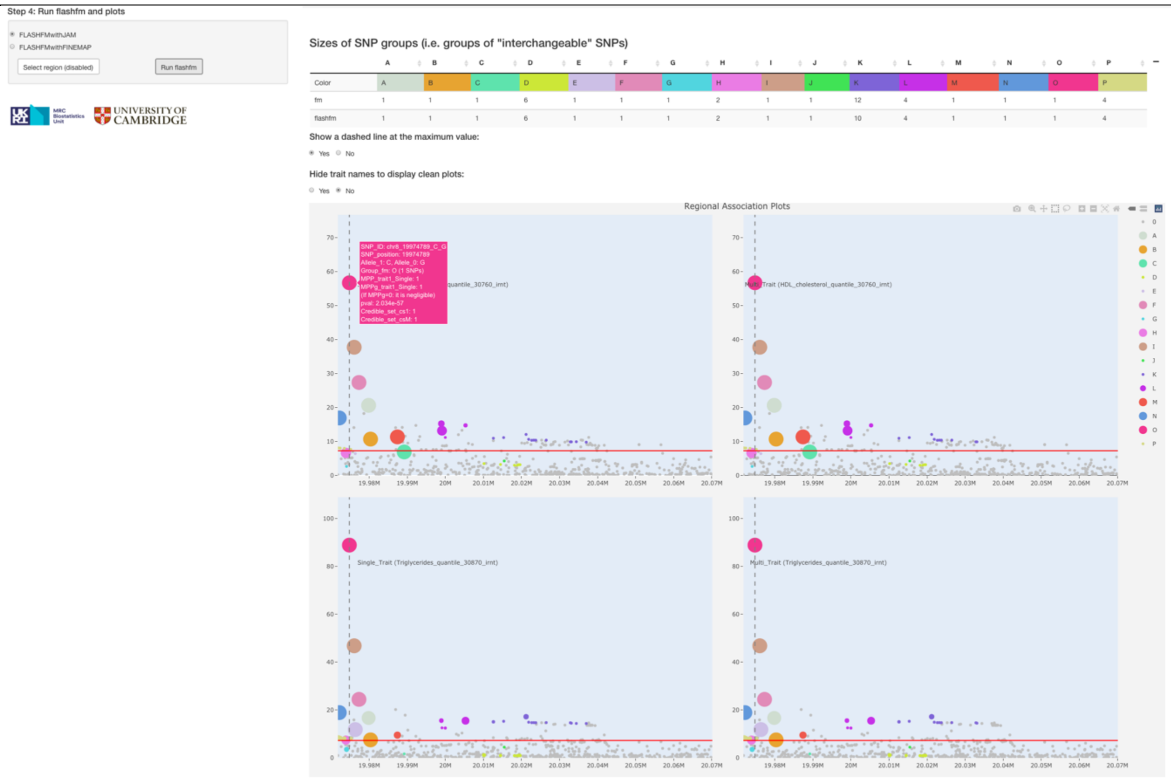

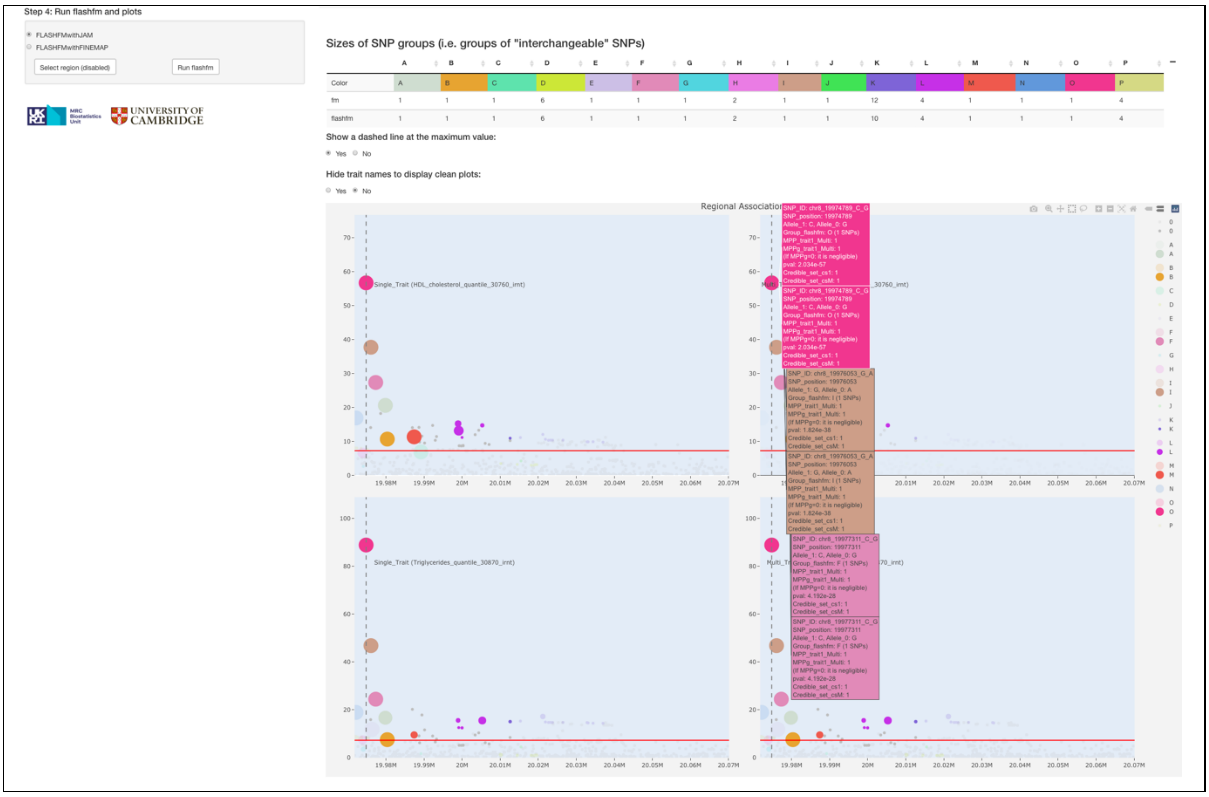

Step 4: Run flashfm and plots
FLASHFMwithJAM
FLASHFMwithFINEMAP
Select region (disabled)
Run flashfm
UNIVERSITY OF CAMBRIDGE
Sizes of SNP groups (i.e. groups of "interchangeable" SNPs)
Color: A B C D E F G H I J K L M N O P
fm: 1 1 1 6 1 1 1 2 1 1 12 4 1 1 1 4
flashfm: 1 1 1 6 1 1 1 2 1 1 10 4 1 1 1 4
Show a dashed line at the maximum value:
Yes No
Hide trait names to display clean plots:
Yes No
Regional Association
Single_Trait (HDL_cholesterol_quantile_30760_irnt)
Single_Trait (Triglycerides_quantile_30870_irnt)

Other plots in relation to Credible Sets and SNP Groups are displayed in the two tabs of the main panel: "Plot_CredibleSet" and "Plot_SnpGroup". Users can adjust the control widgets (i.e. the blue slider bars and radio buttons) to refine the interactive visualization. Both single- and multi-trait fine-mapping are displayed in separated plots, as well as individual results for the selected pair of traits.

More details about the following plots:

**Interactive regional association plots with users' defined credible sets**

a) The colour of the dots represents the SNPs in different credible sets
b) The size (i.e. diameter) of the dots defines the MPP values

**Area-proportional Venn diagrams - intuitive view of overlap between credible sets**

c) The segments show the number of SNPs that are shared between the intersecting credible sets
d) The area of each segment is proportional to its SNP count

**Interactive Venn diagram 1:CS_1 – overlap of fm credible sets between traits**

a) The segments show the number of SNPs that are shared between the intersecting credible sets
b) Hovering over a segment shows the SNP ids belonging to that intersection
c) The downloadable table shows the details of the credible set intersections, such as count and SNP ids. The columns can be sorted by clicking the top row options.

**Interactive Venn diagram 1:CS_M – overlap of flashfm credible sets between traits**

a) The segments show the number of SNPs that are shared between the intersecting credible sets
b) Hovering over a segment shows the SNP ids belonging to that intersection
**c)** The downloadable table shows the details of the credible set intersections, such as count and SNP ids. The columns can be sorted by clicking the top row options.

**Sankey diagrams of traits showing SNP group membership for each method**

a) Sankey diagram shows the SNPs that belong to each flashfm SNP group and to each fm SNP group
b) The flashfm groups tend to be subsets of the fm groups
c) There is an "All traits" tab and a tab for each trait
d) For a given fine-mapping method, SNP groups are the same for each trait
e) Each trait may have different SNP groups that appear in their models
f) To change the plot perspective (or show the names of SNPs clearly), the positions of the SNPs and groups may be moved by dragging them.
g) **All traits:** A combined Sankey diagram over all traits. The width of the lines joining the SNPs to their SNP group is proportional to the average MPP (over all traits, including some MPPs that are zero or very close to zero) for the SNP.
h) **Individual traits:** The width of the lines joining the SNPs to their SNP group is proportional to the trait-specific MPP for the SNP

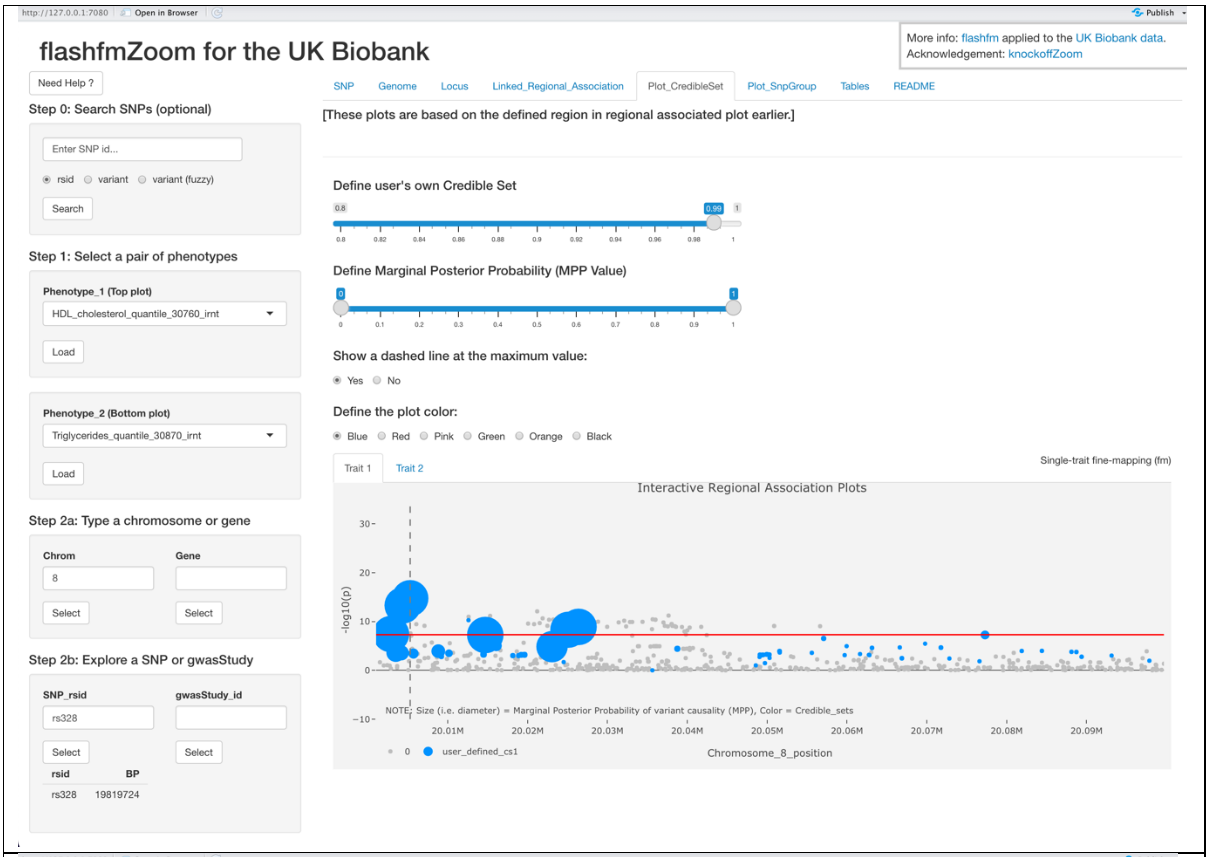
http://127.0.0.1:7080
Open in Browser
Publish
flashfmZoom for the UK Biobank
More info: flashfm applied to the UK Biobank data.
Acknowledgement: knockoffZoom
Need Help ?
Step 0: Search SNPs (optional)
Enter SNP id...
rsid
variant
variant (fuzzy)
Search
Step 1: Select a pair of phenotypes
Phenotype_1 (Top plot)
HDL_cholesterol_quantile_30760_irnt
Load
Phenotype_2 (Bottom plot)
Triglycerides_quantile_30870_irnt
Load
Step 2a: Type a chromosome or gene
Chrom
Gene
8
Select
Select
Step 2b: Explore a SNP or gwasStudy
SNP_rsid
gwasStudy_id
rs328
Select
Select
rsid
BP
rs328
19819724
SNP
Genome
Locus
Linked_Regional_Association
Plot_CredibleSet
Plot_SnpGroup
Tables
README
[These plots are based on the defined region in regional associated plot earlier.]
Define user's own Credible Set
Define Marginal Posterior Probability (MPP Value)
Show a dashed line at the maximum value:
Yes
No
Define the plot color:
Blue
Red
Pink
Green
Orange
Black
Trait 1
Trait 2
Single-trait fine-mapping (fm)
Interactive Regional Association Plots
-log10(p)
NOTE: Size (i.e. diameter) = Marginal Posterior Probability of variant causality (MPP), Color = Credible_sets
20.01M
20.02M
20.03M
20.04M
20.05M
20.06M
20.07M
20.08M
20.09M
0
user_defined_cs1
Chromosome_8_position

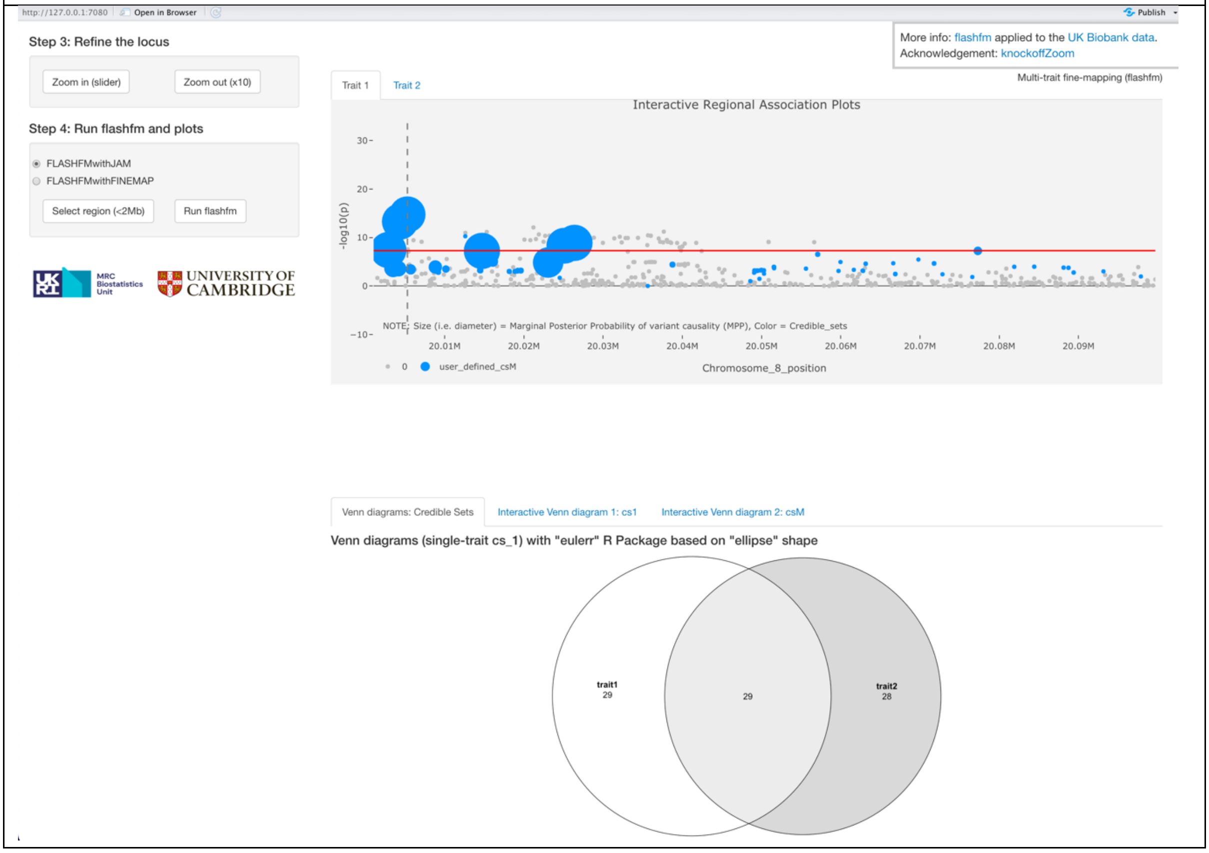
http://127.0.0.1:7080
Open in Browser
Publish
Step 3: Refine the locus
Zoom in (slider)
Zoom out (x10)
Step 4: Run flashfm and plots
FLASHFMwithJAM
FLASHFMwithFINEMAP
Select region (<2Mb)
Run flashfm
MRC Biostatistics Unit
UNIVERSITY OF CAMBRIDGE
More info: flashfm applied to the UK Biobank data.
Acknowledgement: knockoffZoom
Trait 1
Trait 2
Multi-trait fine-mapping (flashfm)
Interactive Regional Association Plots
-log10(p)
NOTE: Size (i.e. diameter) = Marginal Posterior Probability of variant causality (MPP), Color = Credible_sets
20.01M
20.02M
20.03M
20.04M
20.05M
20.06M
20.07M
20.08M
20.09M
0
user_defined_csM
Chromosome_8_position
Venn diagrams: Credible Sets
Interactive Venn diagram 1: cs1
Interactive Venn diagram 2: csM
Venn diagrams (single-trait cs_1) with "eulerr" R Package based on "ellipse" shape
trait1
29
29
trait2
28

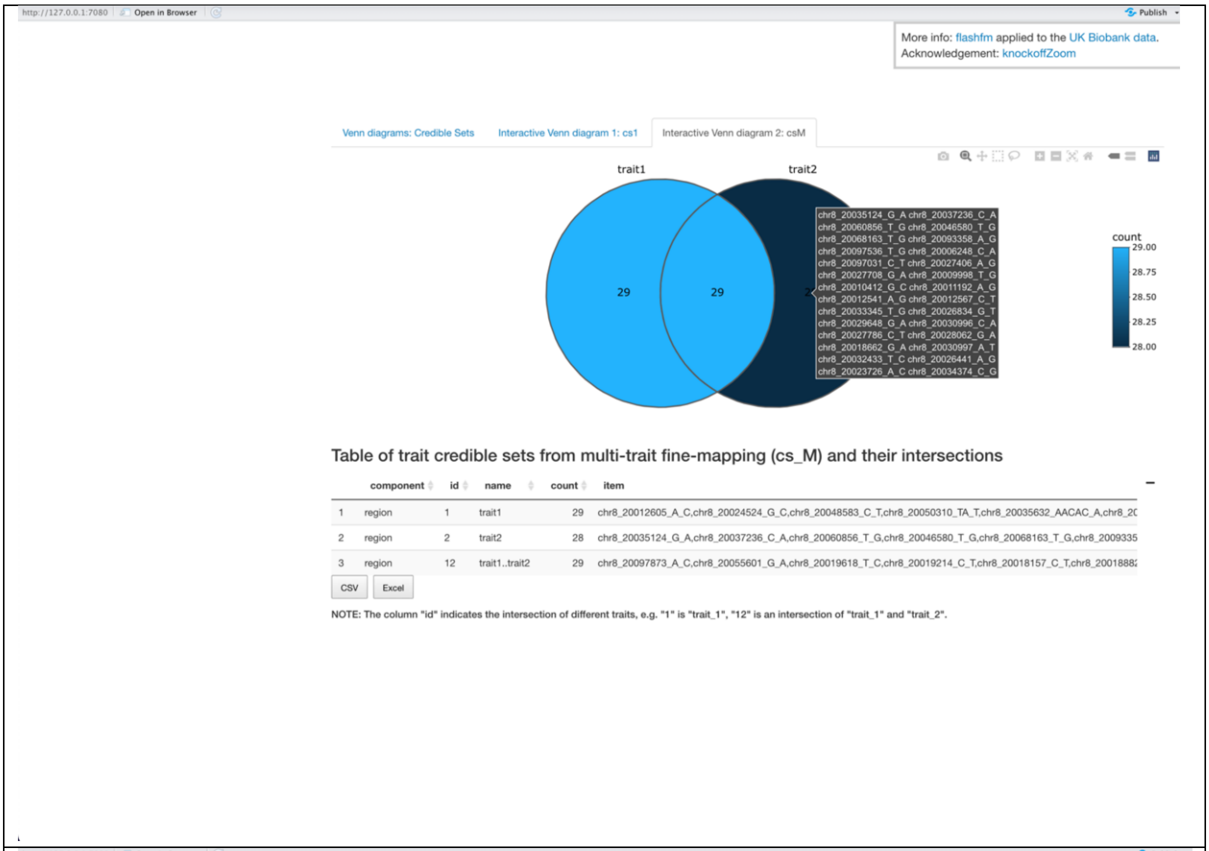
More info: flashfm applied to the UK Biobank data.
Acknowledgement: knockoffZoom
Venn diagrams: Credible Sets
Interactive Venn diagram 1: cs1
Interactive Venn diagram 2: csM
trait1
trait2
29
29
count
29.00
28.75
28.50
28.25
28.00
Table of trait credible sets from multi-trait fine-mapping (cs_M) and their intersections
component
id
name
count
item
1 region 1 trait1 29
2 region 2 trait2 28
3 region 12 trait1..trait2 29
CSV
Excel
NOTE: The column "id" indicates the intersection of different traits, e.g. "1" is "trait_1", "12" is an intersection of "trait_1" and "trait_2".

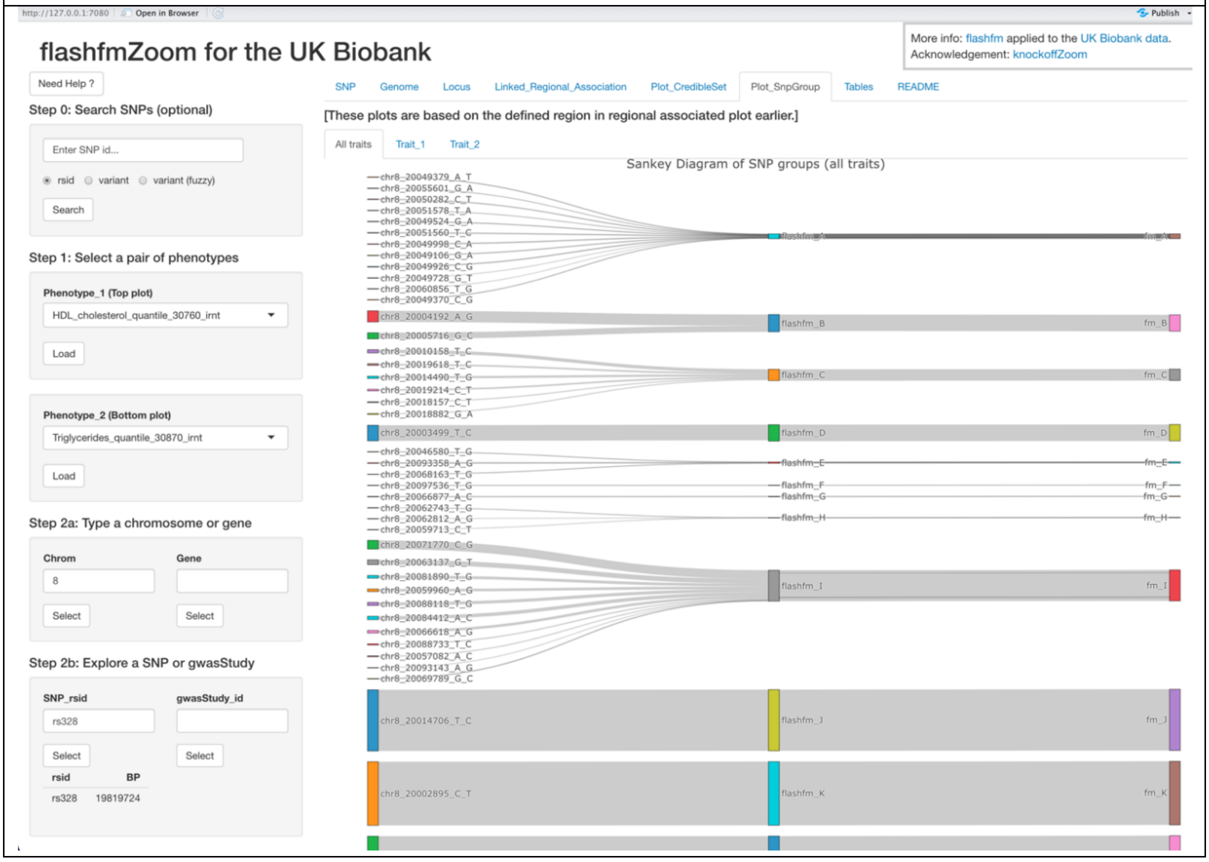
flashfmZoom for the UK Biobank
More info: flashfm applied to the UK Biobank data.
Acknowledgement: knockoffZoom
Need Help ?
Step 0: Search SNPs (optional)
Enter SNP id...
rsid
variant
variant (fuzzy)
Search
Step 1: Select a pair of phenotypes
Phenotype_1 (Top plot)
HDL_cholesterol_quantile_30760_irnt
Load
Phenotype_2 (Bottom plot)
Triglycerides_quantile_30870_irnt
Load
Step 2a: Type a chromosome or gene
Chrom
Gene
8
Select
Select
Step 2b: Explore a SNP or gwasStudy
SNP_rsid
gwasStudy_id
rs328
Select
Select
rsid
BP
rs328
19819724
SNP
Genome
Locus
Linked_Regional_Association
Plot_CredibleSet
Plot_SnpGroup
Tables
README
[These plots are based on the defined region in regional associated plot earlier.]
All traits
Trait_1
Trait_2
Sankey Diagram of SNP groups (all traits)
flashfm_B
flashfm_C
flashfm_D
flashfm_E
flashfm_F
flashfm_G
flashfm_H
flashfm_I
flashfm_J
flashfm_K
fm_B
fm_C
fm_D
fm_E
fm_F
fm_G
fm_H
fm_I
fm_J
fm_K
chr8_20014706_T_C
chr8_20002895_C_T

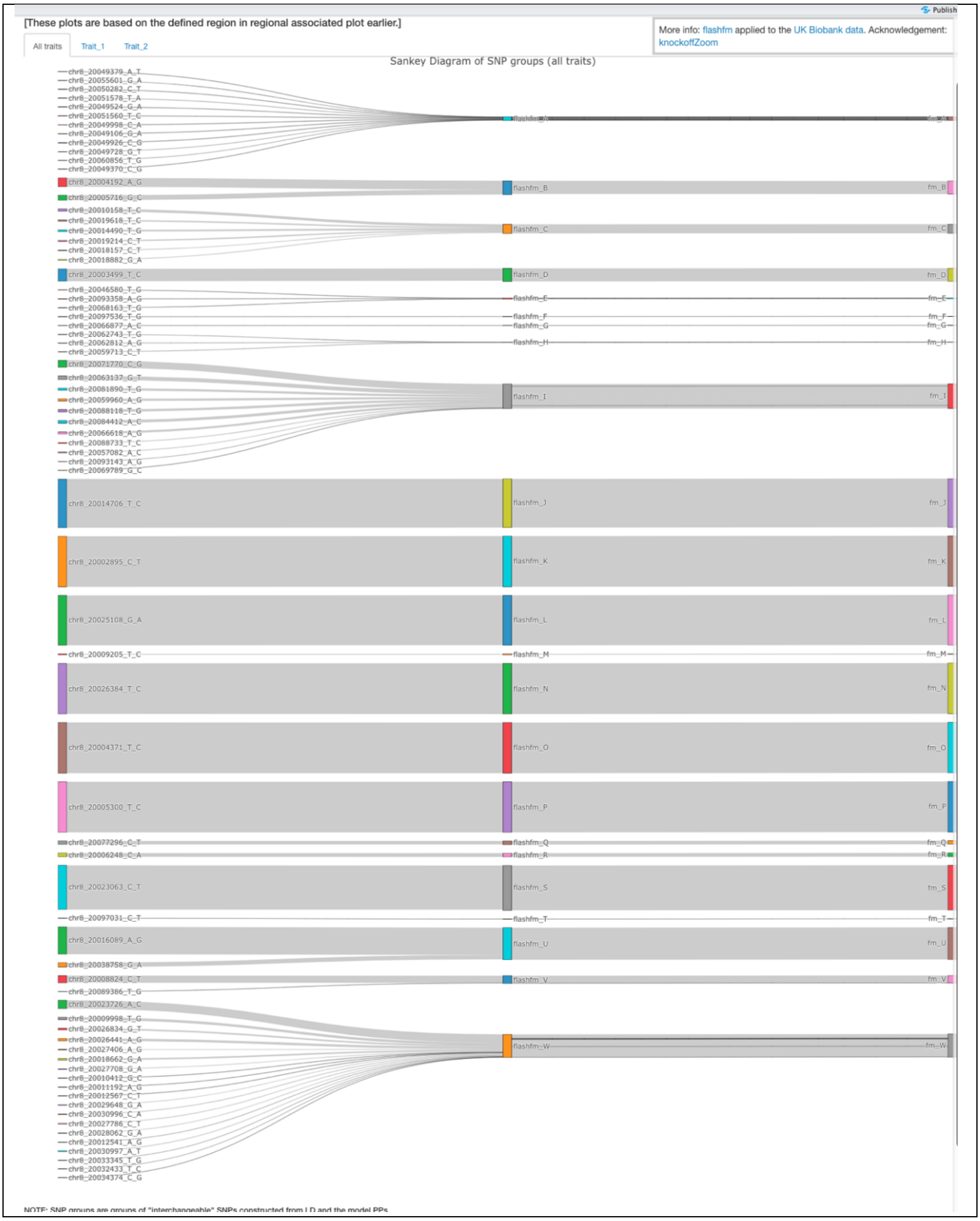
[These plots are based on the defined region in regional associated plot earlier.]
More info: flashfm applied to the UK Biobank data. Acknowledgement: knockoffZoom
Publish
All traits
Trait_1
Trait_2
Sankey Diagram of SNP groups (all traits)
chr8_20049379_A_T
chr8_20055601_G_A
chr8_20050282_C_T
chr8_20051578_T_A
chr8_20049524_G_A
chr8_20051560_T_C
chr8_20049998_C_A
chr8_20049106_G_A
chr8_20049926_C_G
chr8_20049728_G_T
chr8_20060856_T_G
chr8_20049370_C_G
chr8_20004192_A_G
chr8_20005716_G_C
chr8_20010158_T_C
chr8_20019618_T_C
chr8_20014490_T_G
chr8_20019214_C_T
chr8_20018157_C_T
chr8_20018882_G_A
chr8_20003499_T_C
chr8_20046580_T_G
chr8_20093358_A_G
chr8_20068163_T_G
chr8_20097536_T_G
chr8_20066877_A_C
chr8_20062743_T_G
chr8_20062812_A_G
chr8_20059713_C_T
chr8_20071770_C_G
chr8_20063137_G_T
chr8_20081890_T_G
chr8_20059960_A_G
chr8_20088118_T_G
chr8_20084412_A_C
chr8_20066618_A_G
chr8_20088733_T_C
chr8_20057082_A_C
chr8_20093143_A_G
chr8_20069789_G_C
chr8_20014706_T_C
chr8_20002895_C_T
chr8_20025108_G_A
chr8_20009205_T_C
chr8_20026384_T_C
chr8_20004371_T_C
chr8_20005300_T_C
chr8_20077296_C_T
chr8_20006248_C_A
chr8_20023063_C_T
chr8_20097031_C_T
chr8_20016089_A_G
chr8_20038758_G_A
chr8_20008824_C_T
chr8_20089386_T_G
chr8_20023726_A_C
chr8_20009998_T_G
chr8_20026834_G_T
chr8_20026441_A_G
chr8_20027406_A_G
chr8_20018662_G_A
chr8_20027708_G_A
chr8_20010412_G_C
chr8_20011192_A_G
chr8_20012567_C_T
chr8_20029648_G_A
chr8_20030996_C_A
chr8_20027786_C_T
chr8_20028062_G_A
chr8_20012541_A_G
chr8_20030997_A_T
chr8_20033345_T_G
chr8_20032433_T_C
chr8_20034374_C_G
flashfm_A
flashfm_B
flashfm_C
flashfm_D
flashfm_E
flashfm_F
flashfm_G
flashfm_H
flashfm_I
flashfm_J
flashfm_K
flashfm_L
flashfm_M
flashfm_N
flashfm_O
flashfm_P
flashfm_Q
flashfm_R
flashfm_S
flashfm_T
flashfm_U
flashfm_V
flashfm_W
fm_A
fm_B
fm_C
fm_D
fm_E
fm_F
fm_G
fm_H
fm_I
fm_J
fm_K
fm_L
fm_M
fm_N
fm_O
fm_P
fm_Q
fm_R
fm_S
fm_T
fm_U
fm_V
fm_W
NOTE: SNP groups are groups of "interchangeable" SNPs constructed from LD and the model PPs

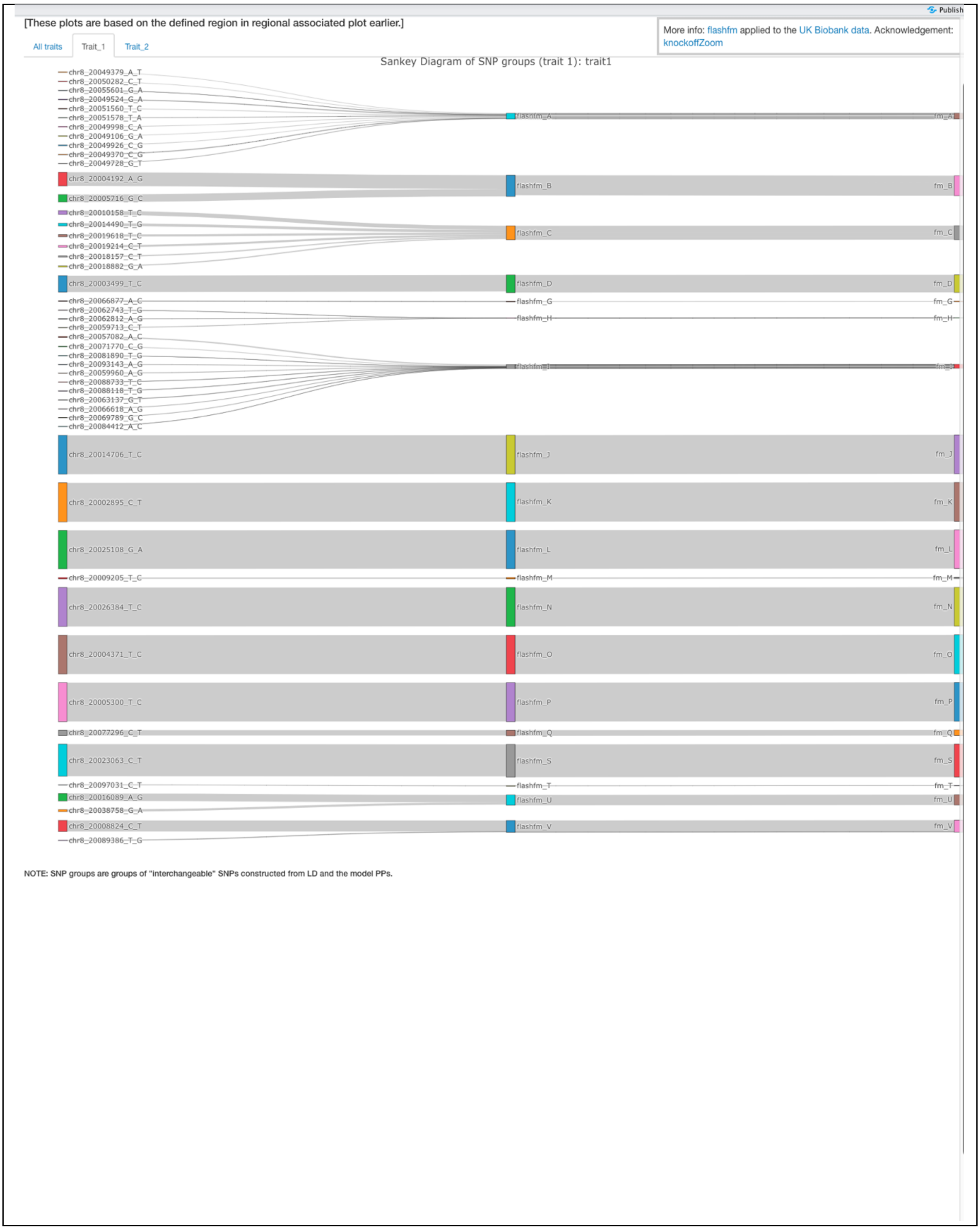
Publish
[These plots are based on the defined region in regional associated plot earlier.]
More info: flashfm applied to the UK Biobank data. Acknowledgement: knockoffZoom
All traits
Trait_1
Trait_2
Sankey Diagram of SNP groups (trait 1): trait1
chr8_20049379_A_T
chr8_20050282_C_T
chr8_20055601_G_A
chr8_20049524_G_A
chr8_20051560_T_C
chr8_20051578_T_A
chr8_20049998_C_A
chr8_20049106_G_A
chr8_20049926_C_G
chr8_20049370_C_G
chr8_20049728_G_T
flashfm_A
fm_A
chr8_20004192_A_G
chr8_20005716_G_C
flashfm_B
fm_B
chr8_20010158_T_C
chr8_20014490_T_G
chr8_20019618_T_C
chr8_20019214_C_T
chr8_20018157_C_T
chr8_20018882_G_A
flashfm_C
fm_C
chr8_20003499_T_C
flashfm_D
fm_D
chr8_20066877_A_C
chr8_20062743_T_G
chr8_20062812_A_G
chr8_20059713_C_T
chr8_20057082_A_C
chr8_20071770_C_G
chr8_20081890_T_G
chr8_20093143_A_G
chr8_20059960_A_G
chr8_20088733_T_C
chr8_20088118_T_G
chr8_20063137_G_T
chr8_20066618_A_G
chr8_20069789_G_C
chr8_20084412_A_C
flashfm_G
fm_G
flashfm_H
fm_H
flashfm_I
fm_I
chr8_20014706_T_C
flashfm_J
fm_J
chr8_20002895_C_T
flashfm_K
fm_K
chr8_20025108_G_A
flashfm_L
fm_L
chr8_20009205_T_C
flashfm_M
fm_M
chr8_20026384_T_C
flashfm_N
fm_N
chr8_20004371_T_C
flashfm_O
fm_O
chr8_20005300_T_C
flashfm_P
fm_P
chr8_20077296_C_T
flashfm_Q
fm_Q
chr8_20023063_C_T
flashfm_S
fm_S
chr8_20097031_C_T
flashfm_T
fm_T
chr8_20016089_A_G
chr8_20038758_G_A
flashfm_U
fm_U
chr8_20008824_C_T
chr8_20089386_T_G
flashfm_V
fm_V
NOTE: SNP groups are groups of "interchangeable" SNPs constructed from LD and the model PPs.

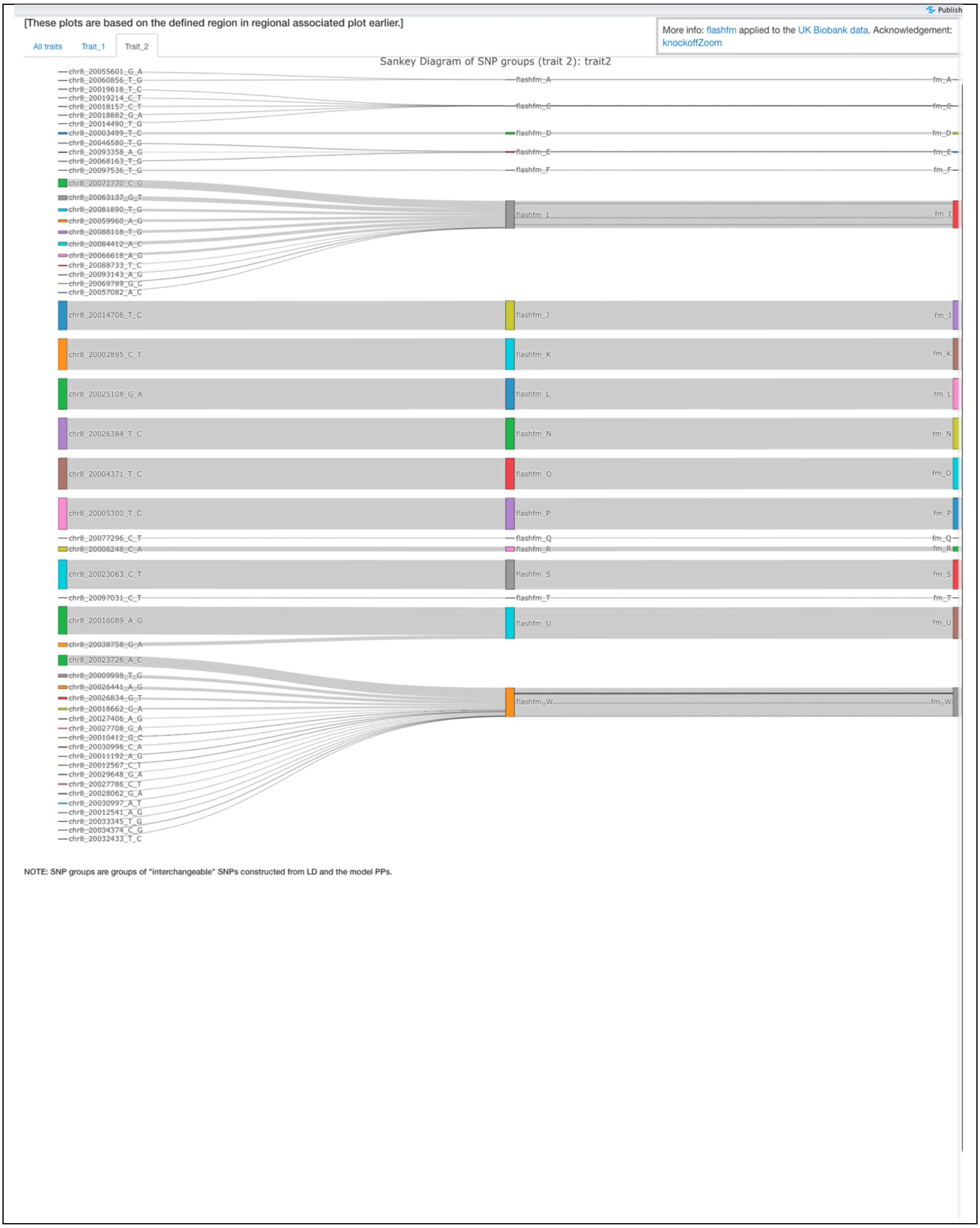
Publish
[These plots are based on the defined region in regional associated plot earlier.]
More info: flashfm applied to the UK Biobank data. Acknowledgement: knockoffZoom
All traits
Trait_1
Trait_2
Sankey Diagram of SNP groups (trait 2): trait2
chr8_20055601_G_A
chr8_20060856_T_G
chr8_20019618_T_C
chr8_20019214_C_T
chr8_20018157_C_T
chr8_20018882_G_A
chr8_20014490_T_G
chr8_20003499_T_C
chr8_20046580_T_G
chr8_20093358_A_G
chr8_20068163_T_G
chr8_20097536_T_G
chr8_20071770_C_G
chr8_20063137_G_T
chr8_20081890_T_G
chr8_20059960_A_G
chr8_20088118_T_G
chr8_20084412_A_C
chr8_20066618_A_G
chr8_20088733_T_C
chr8_20093143_A_G
chr8_20069789_G_C
chr8_20057082_A_C
chr8_20014706_T_C
chr8_20002895_C_T
chr8_20025108_G_A
chr8_20026384_T_C
chr8_20004371_T_C
chr8_20005300_T_C
chr8_20077296_C_T
chr8_20006248_C_A
chr8_20023063_C_T
chr8_20097031_C_T
chr8_20016089_A_G
chr8_20038758_G_A
chr8_20023726_A_C
chr8_20009998_T_G
chr8_20026441_A_G
chr8_20026834_G_T
chr8_20018662_G_A
chr8_20027406_A_G
chr8_20027708_G_A
chr8_20010412_G_C
chr8_20030996_C_A
chr8_20011192_A_G
chr8_20012567_C_T
chr8_20029648_G_A
chr8_20027786_C_T
chr8_20028062_G_A
chr8_20030997_A_T
chr8_20012541_A_G
chr8_20033345_T_G
chr8_20034374_C_G
chr8_20032433_T_C
flashfm_A
flashfm_C
flashfm_D
flashfm_E
flashfm_F
flashfm_I
flashfm_J
flashfm_K
flashfm_L
flashfm_N
flashfm_O
flashfm_P
flashfm_Q
flashfm_R
flashfm_S
flashfm_T
flashfm_U
flashfm_W
fm_A
fm_C
fm_D
fm_E
fm_F
fm_I
fm_J
fm_K
fm_L
fm_N
fm_O
fm_P
fm_Q
fm_R
fm_S
fm_T
fm_U
fm_W
NOTE: SNP groups are groups of "interchangeable" SNPs constructed from LD and the model PPs.

### Download results and tables

All information and outputs that are used and displayed in the plots can be downloaded easily, in order to be more transparent for users to check the results. The full table of GWAS association results (including SNP groups and credible sets for both traits from different fine-mapping methods) can be downloaded as well, so users can do further analyses based on this table in their local machines.

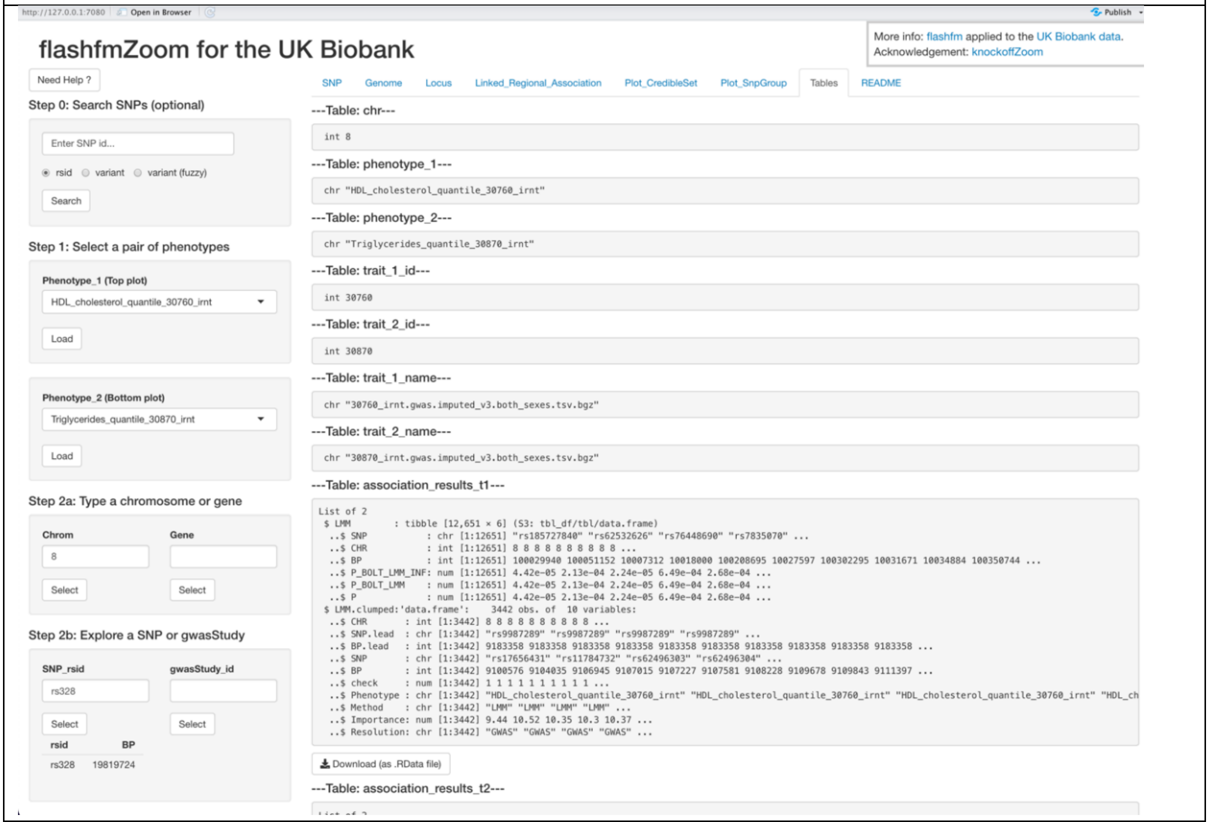


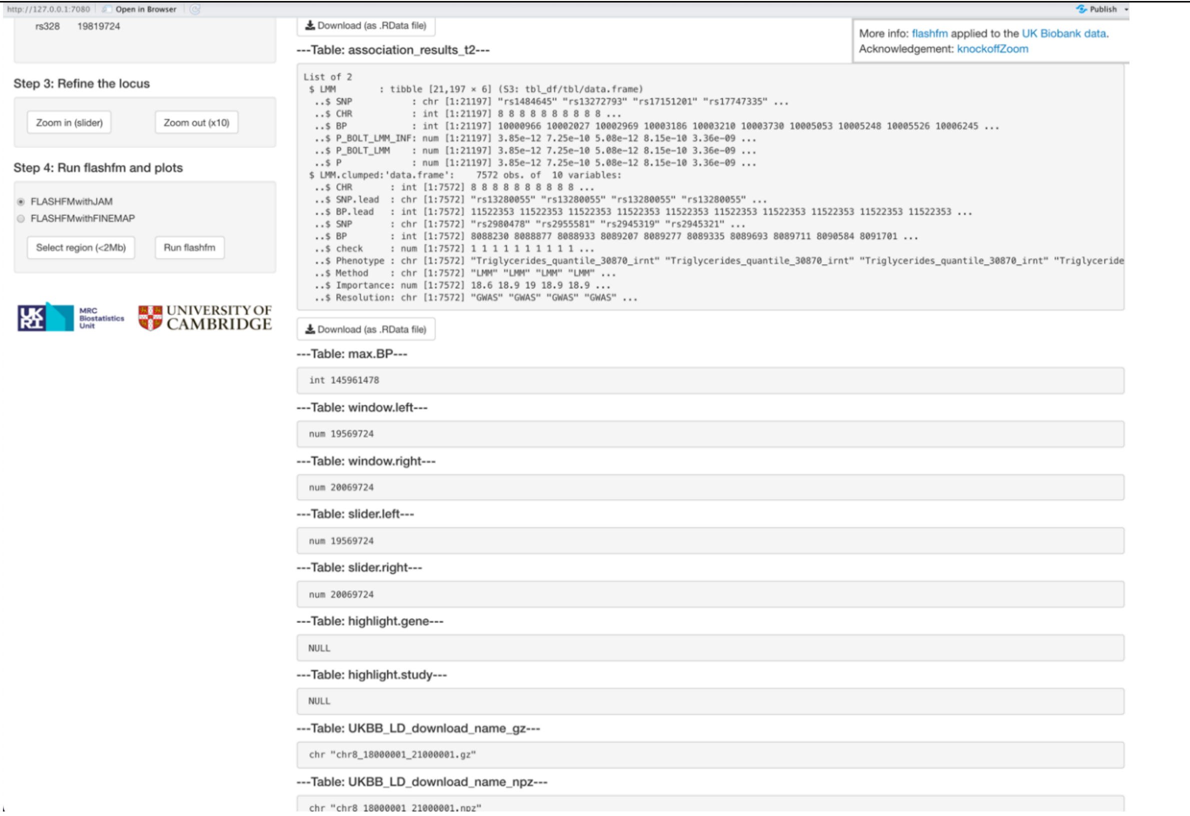

http://127.0.0.1:7080 | Open in Browser | Publish

```
chr "chr8_18000001_21000001.gz"
```

More info: flashfm applied to the UK Biobank data.
Acknowledgement: knockoffZoom

---Table: UKBB_LD_download_name_npz---

```
chr "chr8_18000001_21000001.npz"
```

---Table: state$flashfm_sel.left---

```
int 20000972
```

---Table: state$flashfm_sel.right---

```
int 20099697
```

---Table: state$fm_LD_matrix---

```
num [1:544, 1:544] 1 1 1 -0.0327 0.419 ...
- attr(*, "dimnames")=List of 2
 ..$ : chr [1:544] "8:20001033:C:A" "8:20001324:G:A" "8:20001454:C:T" "8:20001513:AC:A" ...
 ..$ : chr [1:544] "8:20001033:C:A" "8:20001324:G:A" "8:20001454:C:T" "8:20001513:AC:A" ...
```

Download (as .RData file)

---Table: state$fm_LD_variant---

```
'data.frame':   544 obs. of  6 variables:
 $ rsid      : chr  "rs13268842" "rs11204096" "rs11204097" "8:20001513_AC_A" ...
 $ chromosome: int  8 8 8 8 8 8 8 8 8 8 ...
 $ position  : int  20001033 20001324 20001454 20001513 20001995 20002426 20002739 20002888 20002895 20003431 ...
 $ allele1   : chr  "C" "G" "C" "AC" ...
 $ allele2   : chr  "A" "A" "T" "A" ...
 $ variant   : chr  "8:20001033:C:A" "8:20001324:G:A" "8:20001454:C:T" "8:20001513:AC:A" ...
```

Download (as .RData file)

---Table: state$fm_outputs_all---

```
List of 8
 $ fm_cs1      :List of 2
  ..$ : chr [1:58] "chr8_20004192_A_G" "chr8_20002895_C_T" "chr8_20004371_T_C" "chr8_20005300_T_C" ...
  ..$ : chr [1:57] "chr8_20002895_C_T" "chr8_20004371_T_C" "chr8_20005300_T_C" "chr8_20014706_T_C" ...
 $ fm_csM      :List of 2
  ..$ : chr [1:58] "chr8_20004192_A_G" "chr8_20002895_C_T" "chr8_20004371_T_C" "chr8_20005300_T_C" ...
  ..$ : chr [1:57] "chr8_20002895_C_T" "chr8_20004371_T_C" "chr8_20005300_T_C" "chr8_20014706_T_C" ...
 $ fm_LD       : num [1:544, 1:544] 1 1 1 -0.0327 0.419 ...
  ..- attr(*, "dimnames")=List of 2
  .. ..$ : chr [1:544] "chr8_20001033_C_A" "chr8_20001324_G_A" "chr8_20001454_C_T" "chr8_20001513_AC_A" ...
  .. ..$ : chr [1:544] "chr8_20001033_C_A" "chr8_20001324_G_A" "chr8_20001454_C_T" "chr8_20001513_AC_A" ...
 $ fm_GWAS     :List of 2
  ..$ trait_1:'data.frame':     544 obs. of  11 variables:
  .. ..$ variant   : chr [1:544] "chr8_20001033_C_A" "chr8_20001324_G_A" "chr8_20001454_C_T" "chr8_20001513_AC_A" ...
  .. ..$ rsid      : chr [1:544] "rs13268842" "rs11204096" "rs11204097" "8:20001513_AC_A" ...
  .. ..$ chr       : chr [1:544] "8" "8" "8" "8" ...
  .. ..$ pos       : num [1:544] 2e+07 2e+07 2e+07 2e+07 2e+07 ...
  .. ..$ ref       : chr [1:544] "C" "G" "C" "AC" ...
  .. ..$ alt       : chr [1:544] "A" "A" "T" "A" ...
  .. ..$ beta      : num [1:544] -0.01378 -0.01375 -0.01393 0.05942 -0.00401 ...
  .. ..$ se        : num [1:544] 0.00255 0.00255 0.00255 0.02594 0.00244 ...
  .. ..$ pval      : num [1:544] 6.82e-08 7.25e-08 4.87e-08 2.20e-02 1.01e-01 ...
  .. ..$ MAF_trait : num [1:544] 0.27373 0.27371 0.27353 0.00229 0.31811 ...
  .. ..$ no_trait_1: num [1:544] 381439 381439 381439 381439 381439 ...
  ..$ trait_2:'data.frame':     544 obs. of  11 variables:
```

http://127.0.0.1:7080 | Open in Browser | Publish

```
[illegible]
  ..- attr(*, "names")= chr [1:544] "chr8_20001033_C_A" "chr8_20001324_G_A" "chr8_20001454_C_T" "chr8_20001513_AC_A" ...
```

Acknowledgement: knockoffZoom

Download (as .RData file)

--- GWAS summary statistics---

Show 10 entries

Search:

| | snpGroups_flashfm | snpGroups_fm | variant | rsid | chr | pos | ref | alt | beta_1 | se_1 | pval_1 | MAF_A |
|---|---|---|---|---|---|---|---|---|---|---|---|---|
| 1 | 0 | 0 | chr8_20001033_C_A | rs13268842 | 8 | 20001033 | C | A | -0.013782 | 0.002554 | 6.8165e-8 | |
| 2 | 0 | 0 | chr8_20001324_G_A | rs11204096 | 8 | 20001324 | G | A | -0.013749 | 0.0025532 | 7.2466e-8 | |
| 3 | 0 | 0 | chr8_20001454_C_T | rs11204097 | 8 | 20001454 | C | T | -0.013931 | 0.0025533 | 4.8682e-8 | |
| 4 | 0 | 0 | chr8_20001513_AC_A | 8:20001513_AC_A | 8 | 20001513 | AC | A | 0.059415 | 0.025936 | 0.021973 | |
| 5 | 0 | 0 | chr8_20001995_G_A | rs7003858 | 8 | 20001995 | G | A | -0.0040079 | 0.0024412 | 0.10063 | |
| 6 | 0 | 0 | chr8_20002426_G_A | rs1497020 | 8 | 20002426 | G | A | -0.0041822 | 0.002443 | 0.086919 | |
| 7 | 0 | 0 | chr8_20002739_T_TG | rs34201967 | 8 | 20002739 | T | TG | -0.0038998 | 0.0024475 | 0.11107 | |
| 8 | 0 | 0 | chr8_20002888_A_G | rs17092091 | 8 | 20002888 | A | G | 0.011931 | 0.015152 | 0.43105 | |
| 9 | 0 | 0 | chr8_20024050_G_C | rs376551459 | 8 | 20024050 | G | C | -0.0075333 | 0.029592 | 0.79905 | |
| 10 | 0 | 0 | chr8_20003431_AT_A | rs749194432 | 8 | 20003431 | AT | A | -0.018139 | 0.0025424 | 9.7147e-13 | |

Showing 1 to 10 of 544 entries CSV Excel Previous 1 2 3 4 5 … 55 Next

Download the full table (as .RData file) | Download the full table with updated CredibleSets (as .RData file)

---Table: state---

```
List of 3
 $ impl    :Classes 'ReactiveValues', 'R6' <ReactiveValues>
  Public:
    .allValuesDeps: environment
    .dedupe: TRUE
    .dependents: Map, R6
    .hasRetrieved: list
    .label: reactiveValues1090
    .metadata: Map, R6
    .namesDeps: environment
    .reactId: r505
    .values: Map, R6
    .valuesDeps: environment
    clone: function (deep = FALSE)
    freeze: function (key, invalidate = FALSE)
    get: function (key)
    getMeta: function (key, metaKey)
    initialize: function (dedupe = TRUE, label = paste0("reactiveValues", p_randomInt(1000,
    isFrozen: function (key)
    mset: function (lst)
    names: function ()
    self: ReactiveValues, R6
    set: function (key, value, force = FALSE)
```

### Useful feature: Need help?

The dynamic "Need Help?" magic button in the top-left corner of the *flashfmZoom* interface contains information that helps users to understand the key features of each tab in the main panel - it provides dynamic and unique information based on the users' selected tab. If there is any error or warning message during the analysis, the text of the message will be highlighted in red and displayed under the "Need help?" button.

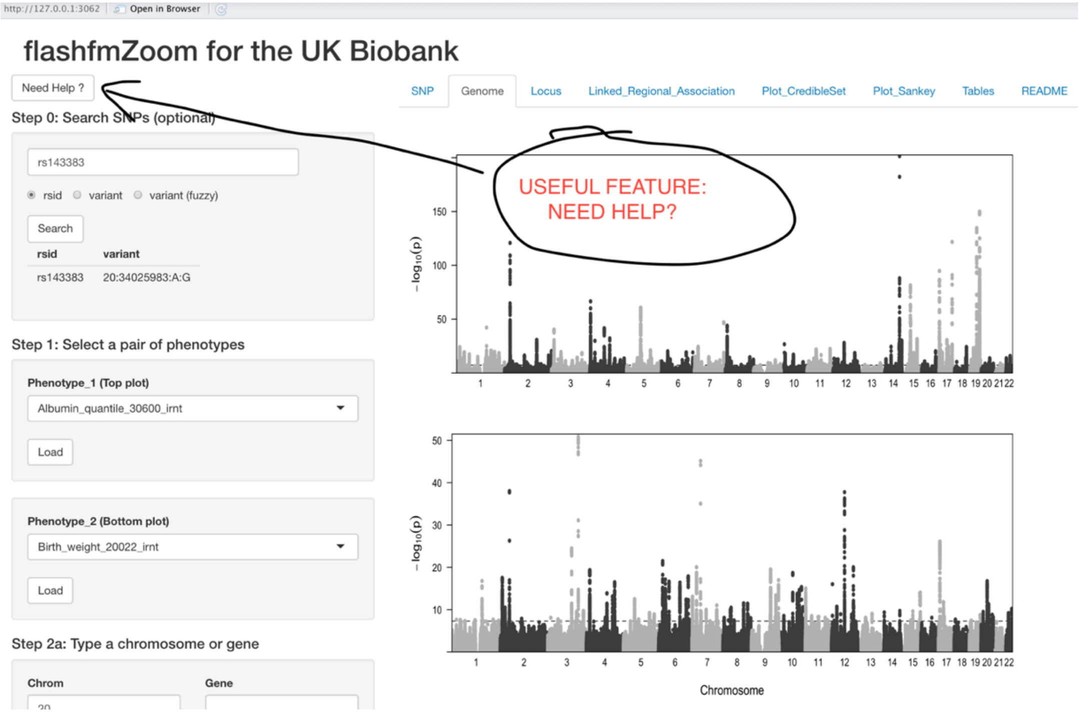


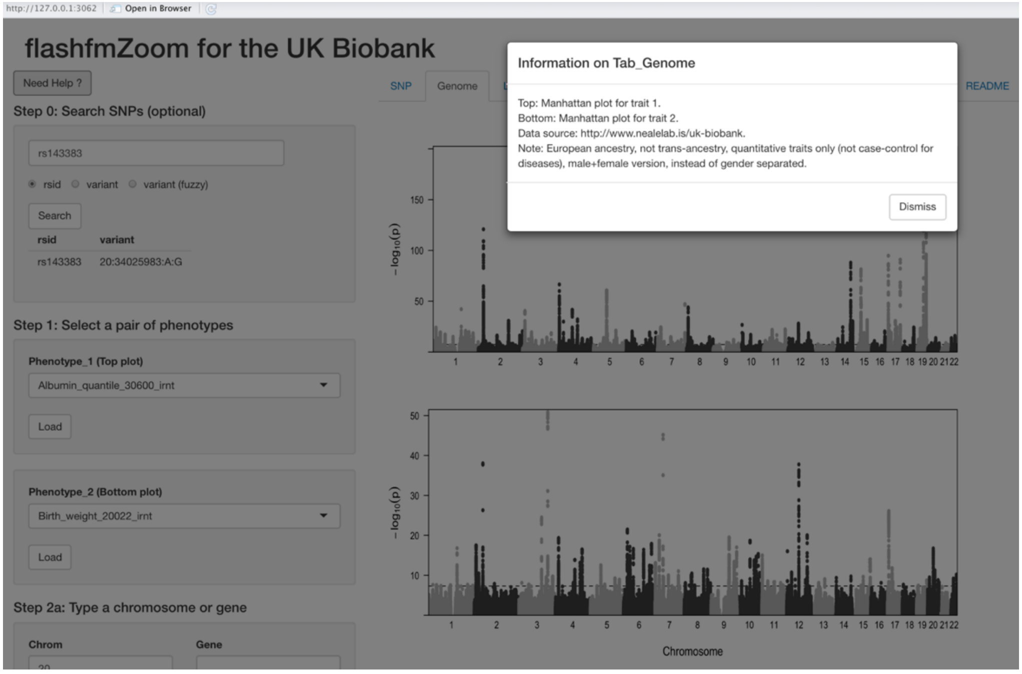

**Table S4: Modebar icons that offer users different interactive features in plots**

| Icon | Use |
|---|---|
| 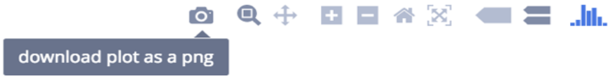  | Click on the camera icon and get/download the plot in PNG format. |
| 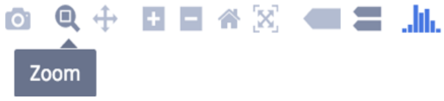  | Clicking this icon selects the Zoom mode. To zoom in on a region of a graph, click and hold your mouse, moving across the region. Release your mouse. To return to the original view, double-click anywhere on the plot. |
| 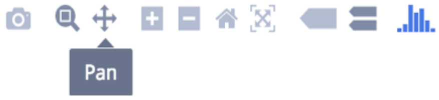  | To pan across regions of your graph, select the Pan mode. Click and hold your mouse to explore the data. Double-click anywhere to return to the original view. |
| 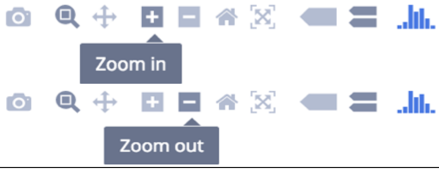  | You can zoom in and out by clicking on the + and - buttons. The plot keeps axes labels and annotations the same size to preserve readability. |
| 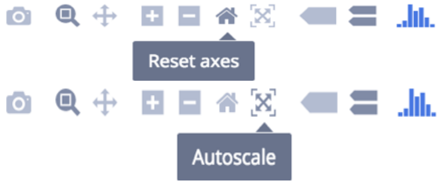  | Clicking this icon zooms to include your Axes Range, if this has been set. If it has not been set it zooms to a setting that is optimized to include all the viewable data, the same as if Autoscale had been clicked. |
| 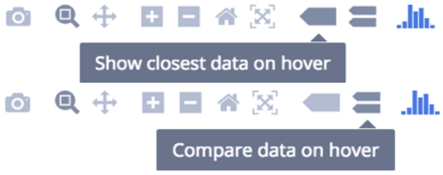  | One of these two buttons is always selected. Clicking 'Show closest data on hover' will display the data for just one point under the cursor. Clicking 'Compare data on hover' will show you the data for all points with the same x-value. |
| 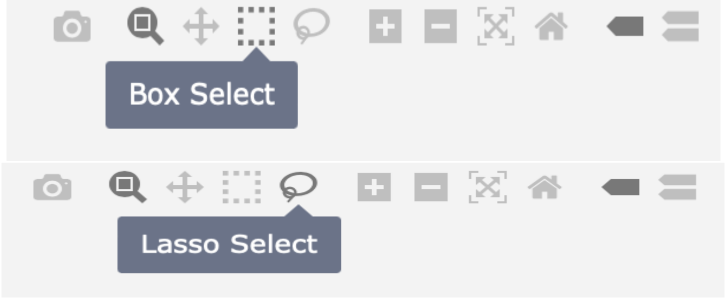  | These are basic selection tools. Box Select is to select data on the plot by creating a box, while the Lasso Select is used to draw with freehand selection. |
| | **In general, by double-clicking anywhere on the plot, users can return to the original view.** |
| **More reference:** | [https://plotly.com/chart-studio-help/getting-to-know-the-plotly-modebar/](https://plotly.com/chart-studio-help/getting-to-know-the-plotly-modebar/) |

### Terminology in the plots and the flashfm package (summary)

The user may also interact with the plots to select different perspectives and download their selected version.

Below we refer to:

- **PP** (posterior probability) for each multi-SNP model: the model PP for each configuration of variants being joint causal variants for the trait (i.e., multiple causal variants). The PPs sum to 1.
- **MPP** (marginal posterior probability of variant causality): the posterior probability that the SNP is included in any causal model; it is the sum over all model PPs that include the SNP. MPP is also known as PIP (Posterior Inclusion Probability).
- **SNP groups:** SNPs belonging to the same group can be viewed as exchangeable. SNPs with MPP>0.001 are assigned to the same group if they are strongly correlated (pairwise $r^2$>0.6) and rarely appear in a model together (marginal posterior probability that a pair of SNPs are both included in a model < 0.01). Each SNP group based on flashfm tends to contain a subset of variants from groups based on single-trait fine-mapping.
- **PPg (PP based on SNP groups):** for a particular group-based model containing multiple SNP groups (e.g. groups A and B), PPg is the sum of PPs for all models composed of exactly one SNP from group A and one SNP from group B. As the SNPs in each group are exchangeable, the PPg represents the probability that the correct causal model involves a SNP from group A and a SNP from group B.
- **MPPg (MPP based on SNP groups):** MPPg for SNP group A is the sum of MPPs for all SNPs that belong to group A; it is the sum over all model PPs that include a SNP from group A. This represents the probability of at least one of the SNPs in SNP group A being a causal variant for the traits.
- **fm** refers to a single-trait fine-mapping method, either JAM (Newcombe et al. 2016) if using the internal flashfm function 'FLASHFMwithJAM' or FINEMAP (http://www.christianbenner.com/) if using the combination of FINEMAP and flashfm.
- **cs1:** a 99% credible set that is constructed from the single-trait fine-mapping (fm) model PPs such that it has a 99% chance of containing all the causal variants. The models are sorted by decreasing PP and the credible set includes all variants that appear in the top models, with PPs that sum to at least 99%. A "1" in the interactive text box for a SNP indicates the SNP is in the fm 99% credible set, and "0" otherwise.
- **csM:** a 99% credible set that is constructed from multi-trait fine-mapping (flashfm) model PPs such that it has a 99% chance of containing all the causal variants. The models are sorted by decreasing PP and the credible set includes all variants that appear in the top models, with PPs that sum to at least 99%. A "1" in the interactive text box for a SNP indicates the SNP is in the flashfm 99% credible set, and "0" otherwise.

### Reference (as included in Supplementary Material)


1. Benner, C., Spencer, C. C., Havulinna, A. S., Salomaa, V., Ripatti, S., & Pirinen, M. (2016). FINEMAP: efficient variable selection using summary data from genome-wide association studies. *Bioinformatics*, 32(10), 1493-1501.
2. Bulik-Sullivan, B., Finucane, H. K., Anttila, V., Gusev, A., Day, F. R., Loh, P. R., ... & Neale, B. M. (2015). An atlas of genetic correlations across human diseases and traits. *Nature genetics*, 47(11), 1236-1241.
3. Hernandez, N., Soenksen, J., Newcombe, P., Sandhu, M., Barroso, I., Wallace, C., Asimit, J.L. (2021). The flashfm approach for fine-mapping multiple quantitative traits. *Nat Commun* 12, 6147.
4. Newcombe, P. J., Conti, D. V. & Richardson, S. (2016). JAM: a scalable Bayesian framework for joint analysis of marginal SNP effects. *Genetic Epidemiology*. 40, 188–201.
